# Single-component multilayered self-assembling protein nanoparticles displaying extracellular domains of matrix protein 2 as a pan-influenza A vaccine

**DOI:** 10.1101/2023.06.02.543464

**Authors:** Keegan Braz Gomes, Yi-Nan Zhang, Yi-Zong Lee, Mor Eldad, Alexander Lim, Garrett Ward, Sarah Auclair, Linling He, Jiang Zhu

## Abstract

The development of a cross-protective pan-influenza A vaccine remains a significant challenge. In this study, we designed and evaluated single-component self-assembling protein nanoparticles (SApNPs) presenting the conserved extracellular domain of matrix protein 2 (M2e) as vaccine candidates against influenza A viruses. The SApNP-based vaccine strategy was first validated for human M2e (hM2e) and then applied to tandem repeats of M2e from human, avian and swine hosts (M2ex3). Vaccination with M2ex3 displayed on SApNPs demonstrated higher survival rates and lower weight loss compared to the soluble M2ex3 antigen against lethal challenges of H1N1 and H3N2 in mice. M2ex3 I3-01v9a SApNPs formulated with a squalene-based adjuvant were retained in the lymph node follicles over eight weeks and induced long-lived germinal center reactions. Notably, a single low dose of M2ex3 I3-01v9a SApNP formulated with a potent adjuvant, either a Toll-like receptor 9 (TLR9) agonist or a stimulator of interferon genes (STING) agonist, conferred 90% protection against a lethal H1N1 challenge in mice. With the ability to induce robust and durable M2e-specific functional antibody and T cell responses, the M2ex3-presenting I3-01v9a SApNP provides a promising pan-influenza A vaccine candidate.

**ONE-SENTENCE SUMMARY:** Single-component self-assembling protein nanoparticles (SApNPs) displaying tandem M2e elicit robust and durable immunity that may protect against influenza A viruses of diverse origins.

## INTRODUCTION

Influenza (flu) is a respiratory disease caused by influenza viruses of the Orthomyxoviridae family (*1–5*). Influenza viruses are enveloped negative-sense, single-stranded RNA viruses (*6*) that can be classified as type A, B, C, or D, with influenza A and B viruses (IAVs and IBVs) posing a major threat to human health. The most abundant surface glycoprotein, hemagglutinin (HA), binds to sialic acid receptors on host cells to facilitate cell entry (*7, 8*). Under the host’s immune selection pressure, HA can acquire amino acid substitutions that lead to escape mutants (*8*). Another surface glycoprotein, neuraminidase (NA), aids in the release of viral particles via cleavage of residues on the host cell’s surface (*9, 10*). Matrix protein 1 (M1) is involved in virus budding, while matrix protein 2 (M2) functions as a proton channel to facilitate the maintenance of pH during viral entry and replication in host cells (*11*). IAVs can be classified into subtypes based on the antigenic properties of HA and NA (*4*), with H1N1 and H3N2 being responsible for most human infections (*8*). IBVs have a single HA/NA subtype, which can be classified into two lineages, Victoria and Yamagata (*1, 2*). IAVs can infect many hosts, whereas IBVs are restricted to humans (*12*).

Seasonal flu vaccines have been used as a cost-effective public health tool since the 1940s (*13–15*). Current flu vaccines are typically quadrivalent, covering two IAV subtypes (H1N1 and H3N2) and two IBV lineages (Victoria and Yamagata), and are produced in chicken eggs (*16*). As a result, current flu vaccines mainly generate strain-specific neutralizing antibodies (NAbs) and may not protect against mismatched seasonal strains or more distinct strains generated through “antigenic drift,” in which HA and NA accumulate small mutations over time. Occasionally, IAVs have the potential to cause global pandemics through “antigenic shift,” in which HAs and NAs from different host species recombine to form novel IAV strains against which the human population lacks pre-existing immunity (*17*). Viral reassortment resulting in highly pathogenic avian influenza (HPAI) acquiring animal-to-human transmissibility has been on the rise in recent years. As of 2021, there have been 863 cases of HPAI H5N1 and 66 cases of H5N6 in humans, with a > 50% fatality rate (*18*). Therefore, there is an urgent need for cross-protective flu vaccines (*16*), especially for potential pandemic strains originating from diverse animal reservoirs (*19*).

Various antigen and vaccine strategies have been explored to develop a universal influenza vaccine (*20–28*). One strategy targets conserved internal proteins, such as nucleoprotein and M1, to induce influenza-specific T cell responses (*29*). A second strategy aims to generate broadly neutralizing antibodies (bNAbs) to the conserved regions of HA, such as the stem and parts within the head domain (*30–34*), and of NA (*35*). Notably, the highly conserved ectodomain of the M2 protein (M2e) presents an attractive target for universal IAV vaccine development (*31, 36–39*) because of the sequence conservation across IAVs and functional importance of the M2 proton channel to virus fitness and replication. Although M2e is small (∼23 aa) and poorly immunogenic, it can be conjugated to large molecular carriers to elicit antibody responses that effectively reduce viral replication (*40*). Unlike HA and NA, M2e-specific antibodies protect via FcγR-dependent mechanisms, such as antibody-dependent cellular cytotoxicity (ADCC) and phagocytosis (ADCP), rather than direct neutralization (*39, 41–43*). Various carriers have been used to increase the immunogenicity of M2e vaccines, including hepatitis B core protein (HBc) (*41*), tobacco mosaic virus (TMV) coat protein (*44*), keyhole limpet hemocyanin (KLH) (*40*), rotavirus NSP4 (*45*), GCN4 (*46*), bacterial flagellin (*47*), liposomes (*48*), polymers (*49–51*), and gold nanoparticles (NPs) (*52, 53*). Early human trials confirmed the immunogenicity and tolerance of M2e vaccines but also revealed several weaknesses. For example, an adjuvanted M2e-HBc fusion vaccine (*41*) induced a short-lived anti-M2e antibody response, and an M2e-flagellin fusion vaccine (*47*) caused undesirable side effects at higher doses. A vaccine combining M2e with cytotoxic T lymphocyte (CTL) epitopes (*54*) induced strong cellular immunity, but this response was narrow and slow, making it unsuitable for effectively mitigating a future influenza pandemic. These clinical trials highlight the challenges facing M2e-based vaccine development (*37, 55*), as well as the importance of vaccine carriers, adjuvants, balanced antibody and T cell responses, and durability.

In recent studies, we have demonstrated a rational vaccine strategy that combines antigen optimization and protein NP display (*56–61*). This strategy is inspired by the success of virus-like particles (VLPs), which have been used as vaccines against cognate viruses or as carriers for foreign antigens (*62–69*). Due to their inherent complexity, it remains a challenge to produce VLPs with yield, purity, and quality acceptable for clinical use. As an alternative, protein NPs can be constructed to mimic VLPs, with the capability of displaying diverse antigens (*70, 71*). We have previously designed single-component self-assembling protein nanoparticles (SApNPs) based on two bacterial proteins – E2p from *Bacillus stearothermophilus* (*72*) and I3-01 from *Thermotoga maritima* (*73*), which form 60-mers of 22-25 nm in diameter (*56–61*). Genetic fusion of an antigen to the N-terminus of an SApNP subunit and a locking domain (LD) and pan-reactive T cell epitope (PADRE) to the C-terminus creates a “vaccine construct” encoding a single polypeptide (hence single-component), which assembles with identical polypeptides into a multilayered NP structure with an array of antigens on the surface, a stabilizing inner LD layer, and a hydrophobic PADRE core. The incorporation of LD and PADRE increased the yield, purity, and stability of resulting SApNPs (*56–58*), highlighting the beneficial effects of this multilayered NP design. These SApNPs can be readily expressed in Chinese hamster ovary (CHO) cells and have been successfully applied to vaccine development for HIV-1 (*56, 60, 61*), HCV (*59*), Ebola virus (EBOV) (*58*), and SARS-CoV-2 (*57, 74*). Notably, the multilayered SApNPs can be retained in lymph nodes for weeks, enabling them to interact with immune cells and generate robust germinal center (GC) reactions, whereas individual soluble antigens are cleared within a few hours (*56, 74*).

In this study, we rationally designed M2e-presenting SApNPs as cross-protective, pan-influenza A vaccine candidates and characterized them both in vitro and in vivo. We first designed I3-01v9a, a new version of the I3-01v9 SApNP, for optimal presentation of monomeric antigens. We then displayed human M2e (hM2e) on ferritin (FR), E2p, and I3-01v9a SApNPs, along with a trimeric scaffold, for initial assessment. Following detailed in vitro characterization, these hM2e immunogens were tested in mice, which were sequentially challenged with mouse-adapted H1N1 and H3N2 after a two-dose vaccination. Based on the results, we next displayed tandem copies of M2e (human, avian and swine), termed M2ex3, on the same carriers and characterized these immunogens following a similar protocol. M2ex3 presented on SApNPs elicited significantly higher M2e-specific antibody and T cell responses in immunized mice compared with the soluble M2ex3 antigen. As a result, mice immunized with M2ex3 SApNPs showed higher survival rates against lethal heterosubtypic challenges. In the mechanistic analysis, M2ex3 SApNPs exhibited prolonged retention (8 weeks) in lymph node follicles and robust GC reactions, which may explain the vaccine-induced immunity and protection. Lastly, a single low-dose immunization of M2ex3 I3-01v9a SApNP formulated with a potent adjuvant identified in our previous study (*74*), either a Toll-like receptor 9 (TLR9) agonist or a stimulator of interferon genes (STING) agonist, conferred 90% protection against a lethal H1N1 challenge in mice. Therefore, the tandem M2e presented on an optimized I3-01v9a SApNP may provide an effective vaccine candidate for durable protection against seasonal and pandemic influenza A viruses.

## RESULTS

### Rational design of an I3-01v9a NP scaffold for presenting monomeric antigens

In our early studies, we utilized 24-mer ferritin (FR) and two 60-mers, E2p and I3-01, to display HIV-1 and HCV antigens (*56, 59*). Locking domains (LDs) and a CD4 T-helper epitope (PADRE) were later incorporated into E2p and I3-01 (and its variant I3-01v9) to generate “multilayered” NP carriers, which were successfully used to display stabilized EBOV glycoprotein (GP) trimers (*58*), SARS-CoV-1/2 spikes (*57, 74*), and HIV-1 envelope (Env) trimers (*56*) for vaccine development. Notably, the I3-01 NP scaffold appeared to be particularly amendable to structural modification, with multiple design variants (e.g., I3-01v9) tested in our previous studies (*56–60*). In this study, we rationally optimized the I3-01v9 NP scaffold to achieve the optimal surface display of various M2e antigens (**Figure S1**). The N-termini of I3-01v9 forms a wide triangle of 50.5 Å, making it more suitable than E2p for displaying monomeric antigens. However, the first amino acid (the antigen anchoring site) is below the NP surface, and as a result, a long flexible peptide linker must be used to connect the antigen to the I3-01v9 N-terminus, leading to the increased structural instability of the antigen-NP fusion constructs. Here, we hypothesized that extending the I3-01v9 N-terminal helix would allow its first residue to reach the NP surface, and consequently, a short peptide linker between the antigen and NP backbone would be sufficient. A computational procedure was devised to facilitate rational design (**Figure S1**). Briefly, the backbone of a helix (residues 953-982) from a c-MYC transcription factor protein (PDB ID: 6G6L) was grafted onto an I3-01v9 subunit by using residues Glu2 and Glu3 of I3-01v9 for structural fitting. The extended N-terminal helix was then truncated to 11 residues so that its first residue would be just above the NP surface. Then, a protein structure sampling program, CONCOORD (*75*), was used to generate 1,000 slightly perturbed conformations for the modified I3-01v9 subunit. Next, an ensemble-based protein design program that was previously used to optimize HIV-1 Env trimers (*56*) and HCV E2 cores (*59*) was employed to predict the sequence for the first 9 residues of the 11-residue segment using C_α_ and C_β_-based RAPDF scoring functions (*76*). The final design, termed I3-01v9a, was determined by combining results from predictions using both scoring functions (**Figure S1**).

### Human M2e (hM2e) on multilayered SApNPs as human influenza A vaccines

The M2 protein from IAVs is a highly conserved proton channel, with a small ectodomain of 24 amino acids in length (M2e) (*37*). Anti-human M2e (hM2e) antibodies have been shown to reduce viral replication, thus decreasing clinical symptoms and severity of the disease. In the immunogen design, M2e is hereafter defined as residues 2-24, excluding the first methionine.

Previously, we rationally redesigned viral antigens and engineered antigen-presenting SApNPs for in vitro and in vivo characterization (*56–61*). Following a similar strategy, we designed an hM2e scaffold and three SApNP constructs. The crystal structures of hM2e in complex with antibodies mAb65 and mAb148 (*77, 78*) indicate that hM2e is flexible and can adopt different conformations upon antibody binding. MAb65 recognizes a short turn of Pro10 to Asn13, whereas mAb148 binds to an N-terminal epitope (Ser2-Glu8). We first utilized a capsid-stabilizing protein of lambdoid phage 21, SHP (PDB ID: 1TD0), as a trimeric scaffold to present hM2e. With a 5GS linker, two hM2e epitopes would span ∼9.1 nm (measured at Pro10) when all three 1TD0-attached hM2e segments were in a fully open conformation (**Figure 1A**). We then displayed hM2e on FR 24-mer and reengineered E2p and I3-01v9a 60-mers, all with a 5GS linker (**Figure 1A**). Molecular modeling revealed well-spaced hM2e peptides on the particle surface, with diameters of 20.9 nm, 29.1 nm, and 32.4 nm for FR, E2p, and I3-01v9a, respectively (**Figure 1A**). Following a similar terminology, the “multilayered” E2p and I3-01v9a are named E2p-LD4-PADRE (E2p-L4P or simply E2p) and I3-01v9a-LD7-PADRE (I3-01v9a-L7P or simply I3-01v9a), respectively. One hM2e scaffold and three hM2e-presenting SApNPs were subjected to in vitro characterization.

**Figure 1.**
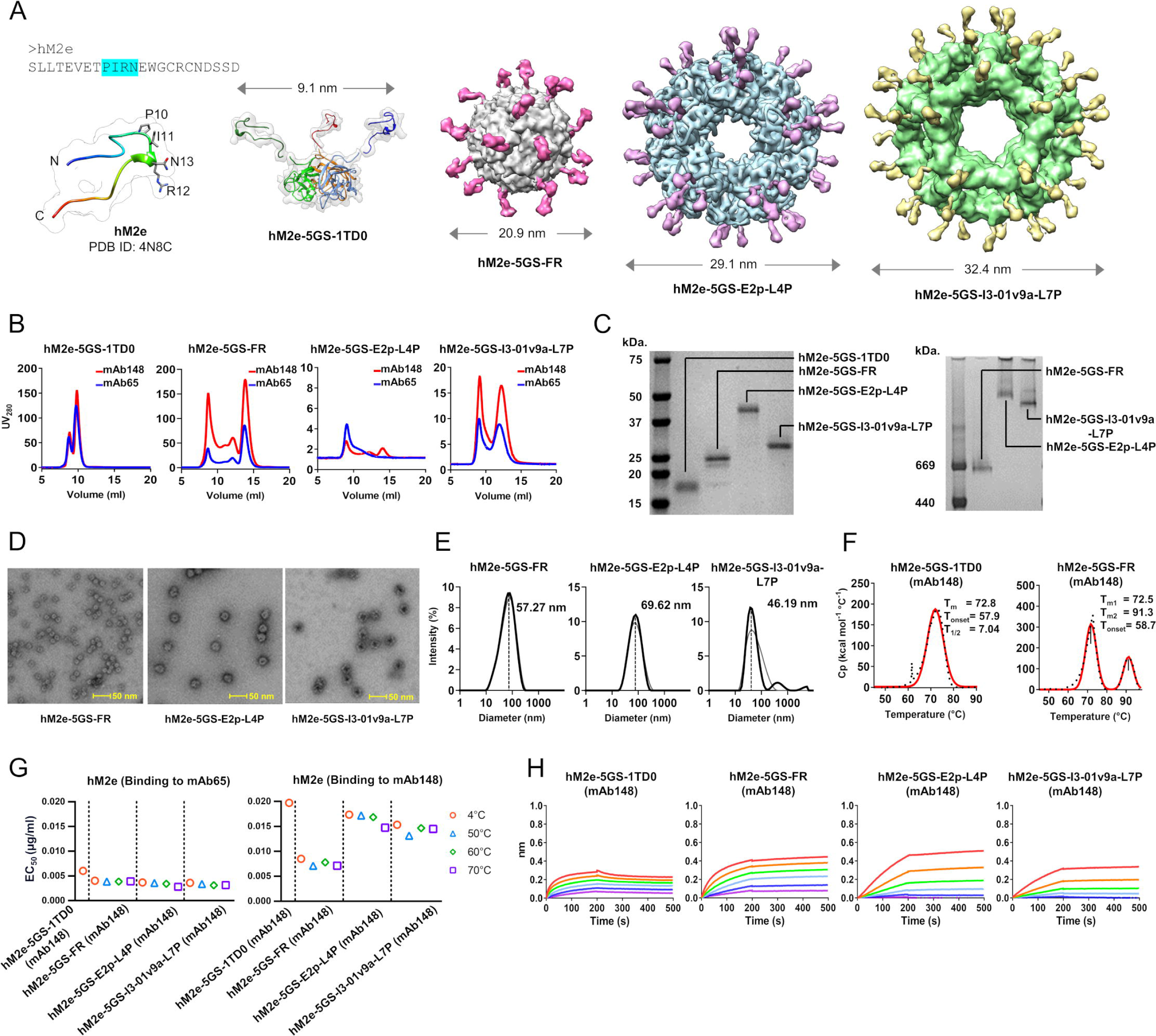
Design and in vitro characterization of hM2e immunogens. (**A**) Structural models of human M2e (hM2e), hM2e-5GS-1TD0 trimer, and three hM2e-presenting nanoparticles (NPs). Left: Amino acid sequence and ribbons/surface model of hM2e (from PDB ID 4N8C). Middle: Ribbons/surface model of hM2e-5GS-1TD0 trimer, in which a trimeric viral capsid protein SHP (PDB ID: 1TD0) is used to display hM2e. Right: Surface models of hM2e on 24-meric ferritin (FR) and 60-meric E2p-L4P and I3-01-L7P SApNPs. The SApNP size is indicated by diameter (in nm). (**B**) SEC profiles of hM2e-5GS-1TD0 trimer and hM2e-presenting FR, E2p-L4P and I3-01-L7P SApNPs. The hM2e trimer was processed on a Superdex 75 10/300 increase GL column, while three SApNPs were processed on a Superose 6 increase 10/300 GL column. (**C**) SDS-PAGE under reducing conditions (left) and BN-PAGE (*34*) of hM2e-presenting FR, E2p-L4P and I3-01v9a-L7P SApNPs. Notably, hM2e-5GS-1TD0 is included on the SDS gel for comparison. (**D**) Negative-stain EM micrographs of mAb148-purified FR, E2p-L4P and I3-01v9a-L7P SApNPs. (**E**) DLS profiles of mAb148-purified FR, E2p-L4P and I3-01v9a-L7P SApNPs. Average particle size derived from DLS are labeled. (**F**) Thermostability of the hM2e-5GS-1TD0 trimer and hM2e-5GS-FR SApNP with *T*m, Δ*T*_1/2_ and *T*_on_ measured by DSC. (**G**) ELISA analysis of the hM2e trimer and SApNPs (FR, E2p-L4P and I3-01v9a-L7P) binding to mAb65 (left) and mAb148 (*34*) after heating to 50 °C, 60 °C and 70 °C for 15 minutes. (**H**) Antigenic profiles of the hM2e trimer and SApNPs (FR, E2p-L4P and I3-01v9a-L7P) to mAb148 using BLI.

All four hM2e constructs (**Figure S2A**) were transiently expressed in 25 ml ExpiCHO cells, purified by immunoaffinity chromatography (IAC) (*79*) using mAb65 or mAb148 columns, and analyzed by size exclusion chromatography (SEC) (**Figure 1B**). The SEC profiles indicated high yield for hM2e-5GS-1TD0 and hM2e-5GS-FR and, in contrast, a notably lower yield for hM2e-5GS-I3-01v9a-L7P, as shown by the ultraviolet absorbance at 280 nm (UV_280_). Among the four constructs, hM2e-5GS-E2p-L4P had the lowest yield. Of note, all three SApNPs showed two SEC peaks at 8-9 ml and 13-14 ml, corresponding to different NP species. Sodium dodecyl sulfate-polyacrylamide gel electrophoresis (SDS-PAGE) under reducing conditions showed bands for hM2e-5GS-1TD0 (13.9 kDa), FR (21.0 kDa), E2p-L4P (38.9 kDa) and I3-01v9a-L7P (33.3 kDa) that were consistent with their calculated molecular weights (MW) (**Figure 1C**, left). Blue native-polyacrylamide gel electrophoresis (BN-PAGE) confirmed the high purity of SApNP samples after IAC using an mAb148 column, displaying a single high-MW band for each SApNP with no sign of unassembled species (**Figure 1C**, right). The structural integrity of IAC-purified SApNPs was validated by negative-stain electron microscopy (nsEM), which showed distinct morphologies for three hM2e-presenting SApNPs (**Figure 1D**). Notably, hM2e SApNPs appeared to form “clusters” in solution, which likely correspond to the high-MW peak (8-9 ml) in their SEC profiles (**Figure 1B**). Analysis of mAb148-purified SApNPs by dynamic light scattering (DLS) revealed larger-than-expected “particle” size for hM2e FR (57.2 nm), E2p-L4p (69.6 nm), and I3-01v9a-L7P (46.1 nm) (**Figure 1E**), consistent with the nsEM results. Interestingly, DLS analysis of the SEC fraction (13-14 ml) of an mAb148-purified hM2e-5GS-FR sample still showed three particle size populations, suggesting that cluster formation is an intrinsic feature of hM2e SApNPs (**Figure S2B**). Differential screening calorimetry (DSC) (*41*) was used to quantify the thermostability of these hM2e constructs. Thermograms were obtained for hM2e-5GS-1TD0 and hM2e-5GS-FR, which showed a melting temperature (T_m_) of 72.5-72.8°C and a similar T_onset_ of 57.9-58.7°C (**Figure 1F**). For the two large 60-mers, heating, enzyme-linked immunosorbent assay (ELISA), and nsEM were combined to estimate thermostability. Briefly, SApNP samples were heated to 50°C, 60°C, and 70°C for 15 min prior to ELISA analysis against mAb148 and mAb65 (**Figure 1G**, **Figure S2C**) and nsEM (**Figure S2D**). While antibody binding, measured by half maximal effective concentration (EC_50_), remained largely consistent within a temperature range of 4-70°C, nsEM images showed signs of irregular particle shapes at 70°C, suggesting that the melting points for hM2e-5GS-E2p-L4P and I3-01v9a-L7P may be between 60 and 70°C. Lastly, we performed bio-layer interferometry (BLI) to quantify antibody binding kinetics for the four hM2e constructs. Although the three SApNPs outperformed the trimeric hM2e scaffold regardless of the antibody tested (**Figure 1H**, **Figure S2E**), mAb148 and mAb65 showed different profiles, with stronger binding observed between mAb65 and the two large 60-meric SApNPs (**Figure S2E**).

In summary, hM2e can be successfully displayed on all three SApNPs, consistent with our previous studies where stabilized HIV-1 (*56, 60, 61*), HCV (*59*), EBOV (*58*), and SARS-CoV-1/2 (*57, 74*) antigens were displayed on the surface of SApNPs. Extensive biochemical, biophysical, structural, and antigenic characterizations provided detailed in vitro profiles of hM2e-presenting SApNPs, thus allowing evaluation of these vaccine immunogens in vivo.

### In vivo evaluation of a scaffolded hM2e trimer and hM2e-presenting SApNPs in mice

The immunogenicity and protective efficacy of hM2e trimer and hM2e-presenting SApNPs were evaluated in BALB/c mice. First, mouse-adapted A/Puerto Rico/8/1934 (PR8) H1N1 and A/Hong Kong/1/1968 (HK/68) H3N2 were grown in Madin-Darby canine kidney (MDCK) cells, and the propagated viruses at various dilutions were used to challenge mice to establish a 50% lethal dose in mice (LD_50_) (**Figure 2A**). Plaque-forming units (PFU) of the virus were measured via plaque assay. Using survival rates of mice at various viral dilutions, the Reed-Muench and Spearman-Karber methods (*80, 81*) were used to calculate the 50% endpoint titers for survival. For a stock of 3.8 × 10^5^ PFU/ml PR8 (H1N1), LD_50_ was determined to be 12 PFU/ml. For a stock of 5.5 × 10^4^ PFU/mL of HK/68 (H3N2), LD_50_ was determined to be 1.2 × 10^4^ PFU/ml.

**Figure 2.**
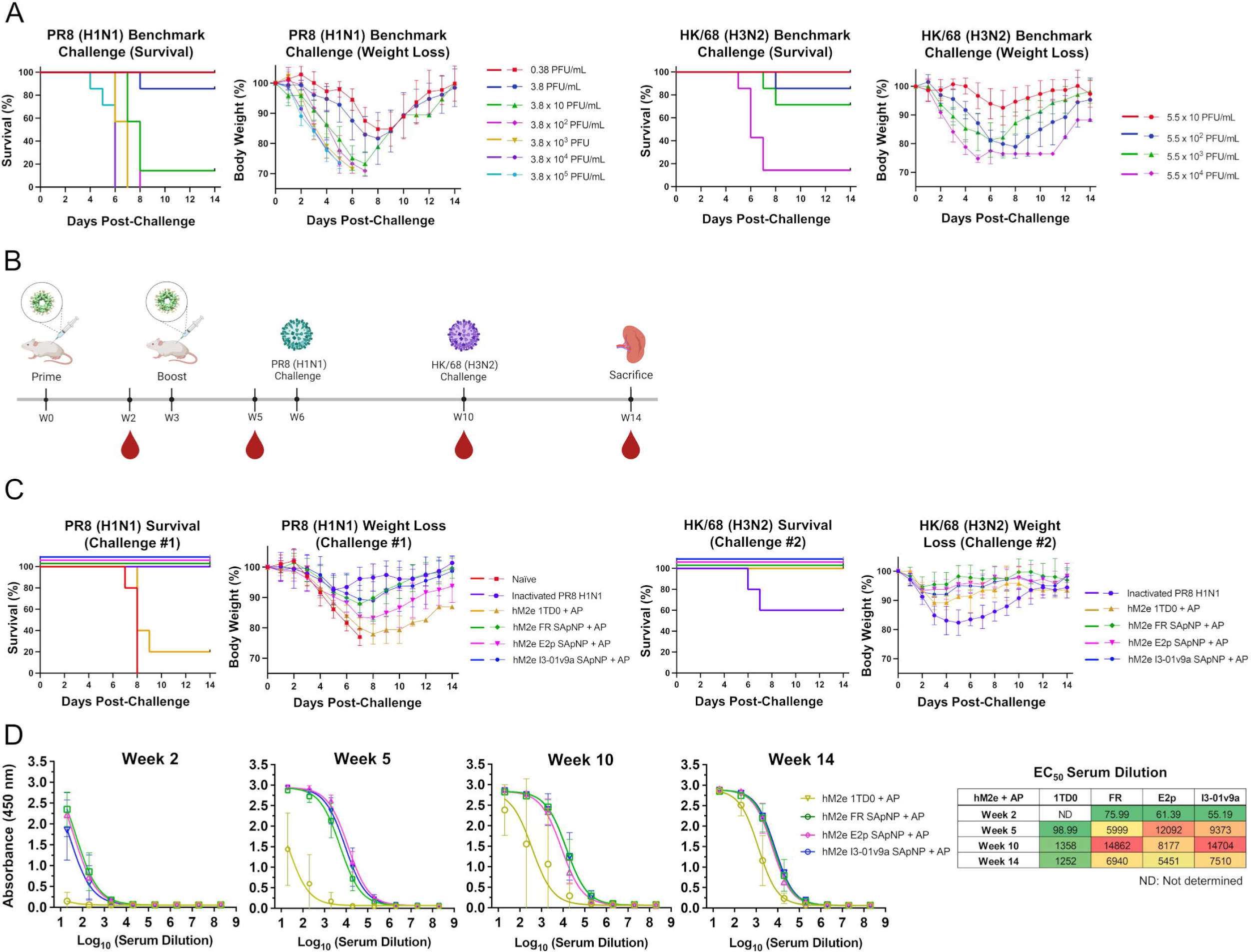
Assessment of hM2e scaffolds and nanoparticles in a mouse challenge model. (**A**) Benchmark challenge studies assessing survival and weight loss to establish the 50% lethal intranasal challenge dose in mice for mouse-adapted A/Puerto Rico/8/1934 (PR8) H1N1 and A/Hong Kong/1/1968 (HK/68) H3N2; N = 7 mice/group. Mice were monitored for survival, weight loss, and morbidities for 14 days. (**B**) Schematic representation of mouse immunization regimen for hM2e constructs, sequential challenges of LD_50_ × 10 of PR8 (H1N1) and HK/68 (H3N2), blood collection, and sacrifice; N = 10 mice/group. (**C**) Survival and weight loss of mice challenged with LD_50_ × 10 of PR8 (H1N1) followed by an LD_50_ × 10 of HK/68 (H3N2). Mice were monitored for survival, weight loss, and morbidities for 14 days. (**D**) ELISA curves showing hM2e-immune sera binding to the hM2e-5GS-foldon trimer probe and calculated 50% effective concentration (EC_50_) values for weeks 2, 5, 10, and 14. The assay was performed in duplicate with a starting serum dilution of 20× followed by seven 10-fold titrations. Images of mouse immunization, virus challenge, and blood and organ collection created with BioRender.com.

BALB/c mice were immunized via intradermal (ID) injection of hM2e vaccines adjuvanted with aluminum phosphate (AP) (2.5 µg/footpad, 10 µg total) at weeks 0 and 3. Immunized mice were challenged with LD_50_ × 10 of PR8 (H1N1) at week 6; surviving mice were then challenged with LD_50_ × 10 of HK/68 (H3N2) at week 10 (**Figure 2B**). Survival and weight loss were measured for 14 days after each challenge (**Figure 2C**). Following the PR8 (H1N1) challenge, all naïve mice succumbed to the challenge by day 8. Only 20% of mice survived the challenge in the hM2e-5GS-1TD0 (trimer) group, whereas 100% of mice survived in the hM2e FR, E2p, and I3-01v9a SApNP groups. Notably, all mice in the strain-matched inactivated PR8 (H1N1) group (positive control) survived the challenge. Similar trends were observed in the average peak weight loss. While naïve mice suffered the most weight loss (22.1 ± 1.3%), 1TD0 mice lost 19.1 ± 3.3% of their total body weight on average. Among the hM2e SApNP groups, FR mice lost 12.1 ± 8.4% of their total body weight on average, E2p mice lost 16.4 ± 3.9%, and I3-01v9a mice lost the least weight with an average of 10.6 ± 4.5%. In the positive control group, mice immunized with inactivated PR8 (H1N1) lost the least weight upon the strain-matched challenge, with an average loss of 6.5 ± 3.8%. Upon the second challenge with HK/68 (H3N2), the lowest survival rate was observed for the inactivated PR8 (H1N1) group, with only 56% of mice surviving the heterologous challenge. The two surviving 1TD0 mice from the previous challenge and all SApNP mice survived the HK/68 (H3N2) challenge. Following a similar trend, the highest body weight loss was observed for the inactivated PR8 (H1N1) group with an average peak weight loss of 16.9 ± 4.9%. All hM2e vaccine groups showed lower peak weight loss with 10.8 ± 2.4%, 4.4 ± 4.0%, 6.2 ± 3.4%, and 7.9 ± 2.8% for hM2e 1TD0, FR, E2p, and I3-01v9a, respectively. The hM2e-binding antibody response in mouse serum was assessed by ELISA using an hM2e-5GS-foldon trimer (**Figure 2D**, **Figure S3A**). The hM2e SApNP groups demonstrated superior serum binding, measured by EC_50_ titers, at all the time points tested, with the highest fold increase observed at week 5 for hM2e FR (60.6), E2p (122.2), and I3-01v9a (94.7) compared to the 1TD0 group (**Figure 2D**). The hM2e 1TD0, FR SApNP, and I3-01v9a SApNP showed the highest EC_50_ titers 4 weeks after the H1N1 challenge at week 10, whereas for hM2e E2p SApNP the EC_50_ titers peaked at week 5.

In summary, hM2e SApNPs significantly outperformed the soluble hM2e trimer in the vaccination/viral challenge experiment, showing a higher survival rate and lower weight loss that were well-correlated with the heightened M2e-specific serum antibody titers. These hM2e SApNPs also demonstrated cross-protection against H1N1 and H3N2 challenges, whereas the inactivated PR8 (H1N1) vaccine only protected against the strain-matched challenge.

### Tandem M2e (M2ex3) on multilayered SApNPs as pan-influenza A vaccines

The effectiveness of seasonal influenza vaccines range between 10 and 60% as estimated by the U.S. Flu Vaccine Effectiveness Network (*15*). In addition to antigenic drift in circulating human influenza virus strains, the unanticipated emergence of novel strains from swine and avian hosts often causes outbreaks with increased mortality and morbidity. Thus, a broadly protective M2e-based influenza vaccine strategy must incorporate M2e from diverse species.

Following the hM2e vaccine strategy, we designed a trimeric scaffold and three SApNPs to present tandem M2e repeats as vaccine immunogens. Briefly, the hM2e, avian/swine M2e, and human/swine M2e sequences were linked in tandem with short G4 linkers. A structural model of M2ex3 was generated from the crystal structure of hM2e in complex with mAb65 (PDB ID: 4N8C) (**Figure 3A**, left). Although this compact structural model may not represent M2ex3 conformations in solution, it facilitated the rational design of the M2ex3 orientation on various carrier scaffolds. For the M2ex3-5GS-1TD0 trimer, the two outmost hM2e epitopes would span ∼9.4 nm (measured at Pro10 of hM2e) when the scaffolded M2ex3 segments adopt an extended conformation (**Figure 3A**, middle). For the three SApNPs, molecular modeling yielded diameters of 23.2 nm, 32.4 nm, and 36.2 nm for M2ex3-5GS-FR, E2p-L4P, and I3-01v9a-L7P, respectively, measured at Pro10 of hM2e (**Figure 3A**, right three). The display of tandem M2ex3 increased not only the particle size but also the number of M2e epitopes, from 60 to 180, for enhanced immune recognition.

**Figure 3.**
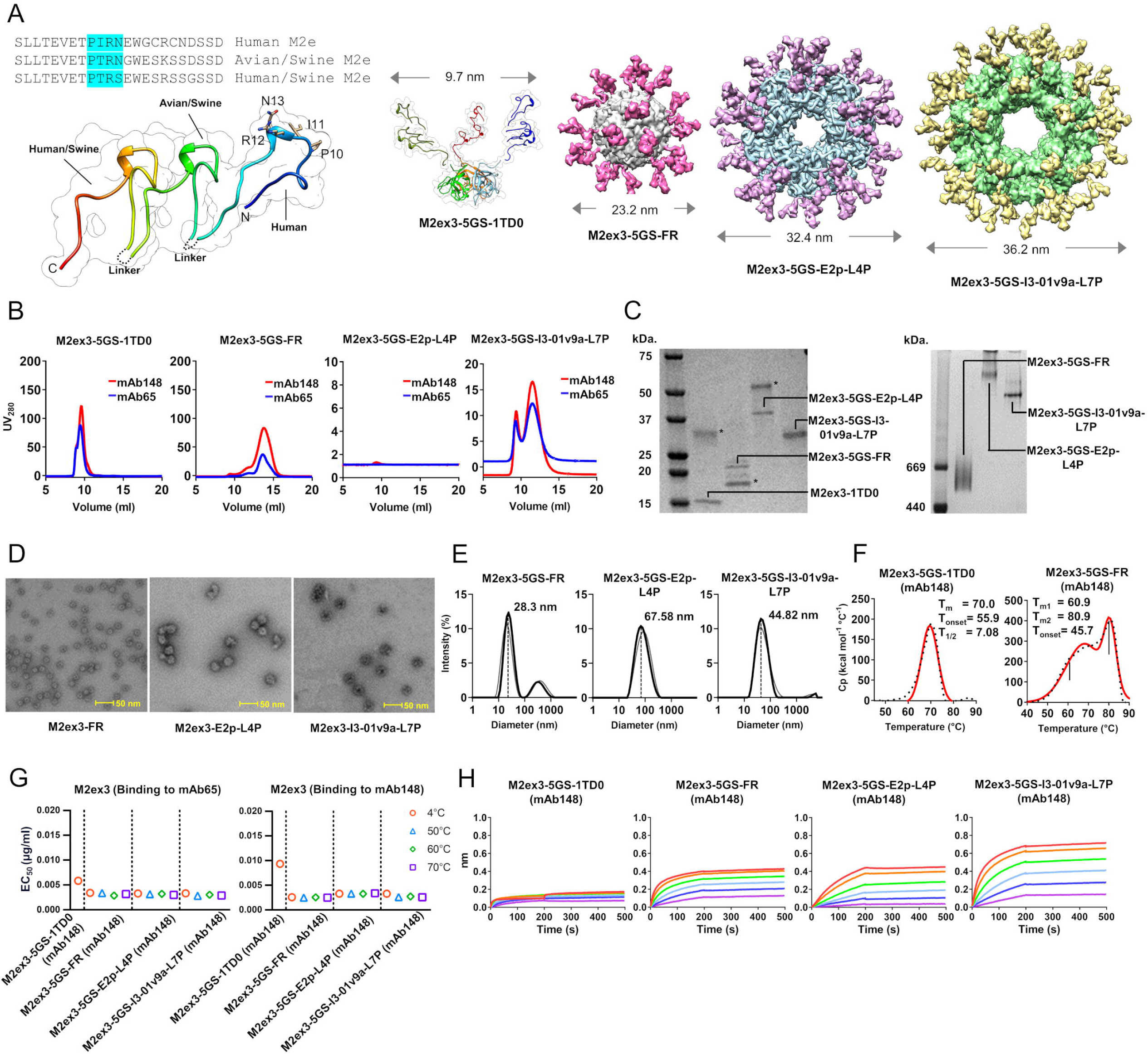
Design and characterization of tandem M2ex3 immunogens. (**A**) Structural models of tandem M2ex3, M2ex3-5GS-1TD0 trimer, and three M2ex3-presenting SApNPs. Left: Amino acid sequences of human, avian/swine, and human/swine M2e and ribbons/surface model of tandem M2ex3 (based on hM2e from PDB ID 4N8C). The G4 linker is shown as a dotted line. Middle: Ribbons/surface model of M2ex3-5GS-1TD0 trimer, in which 1TD0 is a trimeric viral capsid protein. Right: Surface models of M2ex3 on 24-meric ferritin (FR) and 60-meric E2p-L4P and I3-01-L7P SApNPs. The SApNP size is indicated by diameter (in nm). (**B**) SEC profiles of M2ex3-5GS-1TD0 trimer and M2ex3-presenting FR, E2p-L4P and I3-01-L7P SApNPs. The tandem M2ex3 trimer and three SApNPs were processed on a Superdex 75 10/300 increase GL column and a Superose 6 increase 10/300 GL column, respectively. (**C**) SDS-PAGE (left) under reducing conditions and BN-PAGE (*34*) of tandem M2ex3-presenting FR, E2p-L4P and I3-01v9a-L7P SApNPs. Notably, M2ex3-5GS-1TD0 is included on the SDS gel for comparison. (**D**) Negative-stain EM micrographs of mAb148-purified FR, E2p-L4P and I3-01v9a-L7P SApNPs. (**E**) DLS profiles of mAb148-purified FR, E2p-L4P and I3-01v9a-L7P SApNPs. Average particle size derived from DLS are labeled. (**F**) Thermostability of the M2ex3-5GS-1TD0 trimer and M2ex3-5GS-FR SApNP with *T*_m_, Δ*T*_1/2_ and *T*_on_ measured by DSC. (**G**) ELISA analysis of the M2ex3 trimer and SApNPs (FR, E2p-L4P and I3-01v9a-L7P) binding to mAb65 (left) and mAb148 (*34*) after heating to 50 °C, 60 °C and 70 °C for 15 minutes. (**H**) Antigenic profiles of the M2ex3 trimer and SApNPs (FR, E2p-L4P and I3-01v9a-L7P) to mAb148 using BLI.

The four M2ex3 constructs (**Figure S4A**) were expressed and purified using the same strategy as for their hM2e counterparts (**Figure 3B**). Overall, these M2ex3 immunogens showed a similar pattern of expression yield in ExpiCHO cells, with the ranking of 1TD0 > FR > I3-01v9a-L7P >> E2p-L4P. Of note, the SEC profiles showed a less pronounced peak at 9 ml for FR and I3-01v9a-L7P, suggesting a reduced tendency to form NP clusters. Reducing SDS-PAGE showed bands on the gel consistent with MW calculated for M2ex3-5GS-1TD0 (19.3 kDa), FR (26.3 kDa), E2p-L4P (44.4 kDa), and I3-01v9a-L7P (38.7 kDa) (**Figure 3C**, left). However, a second band was observed on the gel for M2ex3-5GS-1TD0, FR, and E2p-L4P under reducing conditions. While the extra bands for M2ex3-5GS-1TD0 and E2p-L4P may indicate higher-MW species that are resistant to the reducing agents, the lower band noted for M2ex3-5GS-FR likely suggests degradation during processing. Nonetheless, BN-PAGE confirmed particle assembly and purity for the three M2ex3 SApNPs **(Figure 3C**, right). Similarly, nsEM micrographs demonstrated well-formed, homogeneous particles for all three M2ex3 SApNP samples following mAb148 purification (**Figure 3D**). In the DLS profiles (**Figure 3E**), M2ex3-5GS-FR exhibited a two-peak distribution with the majority peak showing single particles, indicated by an average size of 28.3 nm. Similarly, M2ex3-5GS-I3-01v9a-L7P yielded a homogenous distribution consistent with single particles. In contrast, M2ex3-5GS-E2p-L4P formed clusters, as seen in nsEM images and indicated by the DLS-derived particle size distribution. The tandem design appeared to cause a reduction in thermostability for M2ex3-5GS-FR, with lower T_m_ (60.9°C) and T_onset_ (45.7°C) values. The melting point was estimated for the two 60-mer SApNPs using the alternative approach devised for hM2e SApNPs. Similar antibody binding affinity, as indicated by the EC_50_ value, was observed for M2ex3 scaffolds and SApNPs across the whole temperature range (4-70°C) (**Figure 3G**, **Figure S4B**), although irregular particle shapes were noted at 70°C in EM micrographs (**Figure S4C**). The interactions of M2ex3 immunogens with two human antibodies (mAb65 and mAb148) were assessed by BLI (**Figure 3H**, **Figure S4D**). The advantage of particulate display was exemplified by the higher binding signals, similar to the case of hM2e SApNPs. Interestingly, the binding profiles (both on-rate and signals) were notably improved for M2ex3-5GS-I3-01v9a-L7P (**Figure 3H**, rightmost) compared to its hM2e counterpart, suggesting that antigenicity may be affected by both the epitope number and spacing on a particular NP scaffold (e.g., I3-01v9a vs. E2p). In summary, tandem M2ex3 SApNPs exhibit greater homogeneity and antigenicity than hM2e SApNPs, while sharing similar yield, structure, and thermostability.

### Protection against influenza A virus challenge by tandem M2ex3 vaccines in mice

BALB/c mice were immunized by ID injection of M2ex3 vaccines adjuvanted with AP or the squalene-based nanoemulsion adjuvant AddaVax (AV) (2.5 µg/footpad, 10 µg total) at weeks 0 and 3. Immunized mice were challenged with LD_50_ × 10 of PR8 (H1N1) at week 6; surviving mice were then challenged with LD_50_ x 10 of HK/68 (H3N2) at week 10 (**Figure 4A**). Survival rate and weight loss were measured for 14 days after each challenge (**Figure 4B**). After the PR8 (H1N1) challenge, all naïve mice died by day 8. In mice immunized with AP-adjuvanted tandem M2ex3 vaccines, 50% of 1TD0 (trimer) mice died by Day 9. Paired with the same AP adjuvant, M2ex3 SApNPs demonstrated higher survival rates: 88% of FR, 100% of E2p, and 100% of I3-01v9a mice survived the H1N1 challenge. In terms of peak weight loss, naïve mice lost the most weight with an average of 21.7 ± 2.9%, and 1TD0 (trimer) mice lost the second highest amount of weight (19.4 ± 7.3%), as expected for the small soluble M2ex3 antigen. In general, M2ex3 SApNP groups showed lower peak weight loss: FR (15.5 ± 8.1%), E2p (11.2. ± 3.8%), and I3-01v9a (15.5 ± 5.3%). Several AV-adjuvanted M2ex3 vaccine groups outperformed their AP counterparts, demonstrating both higher survival rate and lower peak weight loss. The AV-adjuvanted M2ex3 1TD0 (trimer) group showed a slightly higher survival rate at 63% compared to its AP-adjuvanted counterpart (50%). All AV-adjuvanted M2ex3 SApNPs achieved a 100% survival rate, with an improvement noted for the M2ex3 FR group (12% higher survival rate compared to its AP counterpart) and no difference in the E2p and I3-01v9a groups between the two adjuvants (all 100%). For peak weight loss, amongst AV-adjuvanted groups, the M2ex3 trimer showed the highest weight loss with 18.9 ± 5.9%. Most M2ex3 SApNPs adjuvanted with AV showed lower peak weight loss compared with their AP-adjuvanted counterparts: 13.1 ± 7.2% (FR), 12.5. ± 7.7% (E2p), and 10.0 ± 4.2% (I3-01v9a). Inactivated PR8 (H1N1) mice lost the least weight against the strain-matched challenge, with an average of 4.4 ± 5.3%. Next, protection against the second challenge with HK/68 (H3N2) was assessed (**Figure 4B**). While all mice in the naïve group died by day 5, inactivated PR8-immunized mice showed limited protection with a survival rate of 63% against the non-strain-matched challenge. All mice immunized with the M2ex3 vaccines (both AP- and AV-adjuvanted) that survived the prior PR8 challenge also survived the HK/68 (H3N2) challenge. Similar to the survival rate, the highest peak weight loss upon the HK/68 (H3N2) challenge was seen in the naïve group with 19.5 ± 2.9%, followed by the inactivated PR8 H1N1 group with 15.7 ± 4.9%. M2ex3 1TD0 and FR, E2p, and I3-01v9a SApNPs adjuvanted with AP showed lower weight loss with 8.9 ± 5.2%, 4.7 ± 3.2%, 6.5 ± 1.9%, and 4.4 ± 3.0%, respectively (**Figure 4B**). Similarly, M2ex3 1TD0 and SApNP groups adjuvanted with AV showed minimal weight loss: trimer with 4.3 ± 2.1, FR with 4.3 ± 1.0%, E2p with 5.9 ± 3.4%, and I3-01v9a with 4.9 ± 3.3% (**Figure 4B**). Overall, M2ex3 I3-01v9a/AV appeared to provide more effective protection against sequential H1N1 and H3N2 challenges in a mouse model amongst all M2ex3 vaccine formulations tested.

**Figure 4.**
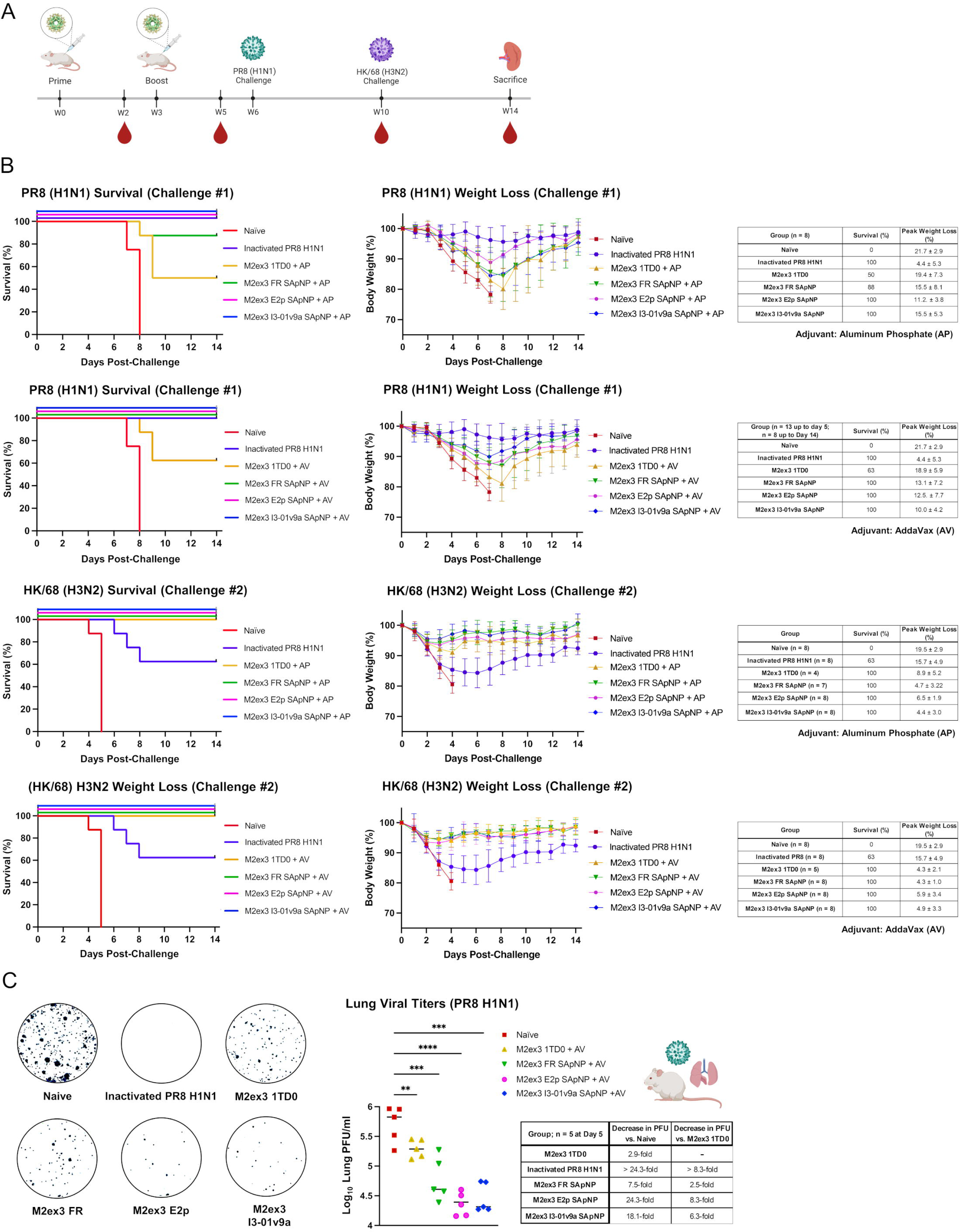
Survival and weight loss assessment of tandem M2ex3 scaffold and nanoparticles in a mouse challenge model. (**A**) Schematic representation of mouse immunization regimen for M2ex3 constructs, sequential intranasal challenges of LD_50_ × 10 of mouse-adapted PR8 (H1N1) and HK/68 (H3N2), blood collection, and sacrifice. Groups were as follows: M2ex3 groups adjuvanted with alum phosphate (n = 8), M2ex3 groups adjuvanted with AddaVax (AV) (n = 13), and inactivated PR8 (H1N1) (n = 8). Inactivated PR8 (H1N1) + AV (n = 13) was used as a positive control for lung viral titers for the PR8 (H1N1) challenge. (**B**) Survival and weight loss of mice challenged with LD_50_ × 10 of PR8 (H1N1) followed by an LD_50_ × 10 of HK/68 (H3N2). Mice were monitored for survival, weight loss, and morbidities for 14 days. (**C**) Lung viral titers in M2ex3-immunized mice on day 5 post-PR8 (H1N1) challenge (n = 5). Visual representation of plaques formed from the lung supernatants of various M2ex3-immunized mice. The highest countable plaques were observed in naïve mice. The lowest number of plaques were observed in the lung supernatants of E2p- and I3-01v9a NP-immunized mice. The assay was performed in duplicate starting at a lung supernatant dilution of 1x followed by 10-fold titrations. Statistical analysis shows significance between M2ex3 groups compared to naïve mice using one-way ANOVA. The error bars indicate mean□±□standard deviation; **p□<□0.01, ***p□<□0.001, and ****p < 0.0001. Images of mouse immunization, virus challenge, and blood and organ collection created with BioRender.com.

The viral load in lungs of mice on day 5 post-PR8 (H1N1) challenge was used as another metric to evaluate the effectiveness of vaccine protection (**Figure 4C**). Mice were immunized and challenged as described above and sacrificed on day 5 post-infection. Lungs were collected, mechanically disaggregated, and centrifuged to pellet cells. PFUs were measured in the supernatants via plaque assay. Overall, naïve mice had the highest virus load, 6.0 × 10^5^ ± 3.3 × 10^5^ PFU/ml. The M2ex3 trimer, FR, E2p, and I3-01v9a groups showed significantly lower viral titers, with 2.9, 7.5, 24.3, and 18.1-fold-lower virus load than the naïve group, respectively. Compared to M2ex3 trimer, FR, E2p, and I3-01v9a showed 2.5, 8.3, and 6.3-fold-lower virus titers in lungs, although this difference was not statistically significant. The mice immunized with inactivated PR8 (H1N1) adjuvanted with AV (positive control) had no detectable viral loads in lungs. Based on the criteria of survival, weight loss, and viral load in lungs, M2ex3 I3-01v9a/AV appeared to be the most effective vaccine among all the SApNP/adjuvant formulations tested.

### Evaluation of M2ex3 vaccine-induced antibody responses

Both M2ex3 E2p and I3-01v9a SApNP groups demonstrated superior serum binding to an M2ex3-5GS-foldon probe at all time points compared to the trimer group **(Figure 5A**, **Figure S5A**). The greatest fold difference between SApNP and 1TD0 (trimer) groups was observed at week 2, suggesting a rapid onset of anti-M2ex3 antibody response elicited by SApNPs. When paired with AP, M2ex3 E2p and I3-01v9a SApNPs yielded 31.6-fold and 83.8-fold higher EC_50_ titers than the 1TD0 trimer, respectively; an even greater fold difference, 47.6 and 102.4 respectively, was noted when these two SApNPs were adjuvanted with AV. The highest EC_50_ titers were observed for the M2ex3 SApNP groups at week 5, at which point M2ex3 E2p and I3-01v9a SApNPs adjuvanted with AP showed 6.7- and 4.9-fold higher EC_50_ values than M2ex3 1TD0 trimer/AP, respectively. M2ex3 E2p and I3-01v9a SApNP adjuvanted with AV showed 5.7- and 4.7-fold higher EC_50_ values than M2ex3 1TD0 trimer/AV at the same time point. Interestingly, M2ex3 FR paired with either adjuvant significantly underperformed M2ex3 E2p and I3-01v9a after the second dose, and trimer at later time points. Notably, AV groups always outperformed AP groups with higher EC_50_ titers, highlighting the importance of a potent adjuvant for eliciting robust M2e-specific antibody responses. Importantly, we also confirmed that the incorporation of two non-human M2e epitopes into the hM2e constructs does not reduce hM2e-specific EC_50_ titers in the M2ex3-immune sera compared to the hM2e-immune sera, as indicated by the hM2e-5GS-foldon probe (**Figure S5B)**. In fact, mouse sera induced by the M2ex3 1TD0 trimer and I3-01v9a SApNP formulated with AV showed similar or higher EC_50_ titers compared to their AP-adjuvanted hM2e counterparts.

**Figure 5.**
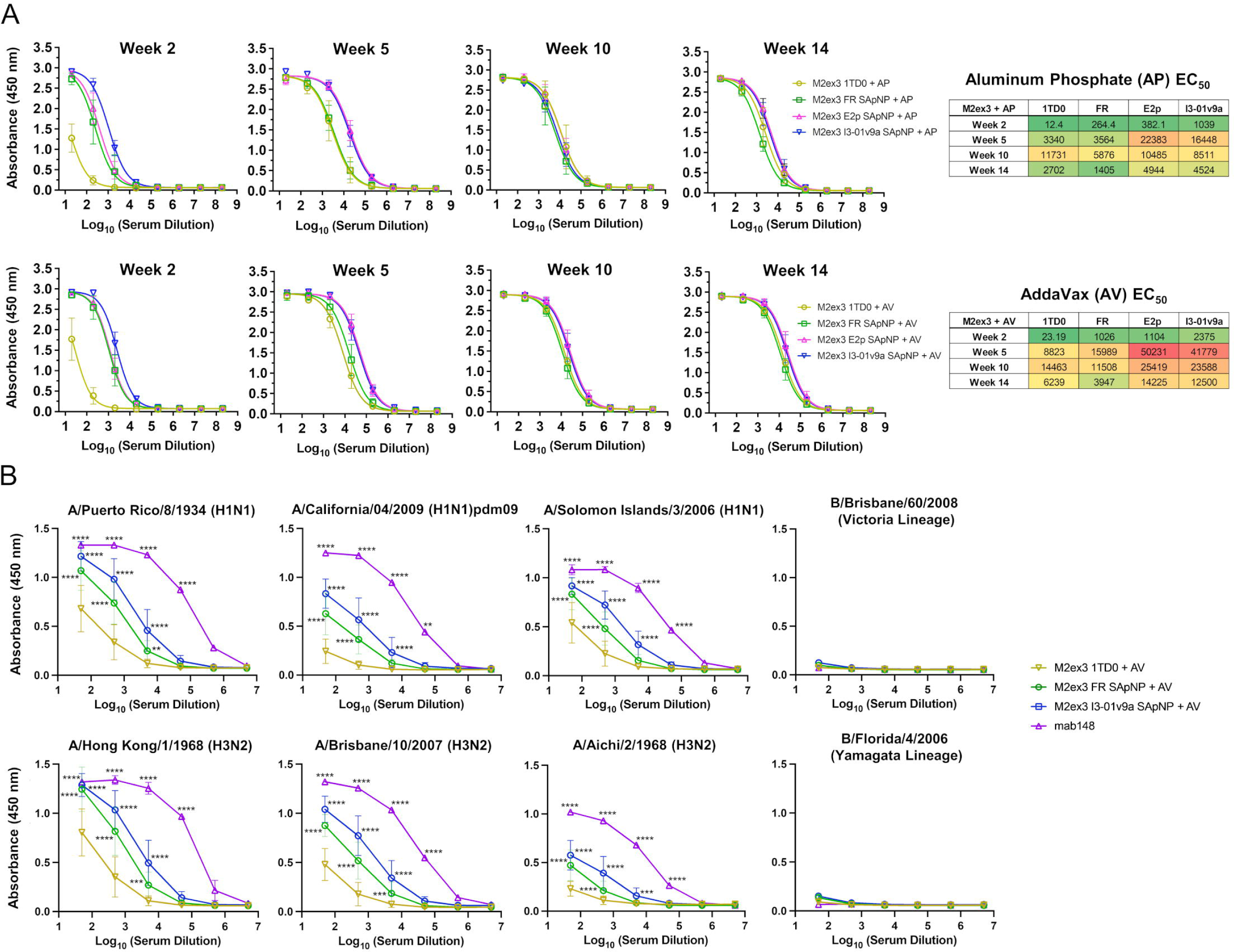
M2ex3-immune sera binding to scaffolded M2ex3 and homotetrameric M2e. (**A**) ELISA curves showing M2ex3-immune sera (adjuvanted with alum phosphate, AP, or AddaVax, AV) binding to the M2ex3-5GS-foldon trimer probe and calculated 50% effective concentration (EC_50_) values for weeks 2, 5, 10, and 14. N = 8 or 13 at weeks 2 and 5. N = variable based on surviving mice/group post-challenge for weeks 10 and 14. The assay was performed in duplicate with a starting serum dilution of 20× followed by seven 10-fold titrations. (**B)** Serum binding to M2e on the surface of MDCK cells infected with various influenza A strains. Strains: A/Puerto Rico/8/1934 (H1N1), A/California/04/2009 (H1N1)pdm09, A Solomon Islands/2/2006 (H1N1), A/Hong Kong/1/1968 (H3N2), A/Brisbane/10/2007 (H3N2), A/Aichi/2/1968 (H3N2), B/Brisbane/60/2008 (Flu B, Victoria Lineage B/Florida/4/2006), and (Flu B, Yamagata Lineage). MAb148 (M2e antibody) was used as a positive control. The assay was performed in duplicate with a starting serum dilution of 50× followed by five 10-fold titrations. Statistical analysis shows significance between trimer and NP groups and positive control using two-way ANOVA. The error bars indicate mean□±□standard deviation; *p□<□0.05, **p□<□0.01, ***p□<□0.001, and ****p□<□0.0001.

The recognition of homotetrameric M2e, which represents the “native” M2e conformation during IAV infection, by M2ex3-immune sera was assessed for the AV-adjuvanted 1TD0, FR, and I3-01v9a groups **(Figure 5B**, **Figure S6**). In this experiment, ELISA was performed to test serum binding to M2e presented on MDCK cells infected with pandemic or seasonal H1N1 and H3N2 strains. The M2ex3 I3-01v9a group demonstrated significantly higher serum binding to M2e expressed on MDCK cells infected with the two challenge strains, A/Puerto Rico/8/1934 (H1N1) and A/Hong Kong/1/1968 (H3N2). Additionally, the M2ex3 I3-01v9a group also showed higher serum binding to pandemic A/California/04/2009 (H1N1) and other H1N1 and H3N2 strains: A/Solomon Islands/2/2006 (H1N1), A/Brisbane/10/2007 (H3N2), and A/Aichi/2/1968 (H3N2). As a negative control, serum binding was also assessed against two IBV strains, which express an M2e that is shorter than and antigenically distinct from IAV M2e. As expected, the M2ex3 1TD0, FR, and I3-01v9a groups showed minimal serum binding to IBV strains B/Brisbane/60/2008 (Victoria Lineage) and B/Florida/4/2006 (Yamagata Lineage). A human M2e antibody in the immunoglobulin form (*82*), termed mAb148 (*77*), was used as a positive control in serum ELISA against IAVs. Based on the in vitro and in vivo evaluation, M2ex3 I3-01v9a SApNP adjuvanted with AV was selected as the lead M2ex3 vaccine candidate for further analysis.

### Distribution, trafficking, and retention of M2ex3 trimers and SApNPs in lymph nodes

Following a similar protocol that was used to analyze HIV-1 and SARS-CoV-2 SApNP vaccines (*56, 74*), we studied the in vivo behavior of the M2ex3 1TD0 trimer and two SApNPs (FR and I3-01v9a) to achieve a better understanding of their interaction with immune cells in lymph nodes and their ability to induce adaptive immune responses. To mount an effective humoral response, these vaccines must be transported through lymphatics and accumulate in lymph node follicles. The immunogens will then be presented to B cells to generate a robust antibody response through crosslinking of B cell receptors (BCRs) (*83–86*). We first studied the transport and distribution of M2ex3-presenting I3-01v9a SApNPs in lymph nodes. We injected a single dose of the immunogen intradermally through the footpads (4 footpads, 10 μg/footpad) and isolated the sentinel brachial and popliteal lymph nodes from both sides of the mouse’s body at 48 h post-immunization. Immunostaining of the excised lymph node sections was carried out using mAb148 and mAb65 (*77, 78*) to detect M2ex3 presented on I3-01v9a SApNPs (**Figure 6A**). Immunohistological images obtained after staining with both antibodies demonstrated a similar distribution of M2ex3 I3-01v9a SApNPs in the center of lymph node follicles (**Figure 6A**, images on the left; **Figure 6B**, schematics on the right). This intra-lymph node distribution pattern is consistent with the results obtained from previous studies assessing SARS-CoV-2 spike SApNPs (*74*), HIV-1 Env SApNPs (*56*) and ovalbumin-conjugated gold NPs (*87, 88*). Due to the better signal-to-noise ratio, mAb148 was used hereafter to examine the trafficking of three M2ex3 immunogens in lymph nodes.

**Figure 6.**
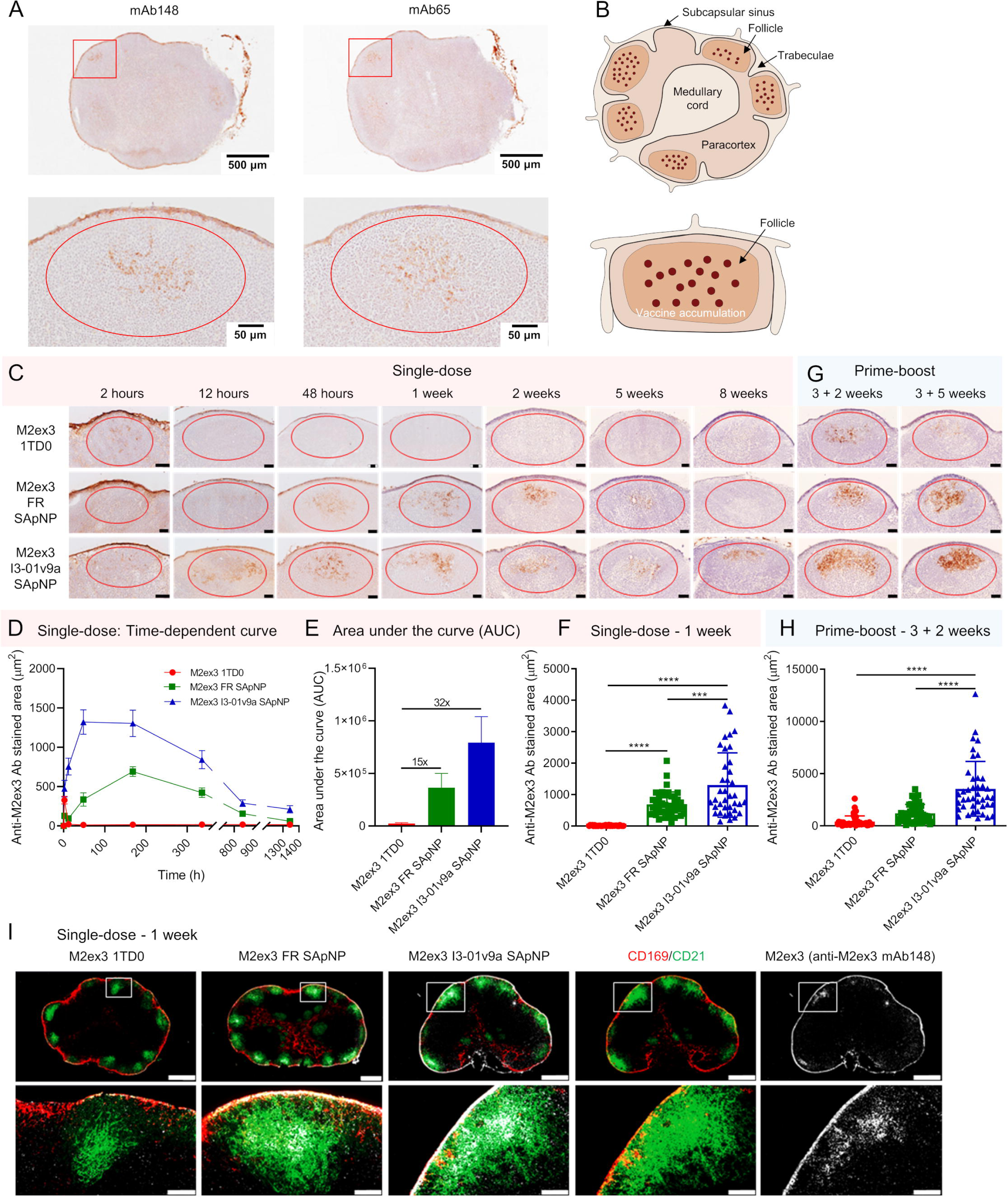
Prolonged retention of M2ex3-presenting SApNPs in lymph node follicles. (**A**) Distribution of I3-01v9a SApNPs displayed M2ex3 trimers in a lymph node at 48 h after a single-dose injection (10 μg/injection, 40 μg/mouse). Anti-M2e Ab148 and Ab65 were used to stain the lymph node tissues. (**B**) schematics of M2ex3 trimer presenting SApNP accumulation in lymph node tissues. (**C**) Trafficking and retention of the M3ex3 trimer and FR and I3-01v9a SApNPs in lymph node follicles at 2 h to 8 weeks after a single-dose injection. Scale bar = 50 μm for each image. (**D**) Time-dependent curve and (**E**) Area under the curve of the Ab148-stained area in immunohistological images of M3ex3 immunogen retention in lymph node follicles up to 8 weeks. (**F**) Quantification of M3ex3 vaccine accumulation in lymph node follicles at 1 week after a single-dose injection. (**G**) Histological images of the M2ex3 trimer and two SApNP vaccine accumulation and retention in lymph node follicles at 2 weeks and 5 weeks after the boost, which occurred at 3 weeks after the first dose. (**H**) Quantification of vaccine accumulation in lymph node follicles at 2 weeks after the boost. In mouse injection, all vaccine immunogens were adjuvanted with AddaVax (AV). Data were collected from more than 10 lymph node follicles (n = 3-5 mice/group). (I) Interaction of M2ex3 trimer presenting SApNPs with FDC networks in lymph nodes at 1 week after a single-dose injection. Both FR and I3-01v9a SApNP immunogens were colocalized with FDC networks. Immunofluorescent images are pseudo-color-coded (CD21^+^, green; CD169^+^, red; Ab148, white). Scale bars = 500 and 100 μm for a complete lymph node and enlarged image of a follicle, respectively. The data points are expressed as mean ± SEM for (D) and SD for (E, F and H). The data were analyzed using one-way ANOVA followed by Tukey’s multiple comparison post hoc test. ***p < 0.001, ****p < 0.0001.

We next studied the trafficking and retention patterns of the M2ex3 trimer and two SApNPs in lymph node follicles up to 8 weeks after a single-dose injection (4 footpads, 10 μg/footpad) (**Figure 6C**). The histological images showed that all M2ex3 immunogens were transported into lymph nodes and accumulated in the subcapsular sinus within 2 h (**Figure 6C**). The M2ex3 trimer was trafficked into follicles within 2 h and completely cleared by 12 h. In contrast, the M2ex3 FR SApNP began to be present in follicles at 12 h, reached peak accumulation at 1 week, and remained detectable up to 5 weeks. The M2ex3 I3-01v9a SApNP demonstrated superior follicular retention with a duration of at least 8 weeks. The mAb148-stained area was quantified in a time-dependent manner, showing a ∼672-fold longer retention for the I3-01v9a SApNP compared to the M2ex3 trimer (**Figure 6C-D**). The area under the curve suggested that the exposure of M2ex3 presented on SApNPs in follicles would be 14-31 times higher than the soluble M2ex3 trimer (**Figure 6E**). Additionally, the M2ex3 FR and I3-01v9a SApNPs also showed 45-86 times greater accumulation compared with the M2ex3 trimer at 1 week (**Figure 6F**). These results are consistent with our previous findings (*56, 74, 87*), in which small particles (< 15 nm) were cleared from lymph node follicles within 48 h, whereas large particles (30-100 nm) remained for weeks. Importantly, M2ex3 I3-01v9a SApNPs displaying M2ex3 antigens or BG505 trimers showed significantly longer follicular retention than those presenting SARS-CoV-2 spikes (∼8 weeks vs. ∼2 weeks), suggesting a correlation between antigen retention and antigen thermostability (a T_m_ value of 65°C or greater for M2ex3 and BG505 Env vs. 48°C for SARS-CoV-2 spike). Of note, a shorter follicular retention (less than 2 weeks) was reported for a newly developed M2e vaccine, in which M2e peptides were encapsulated within a poly(D,L-lactide-co-glycolide), or PLGA, polymer matrix (*51*), possibly due to polymer degradation and/or M2e peptide release. Next, we examined the accumulation and retention patterns of these three M2ex3 immunogens at 2 and 5 weeks using a prime-boost regimen (injected into 4 footpads at weeks 0 and 3, 10 μg/footpad) (**Figure 6G**). A similar pattern to the single-dose injection was observed. Interestingly, M2ex3 trimers were detected in follicles up to 5 weeks after the boost and showed improved retention compared to the single dose, consistent with our previous SARS-CoV-2 study (*74*). Improvement in accumulation and retention after boost was also observed for the M2ex3 FR and I3-01v9a SApNPs. Overall, displaying tandem M2ex3 on the I3-01v9a SApNP showed 8-fold greater accumulation in follicles compared to the soluble 1TD0 trimer 2 weeks after the boost (**Figure 6H**). Prolonged retention of the M2ex3 I3-01v9a SApNP vaccine in lymph node follicles may suggest improved longevity of vaccine-induced immunity.

Follicular dendritic cells (FDCs) located in the center of lymph node follicles are essential for retention and presentation of native-like antigens to stimulate B cell responses (*83, 85, 86*). FDCs can collect and align soluble antigens and large particles such as immune complexes, viruses, and bacteria on their surfaces or dendrites through a complement receptor-dependent mechanism to generate and maintain GC reactions (*84–87, 89, 90*). Our previous studies of ovalbumin-conjugated gold NPs (*87*), and SARS-CoV-2 and HIV-1 antigen-presenting SApNPs (*56, 74*) suggest that FDC networks may be the key cellular component to retain M2ex3 SApNPs. To test this possibility, we collected lymph nodes at the peak of SApNP accumulation (1 week) and other timepoints (48 h to 8 weeks) following a single-dose injection (**Figure 6I**, **Figure S7A-D**). Lymph node tissues were stained with mAb148 (white) for M2ex3, anti-CD21 antibodies (green) for FDCs, and anti-CD169 antibodies for subcapsular sinus macrophages. The signals for both M2ex3 FR and I3-01v9a SApNPs (mAb148 binding) showed their colocalization with FDCs (CD21^+^) at 1 week (**Figure 6I**), confirming the retention of M2ex3 SApNPs in FDC networks.

### Characterization of GC reactions induced by M2ex3 trimers and SApNPs

In GCs, B cells undergo somatic hypermutation, selection, affinity maturation, and class switching, eventually becoming memory or antibody-secreting plasma cells (*84, 91–94*). FDC networks and T follicular helper (T_fh_) cells support GC formation and maintenance (*95, 96*). Here, we hypothesized that M2ex3 SApNPs retained by FDC networks could induce more robust and long-lived GC reactions in lymph node follicles compared to the soluble M2ex3 trimer. First, we examined whether M2ex3 I3-01v9a SApNPs can induce strong GC reactions after single-dose injections (4 footpads, 10 μg/footpad). Vaccine-induced GC B cells (GL7^+^, red) and T_fh_ cells (CD4^+^ Bcl6^+^, co-labeled with cyan and red) were characterized by immunohistology. We observed large GCs attached to FDC networks (CD21^+^, green) with organized dark zone (DZ) and light zone (LZ) compartments in follicles (B220^+^, blue) (**Figure 7A**, left). In addition to antigen retention and presentation by FDC networks, T_fh_ cells appear in the LZ of GCs to support B cell stimulation (**Figure 7A**, right). Next, we performed immunohistological analysis on the M2ex3 trimer and two SApNPs at 2, 5, and 8 weeks after a single-dose injection (**Figure 7B**, **Figure S8A-C**) and at 2 and 5 weeks after the boost (**Figure 7C**, **Figure S8D-E**). Following a previously established protocol (*56, 74*), we quantified GC reactions using two metrics: GC/FDC ratio (the frequency of GC formation associated with FDC networks) and size of GCs. The M2ex3 trimer and both SApNPs induced robust GCs, with the M2ex3 I3-01v9a SApNP showing the largest GCs at 2 weeks after a single-dose injection (**Figure 7B**, **Figure S8A**). As expected, the GC/FDC ratio and GC size declined over time in all groups. Notably, while the GC/FDC ratio for the M2ex3 trimer group decreased to ∼50%, the M2ex3 I3-01v9a SApNP generated strong and durable GC reactions that lasted for 8 weeks (**Figures 7B** and **7D**, **Figure S8C**). GCs were restored for all vaccine groups after the boost, but the GC/FDC ratio for the trimer group decreased again significantly at 5 weeks after the boost. Overall, the M2ex3 I3-01v9a SApNP generated GCs 2.5 times the size of those elicited by the M2ex3 1TD0 trimer after one dose (**Figures 7B** and **7D**) and 1.7 times the size after the boost at 8 weeks (**Figures 7C** and **7E**).

**Figure 7.**
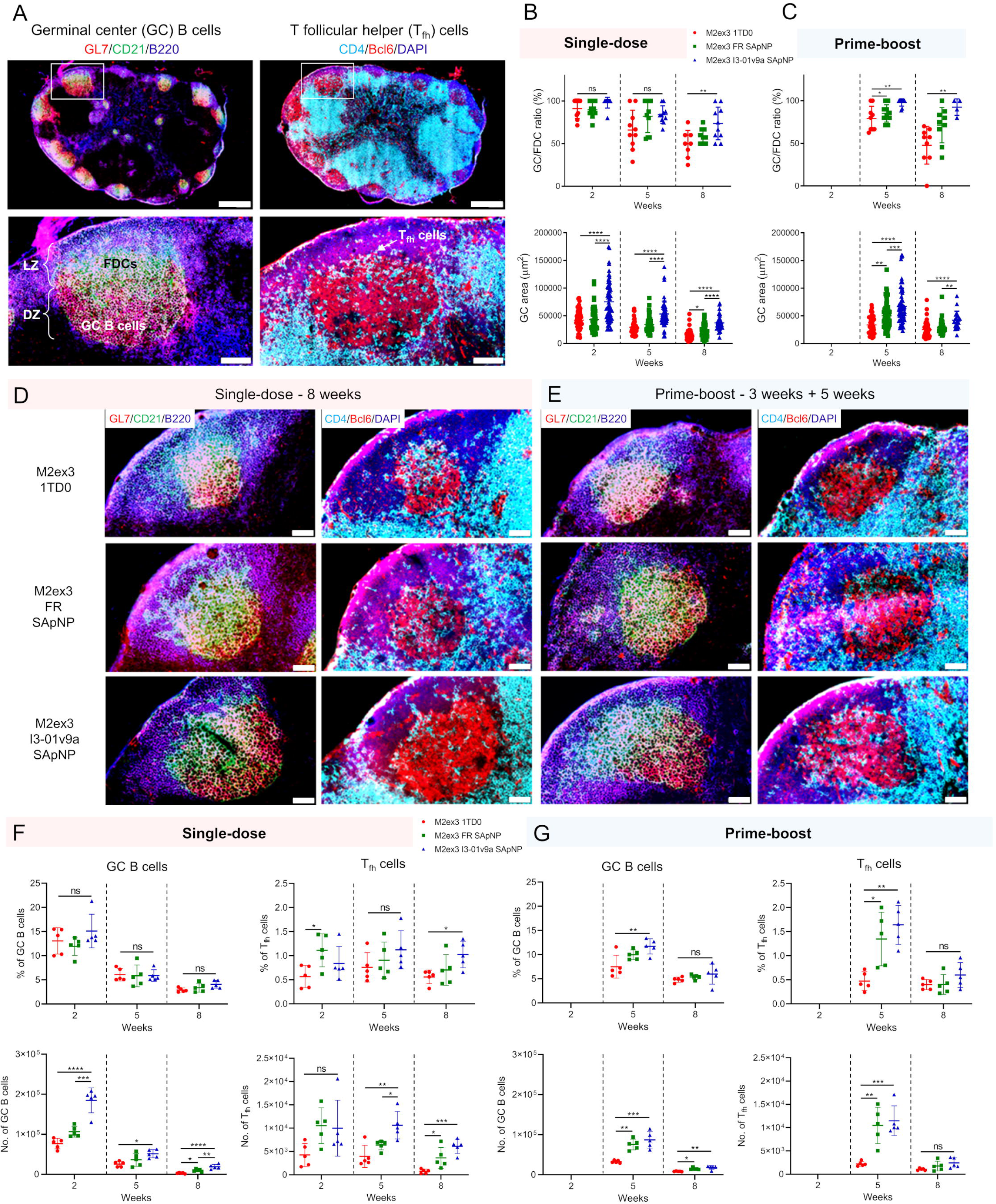
Induction of robust and long-lived germinal center reactions by M2ex3-presenting SApNPs. (**A**) Top image: Immunofluorescent images of M2ex3 trimer presenting I3-01v9a SApNP vaccine candidate induced germinal centers (GCs) at 2 weeks after a single-dose injection (10 μg/injection, 40 μg/mouse). Bottom image: robust GC reaction with organized light zone (LZ) and dark zone (DZ) compartments in lymph node follicles. GC B cells (GL7^+^, red) attached to FDCs (CD21^+^, green) and T_fh_ cells located in LZ of GCs. Scale bars = 500 and 100 μm for a complete lymph node and the enlarged image of a follicle, respectively. (**B**) and (**C**) quantification of GCs in terms of the GC/FDC ratio and the size of GCs induced by the M2ex3 trimer, and FR and I3-01v9a SApNP vaccines using immunohistological images at 2, 5, and 8 weeks after a single-dose injection or at 2 and 5 weeks after the boost, which occurred at 3 weeks after the first dose (n = 5 mice/group). (**D**) and (**E**) representative GC images induced by three M2ex3 vaccine constructs at 8 weeks using a single-dose or prime-boost regimen. Scale bar = 50 μm for the image of an enlarged lymph node follicle. (**F**) and (**G**) Quantification of GC reactions in terms of the percentage and number of GC B cells and T_fh_ cells using flow cytometry after a single-dose or prime-boost immunizations. In mouse immunization, all M2ex3 vaccines were adjuvanted with AddaVax (AV). The data points are shown as mean ± SD. The data were analyzed using one-way ANOVA followed by Tukey’s multiple comparison post hoc test for each timepoint. *p < 0.05, **p < 0.01, ***p < 0.001, ****p < 0.0001.

The GCs were further analyzed by flow cytometry. We collected sentinel lymph nodes at 2, 5, and 8 weeks after a single dose of M2ex3 1TD0 trimer, FR SApNP, or I3-01v9a SApNP (2.5 μg/footpad, 10 μg total) (**Figure 7F**, and **Figure S9**), and at 2 and 5 weeks after the boost (**Figure 7G**) following the prime-boost regimen. The lymph node tissues were disaggregated into a single cell suspension and stained with an antibody cocktail. The percentage and number of GC B cells and T_fh_ cells were used to quantify the GC reactions, which correlate with the immunohistological results (**Figures 7A-E**). Flow cytometry indicated that M2ex3 I3-01v9a SApNP elicited the largest GC B cell and T_fh_ cell populations after a single-dose injection (**Figure 7F**). GC reactions peaked at 2 weeks for all tested groups and declined over time. While the M2ex3 trimer failed to maintain the GC reactions, the M2ex3 I3-01v9a SApNP induced durable GC reactions lasting for 8 weeks (**Figure 7F**). Both the frequency and number of GC B cells and T_fh_ cells could be improved by a boost injection (**Figure 7G**). Interestingly, a significant expansion of T_fh_ cells was noted for the M2ex3 FR and I3-01v9a SApNPs 2 weeks after the boost. Overall, the M2ex3 I3-01v9a SApNP elicited 5.7/1.1 times more GC B cells and 7.0/1.3 times more T_fh_ cells compared with the soluble trimers at 8 weeks after the single-dose/prime-boost injections, respectively (**Figures 7F** and **7G**). In summary, our immunological analysis suggests that M2ex3 I3-01v9a SApNP can generate long-lived GC reactions in lymph nodes more effectively than individual M2ex3 trimers, resulting in potent and long-lasting M2ex3-specific humoral immunity.

### Antibody-dependent cell cytotoxicity (ADCC) and functional T cell responses

The non-neutralizing M2e-specific antibody responses were evaluated for functional activity using a surrogate ADCC assay incorporating a luciferase reporter. The M2ex3 I3-01v9a-immune sera elicited significantly higher luciferase activity, measured in relative light units (RLUs), than naïve, M2ex3 1TD0 trimer, and M2ex3 FR groups, indicating detection of M2e antibodies by mouse Fcγ receptor IV (mFcγRIV) expressed on Jurkat cells (**Figure 8A**). The largest difference in ADCC activity between different M2ex3 vaccine groups was observed at the lowest serum dilution tested, 20×, with M2ex3 I3-01v9a showing 7.2-fold, 5.9-fold, and 2.1-fold higher RLU values than naïve, M2ex3 1TD0 trimer, and M2ex3 FR groups, respectively. This assay thus confirmed that M2ex3-immune sera binding to homotetrameric M2e expressed on virus-infected cells has the potential to activate ADCC pathways, which is an important mechanism for M2e-mediated protection.

**Figure 8.**
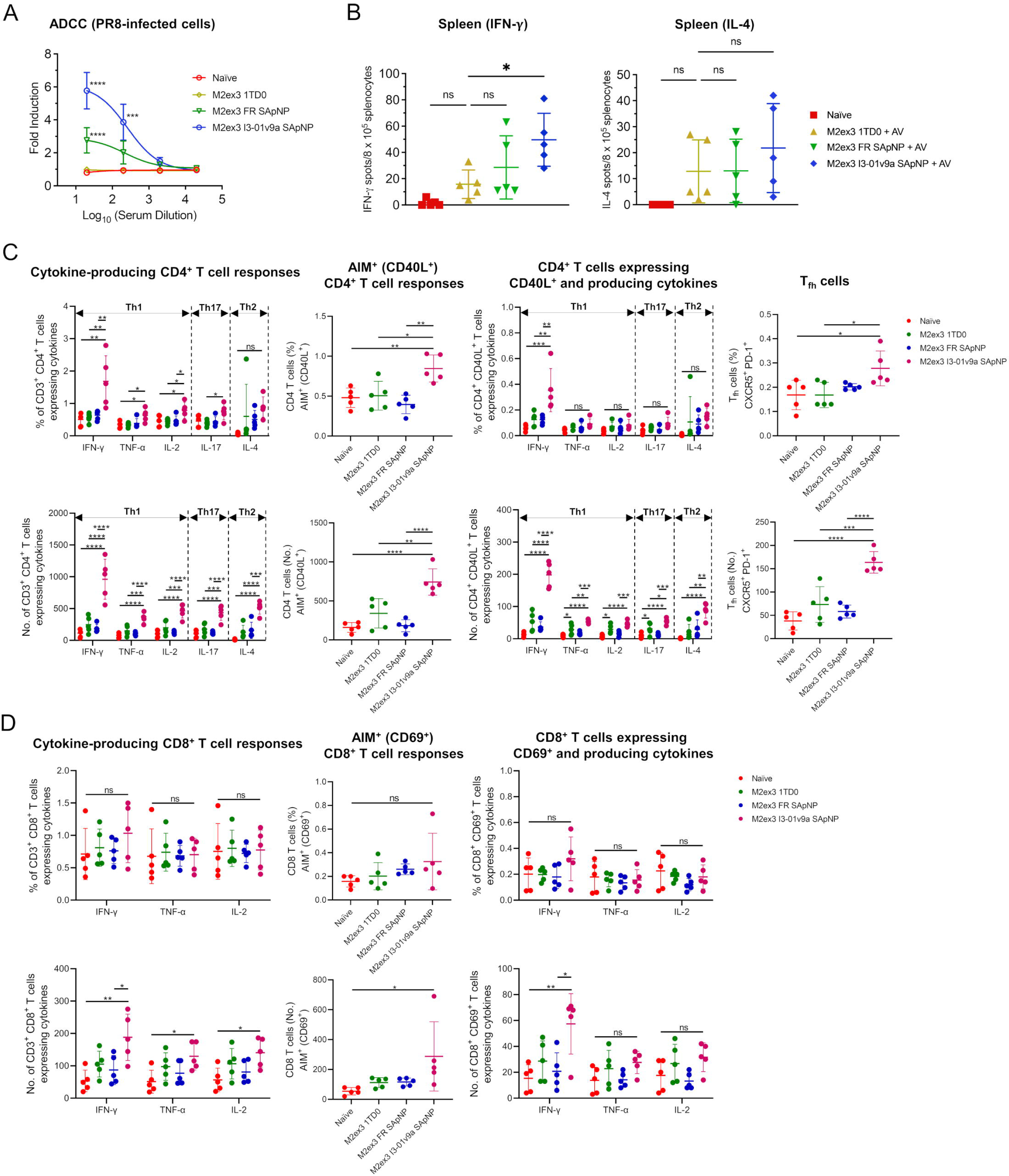
Innate and T cell responses of M2ex3 scaffolds and M2ex3-presenting SApNPs. (**A**) ADCC activity measured by RLU of FcγRIV-expressing Jurkat effector cells binding to M2ex3-immune sera bound to M2e on PR8 (H1N1)-infected MDCK cells. The data is presented as mean ± SEM (**B**) Spot formation of IFN-γ and IL-4-secreting splenocytes from M2ex3-immunized mice on day 5 post-PR8 (H1N1) challenge. Mouse splenocytes were isolated from immunized mice with M2ex3 trimer and FR and I3-01v9a SApNPs (adjuvanted with AV) at 5 days post-PR8 (H1N1) challenge following the prime-boost immunization (n = 5 mice/group). Splenocytes of naïve mice without immunization but with a H1N1 virus challenge were included as control samples. Quantification of the percentage and number of vaccine-induced functional (**C**) CD4^+^ and (**D**) CD8^+^ T cell responses using flow cytometry. In mouse immunization, all M2ex3 vaccines were adjuvanted with AddaVax (AV). The data points are shown as mean ± SD. The data were analyzed using one-way ANOVA followed by Tukey’s multiple comparison post hoc test for each timepoint. *p < 0.05, **p < 0.01, ***p < 0.001, ****p < 0.0001.

While antibody-mediated neutralization plays a key role in blocking virus infection, T cell-mediated cellular immunity effectively reduces disease severity, hospitalization, and death rate (*97, 98*). Enzyme-linked immunosorbent spot (ELISpot) analysis demonstrated that the M2ex3 I3-01v9a SApNP group produced notably higher spot formation in bulk IFN-γ-secreting splenocytes stimulated with the M2ex3-5GS-foldon trimer probe compared to the naïve (> 28-fold), M2ex3 trimer (7-fold), and M2ex3 FR (2.5-fold) groups per 8 × 10^5^ splenocytes (**Figure 8B**). Similarly, the M2ex3 I3-01v9a group also produced, on average, more spots in IL-4-secreting splenocytes compared to the naïve (21.8-fold), M2ex3 trimer (1.7-fold), and M2ex3 FR (1.7-fold) groups, although the findings were not statistically significant.

T cells from splenic tissue can be divided into several subsets, including CD4^+^ helper cells, which have multiple central roles in orchestrating adaptive immune responses, and CD8^+^ cytotoxic T cells, which control virus infection by killing virus-infected cells and producing effector cytokines (*99–102*). Here, we designed a 13-color panel to analyze functional T cell responses by measuring activation induced marker (AIM) and intracellular cytokine staining (ICS) using flow cytometry. We compared various M2ex3 vaccine constructs for the induction of CD4^+^ and CD8^+^ responses specific to the vaccine antigen at 5 days after the PR8 (H1N1) challenge following a two-dose immunization regimen (**Figure 4A**). Mouse splenocytes from the M2ex3 1TD0 trimer, FR SApNP, and I3-01v9a SApNP groups were stimulated with the M2ex3-5GS-foldon trimer prior to analysis. All three M2ex3 constructs generated balanced Th1 and Th2 responses and relatively lower Th17 responses (**Figure 8C**, **Figure S10**). Among the three vaccines, M2ex3 I3-01v9a SApNP was most effective in terms of the frequency and number of intracellular cytokine (IFN-γ, TNF-α, IL-2, IL-17 and IL4)-producing CD4^+^ T cells, CD40L^+^CD4^+^ T cells, and CD40L^+^CD4^+^ T cells that produce intracellular cytokines, as well as T_fh_ cells. Importantly, M2ex3 I3-01v9a SApNP induced significantly more IFN-γ-producing functional CD4^+^ T cells, which included 2.7 times more activated (CD40L^+^) T cells than the M2ex3 trimer, resulting in a Th1-skewed response. Similarly, M2ex3 I3-01v9a SApNP elicited more intracellular cytokine (IFN-γ, TNF-α, IL-2)-producing CD8^+^ T cells, CD69^+^CD8^+^ T cells, and CD69^+^CD8^+^ T cells that produce IFN-γ than naïve, M2ex3 trimer, and M2ex3 FR groups (**Figure 8D**). Overall, M2ex3 I3-01v9a SApNP induced stronger functional CD4^+^ and CD8^+^ T cell responses than M2ex3 trimer and FR SApNP, consistent with the higher survival rate, lower weight loss and viral loads (**Figure 4**), greater M2e-specific antibody responses (**Figure 5**), and more pronounced ADCC activity (**Figure 8A**).

### Evaluation of single-dose M2ex3 SApNP vaccine formulations in mice

To probe the limits of protection conferred by the M2ex3 SApNP vaccine, mice were immunized with a single low dose of M2ex3 I3-01v9a SApNP combined with a TLR9 agonist (unmethylated deoxycytidine-deoxyguanosine or CpG) or a STING agonist (a cyclic dinucleotide analog named MIW815 (*103*)), followed by an LD_50_ × 10 challenge with PR8 (H1N1) (**Figure 9A**). CpG has previously been demonstrated to be safe and effective in a SARS-CoV-2 clinical trial (*104, 105*), and a STING agonist was recently shown to be effective for a pan-sarbecovirus vaccine in mice, rabbits, and non-human primates (*106*). AP, in combination with a high (10 µg) and low (2 µg) SApNP dose, was included here as a control adjuvant so that the same control adjuvant would be used across all immunization-challenge experiments performed in this study. After the challenge, all naïve mice died by day 9 (**Figure 9B**). Mice immunized with 10 μg M2ex3 I3-01v9a SApNP adjuvanted with AP had a 60% survival rate. However, when the SApNP dose was lowered to 2 μg, only 20% of mice survived the challenge. In contrast, both groups of mice immunized with 2 μg of M2ex3 I3-01v9a SApNP adjuvanted with either CpG or MIW815 showed survival rates of 90%. Among the vaccinated mice, mice that received 10 μg or 2 μg of M2ex3 I3-01v9a SApNP with AP adjuvant lost the most weight, with a peak weight loss of 20.1 ± 4.3% and 21.4 ± 4.4%, respectively. Mice that were immunized with 2 μg of M2ex3 I3-01v9a SApNP paired with either CpG or MIW815 lost the least weight, with a peak weight loss of 13.4 ± 6.2% and 14.7 ± 5.0%, respectively. Mice immunized with 2 μg of M2ex3 I3-01v9a SApNP adjuvanted with either CpG or MIW815 also showed the highest sera binding against the M2ex3-5GS-foldon trimer probe. Both groups yielded 2.7-fold higher EC_50_ values than the high-dose (10 µg SApNP) AP group (**Figure 9C, Figure S11**). At the same SApNP dose (2 μg), the CpG and MIW815 groups showed 6.5-fold and 6.3-fold higher M2ex3-specific antibodies compared to the AP group, respectively. Overall, our results suggest that a single low dose of M2ex3 I3-01v9a SApNP, when formulated with a potent adjuvant, is sufficient to protect mice against a lethal influenza A challenge.

**Figure 9.**
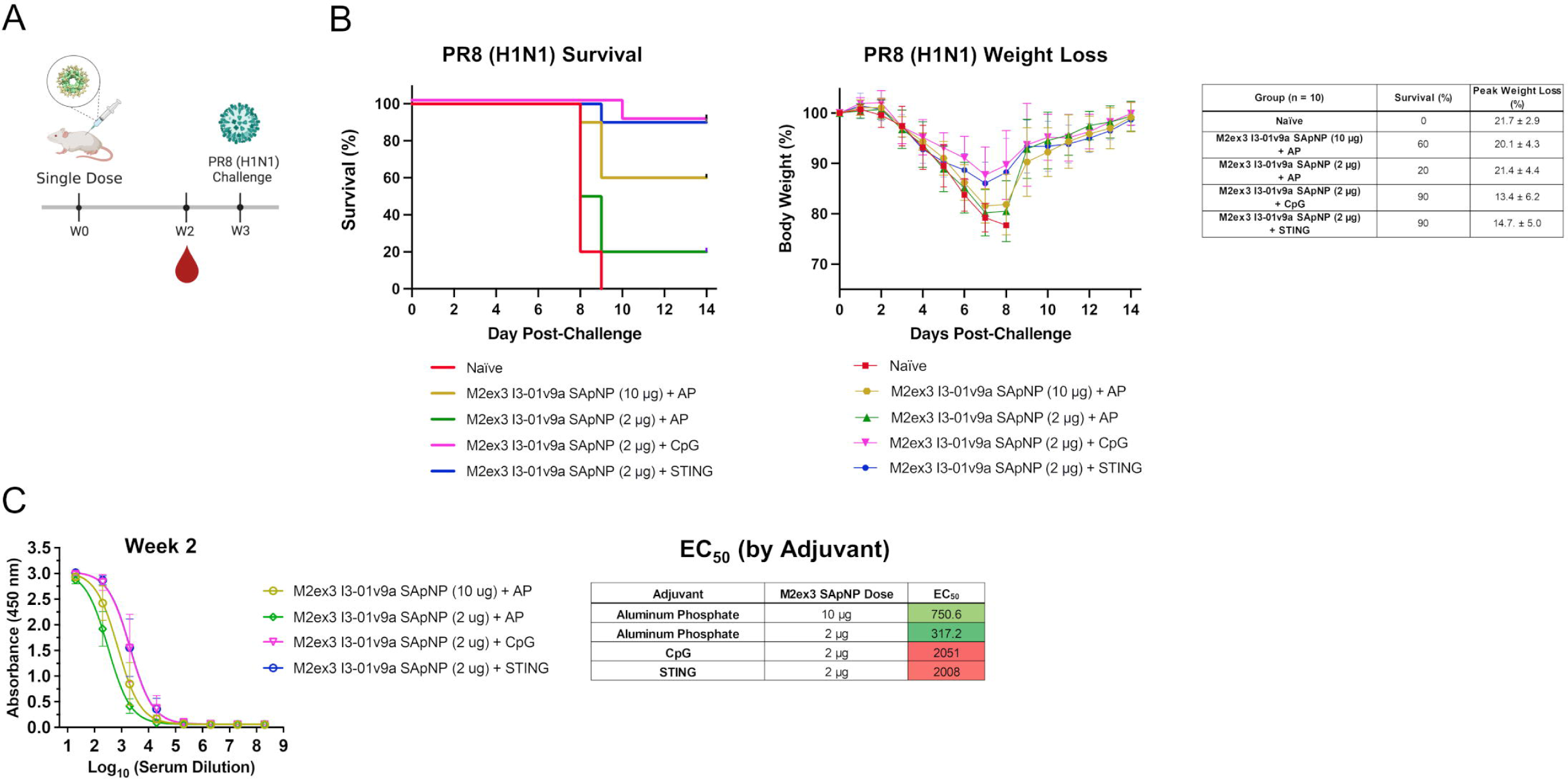
Protection against influenza A virus challenge by a single-dose M2ex3 SApNP vaccine with potent adjuvants in mice. (**A**) Schematic representation of the single-dose mouse immunization regimen for M2ex3 I3-01v9a SApNP with aluminum phosphate (AP), CpG, or a STING agonist (MIW815) adjuvant followed by an intranasal challenge with LD_50_ × 10 of mouse-adapted PR8 (H1N1). Blood collection was carried out at week 2. Groups were as follows: M2ex3 I3-01v9a SApNP (10 μg) + AP, M2ex3 I3-01v9a SApNP (2 μg) + AP, M2ex3 I3-01v9a SApNP (2 μg) + CpG, and M2ex3 I3-01v9a SApNP (2 μg) + STING agonist (n = 10). (**B**) Survival and weight loss data for single-dose-immunized mice that were challenged with PR8 (H1N1). Mice were monitored for survival, weight loss, and morbidities for 14 days. (**C**) ELISA curves showing M2ex3-immune sera (adjuvanted with AP, CpG, or STING agonist) binding to the M2ex3-5GS-foldon trimer probe and calculated 50% effective concentration (EC_50_) values for week 2 (n = 10). Serum ELISA was performed in duplicate with a starting serum dilution of 20× followed by seven 10-fold titrations.

## DISCUSSION

Since the 1918 Spanish flu pandemic, influenza has caused millions of deaths and hospitalizations worldwide and remains a serious public health concern. From 2010 to 2017, seasonal flu caused 9.2 to 35.6 million reported cases of influenza and 140,000 to 710,000 hospitalizations. Each year, seasonal flu causes an estimated 3-5 million cases of severe illness worldwide (*1*). Conventional influenza vaccines, such as inactivated viruses produced in eggs, have been shown to significantly reduce disease burden. However, these vaccines mainly induce NAbs against viral epitopes that are prone to antigenic drift, allowing viruses to evade vaccine-induced immune responses (*17*). As a result, annual updates are necessary for seasonal flu vaccines, even though they may not offer sufficient protection. Additionally, recent evidence has revealed the negative impact of repeated antigen exposure on vaccine efficacy (*107*). Therefore, it is important to develop a vaccine that can provide broad and durable protection against diverse influenza viruses.

An M2e vaccine capable of eliciting durable antibody responses to M2e from diverse IAV subtypes and hosts may be developed towards an effective universal pan-influenza A vaccine. M2e antibodies are non-neutralizing but can engage alveolar macrophages and natural killer cells to promote viral clearance via ADCC (*108, 109*). In principle, a successful M2e vaccine could act as a standalone pan-influenza A vaccine, reducing the severity of disease caused by pandemic strains originating from various animal reservoirs that contain novel HA proteins against which the majority of the human population lacks pre-existing immunity (*17*). Alternatively, M2e can be used to complement seasonal influenza vaccines, significantly boosting the breadth of these strain-specific inactivated virus vaccines (*110*). However, to date, an approved M2e vaccine remains elusive. Here, we approached M2e vaccine development with a rational strategy, in which newly developed single-component, multilayered SApNPs were used as carriers to deliver 60 identical M2e antigens to overcome the intrinsically poor immunogenicity associated with soluble M2e and increase durability of M2e-specific immunity. Notably, we have previously developed vaccine candidates for HIV-1 (*56, 61*), HCV (*59*), SARS-CoV-2 (*57*), and EBOV (*58*) based on the SApNP platform. In this study, we first tested this strategy using hM2e and then presented a tandem M2e antigen, derived from human, avian, and swine IAVs, on FR and two larger 60-meric SApNPs, E2p and I3-01v9a. All the M2e vaccine constructs were evaluated in vitro and in vivo.

Our approach may address the two limitations of previous M2e-based vaccine candidates that were tested in clinical trials: poor immunogenicity and poor durability. Multivalent display of M2e on the surface of 60-meric SApNPs significantly improves the immunogenicity of M2e. The large size and high thermostability of the M2ex3-presenting I3-01v9a SApNP allows for long retention in the lymph nodes follicles (8 weeks or longer), resulting in robust and prolonged GC reactions compared to those elicited by the scaffolded M2e trimer. Importantly, the M2ex3 I3-01v9a SApNP exhibited identical trafficking and retention patterns to the same SApNP presenting 20 highly glycosylated HIV-1 envelope glycoprotein (Env) trimers (*56*), suggesting a minimal impact of glycan content on vaccine transport and retention in lymph node follicles. Combining the M2ex3 I3-01v9a SApNP with commonly used adjuvants such as AP or AV resulted in potent M2e-specific antibody and functional T cell responses that likely conferred protection against sequential H1N1 and H3N2 challenges. In contrast, the inactivated PR8 (H1N1) virus vaccine and a strain-matched challenge offered minimal protection against a follow-up heterologous challenge from a different subtype (H3N2), suggesting that the immunity induced by the previous H1N1 challenge could not protect against a subsequent H3N2 challenge. In comparison, the M2e SApNP-immunized groups showed minimal weight loss against an H3N2 challenge following an H1N1 challenge. Therefore, unlike the inactivated PR8 (H1N1) group, immunizations with M2e SApNP followed by an H1N1 challenge may enhance protection against heterosubtypic H3N2, addressing a major limitation of seasonal inactivated vaccines, which offer mostly strain-specific protection. Overall, our results suggest that the adjuvanted M2ex3 I3-01v9a SApNP can be developed into an effective, cross-protective influenza vaccine that may overcome the limitations of currently marketed influenza vaccines, including the lack of protection against antigenically drifted seasonal or novel pandemic strains (*111*). Furthermore, we also assessed our lead vaccine candidate, M2ex3 I3-01v9a SApNP, in a single, low-dose formulation with potent adjuvants. Both CpG and MIW815 (*103*) were found to significantly improve protection from lethal influenza A challenge compared to a conventional adjuvant, AP. These results align with our recent study (*74*), where we evaluated a SARS-CoV-2 spike SApNP vaccine mixed with adjuvants that target diverse immune signaling pathways. The I3-01v9a SApNP adjuvanted with either CpG or a STING agonist (MIW815 (*103*)) induced more effective Th1-biased CD4^+^ and CD8^+^ T cell responses, in addition to neutralizing titers, compared to non-adjuvanted vaccine groups (*74*). Our study also demonstrated that CpG and MIW815 (*103*) substantially increased anti-M2e antibody titers compared to AP.

Additionally, our SApNP approach is significantly different from previous M2e vaccine strategies. In this study, a tandem M2e antigen derived from human, avian, and swine influenza A viruses was presented on our newly developed SApNP platforms (*56–58*). The multivalent display allows M2e epitopes to directly engage and cluster B cell receptors (BCRs) to stimulate a robust B cell response. In recent studies, M2e peptides were encapsulated within a liposome (*48*) or a polymer matrix (*49–51*) as nanovaccines against influenza, for which the degradation rates of the biomaterials used, the release kinetics of the M2e peptide, and how M2e peptides interact with B cells in vivo can be difficult to quantify. Furthermore, different methods have been used to analyze intra-lymph node transport for M2e-based nanovaccines. In the current study, the trafficking and retention of M2ex3 I3-01v9a SApNP were quantified via immunostaining of tandem M2e antigens displayed on SApNPs using an M2e-specific antibody (mAb148) at individual time points. In comparison, a previous study conjugated fluorescent dyes to the M2e peptide (*51*), which may have influenced the experimental readouts due to signal degradation in vivo over time.

Future studies will focus on several fronts. First, the concept of a “combination vaccine” that includes an M2e component warrants investigation. While M2ex3 SApNP can be used as a standalone vaccine, it may also be combined with a seasonal flu vaccine, such as an inactivated virus vaccine, to boost protection against non-vaccine-matched circulating strains and potential pandemic strains. Second, an M2ex3 SApNP vaccine may reduce the severity of disease caused by highly pathogenic H5N1 and H7N9 influenza strains. Third, the mechanistic analysis of M2ex3 SApNP/adjuvant formulations, including their retention in FDC networks and interactions with various immune cell populations in lymph node cellular compartments, will be investigated in our future studies. Lastly, less commonly explored influenza B M2e can be incorporated (in tandem) with influenza A M2ex3 and displayed on SApNPs to potentially yield a truly universal, single-component vaccine against both influenza A and B viruses. Given the sequence conservation of M2e across HA subtypes, the M2e vaccines developed in this study may protect against diverse IAVs, which account for ∼70% of all influenza cases annually, thereby significantly reducing the deaths and hospitalizations associated with influenza worldwide.

## METHODS

### Computational design of I3-01v9a for optimal nanoparticle display of monomeric antigens

We redesigned the N-terminus of I3-01v9 for optimal surface display of monomeric antigens, such as M2e, without using long linkers. Because I3-01v9 and I3-01 share nearly identical NP structures (*56*), the I3-01 structure (PDB ID: 5KP9) was used here for all modeling purposes. We manually extended the I3-01 N-terminal helix as an initial model. Briefly, an α-helix (residues 953-982) of the transcription factor protein c-MYC (PDB ID: 6G6L) was grafted onto an I3-01 subunit by using Glu2 and Glu3 of I3-01 for fitting. The grafted N-terminal helix was truncated to 11 residues so that the first residue would be just above the surface after particle assembly. The CONCOORD suite (*75*) was used to sample 1000 structures for the modified I3-01 subunit to facilitate ensemble-based protein design. Using default OPLS-UA parameters and a *damp* parameter of 0.1, geometric constraints were generated by the program dist as input for the program disco to generate slightly perturbed conformations (*75*). An ensemble-based design method used in our previous studies (*56, 59*) was employed to predict the first 9 of the 11 residues in the extended N-terminal helix using C_α_ and C_β_-based RAPDF scoring functions (*76*). Given a scoring function, Monte Carlo simulated annealing minimization (*56, 59*) was performed to predict the amino acid composition for each of the 1000 CONCOORD-derived conformations. For each position of the 9-residue helical segment, the frequency of each amino acid type was calculated from the entire ensemble. The final design, I3-01v9a, was determined manually by combining results from both scoring functions.

### Expression and purification of various M2e immunogens

Rationally designed hM2e and tandem M2ex3 scaffolds and SApNPs were characterized in vitro and in vivo. Scaffolded trimers and SApNPs were transiently expressed in ExpiCHO^TM^ cells (Thermo Fisher) using a previously described protocol (*56, 58*). Briefly, ExpiCHO^TM^ cells were thawed and incubated with ExpiCHO^TM^ Expression Medium (Thermo Fisher) in a shaker incubator at 37°C, 135 rpm, and 8% CO_2_. When cells reached a density of 10×10^6^ cell/ml, ExpiCHO^TM^ Expression Medium was added to reduce cell density to 6×10^6^ cell/ml for transfection. ExpiFectamine^TM^ CHO/plasmid DNA complexes were prepared for 100-ml transfections in ExpiCHO cells following the manufacturer’s instructions. For all constructs tested in this study, 100 μg of antigen plasmid and 320 μl of ExpiFectamine^TM^ CHO reagent were mixed in 7.7 ml of cold OptiPRO™ medium (Thermo Fisher). After the first feed on day 1, ExpiCHO cells were cultured in a shaker incubator at 32°C, 120 rpm, and 8% CO_2_ following the Max Titer protocol with an additional feed on day 5 (Thermo Fisher). Culture supernatants were harvested 13-14 days after transfection, clarified by centrifugation at 4000 rpm for 20 min, and filtered using a 0.45 μm filter (Thermo Fisher). For all constructs, the M2e trimer or SApNP was extracted from the culture supernatants by using mAb148 or mAb65 antibody columns. The bound protein was eluted three times by 5 ml of glycine buffer (0.2M glycine, pH 2.2) and neutralized by adding 0.375 ml Tris-base Buffer (2M Tris, pH 9.0). Eluates were pooled and buffer exchanged via ultra-centrifugal filtration to phosphate buffer saline (PBS). The size of trimers and SApNPs was analyzed by size exclusion chromatography using AKTA pure 25 (Cytiva). Trimer was purified on a Superdex 75 Increase 10/300 GL column (GE Healthcare), whereas SApNPs were characterized on a Superose 6 10/300 GL column. Protein concentration was determined using UV_280_ absorbance with theoretical extinction coefficients.

### SDS-PAGE and BN-PAGE

The trimer and SApNPs were analyzed by sodium dodecyl sulfate-polyacrylamide gel electrophoresis (SDS-PAGE) and blue native-polyacrylamide gel electrophoresis (BN-PAGE). The proteins were mixed with loading dye and added to either a 10% Tris-Glycine Gel (Bio-Rad) or a 4-12% Bis-Tris NativePAGE^TM^ gel (Life Technologies). For SDS-PAGE under reducing conditions, the proteins were first treated with dithiothreitol (DTT, 25 mM) and boiled for 5 min at 100°C. SDS-PAGE gels were loaded with 1 μg of the sample and BN-PAGE gels were loaded with 4 μg of the sample. SDS-PAGE gels were run for 20 min at 250 V using SDS running buffer (Bio-Rad), and BN-PAGE gels were run for 2-2.5 h at 150 V using NativePAGE^TM^ running buffer (Life Technologies) according to the manufacturer’s instructions. SDS-PAGE gels were stained using InstantBlue (Abcam) and BN-PAGE gels were stained using Coomassie Brilliant Blue R-250 (Bio-Rad) and de-stained using a solution of 6% ethanol and 3% glacial acetic acid.

### Differential scanning calorimetry

Thermal melting curves of trimer and SApNPs following mAb148 or mAb65 purification were obtained from a MicroCal PEAQ-DSC Man instrument (Malvern). Briefly, the purified SApNP protein was buffer exchanged into 1×PBS buffer and concentrated to 0.5-3 μM prior to the analysis. Melting was probed at a scan rate of 60°C·h^−1^ from 20°C to 100°C. Data processing, including buffer correction, normalization, and baseline subtraction, was conducted using MicroCal PEAQ-DSC software. Gaussian fitting was performed using Origin 9.0 software.

### Dynamic light scattering (DLS)

Particle size distributions of hM2e and M2ex3 based on three NP platforms (FR, E2p-L4P, and I3-01v9a-L7P) were obtained from a Zetasizer Ultra instrument (Malvern). MAb148/SEC-purified NPs from ExpiCHO cells were diluted to 0.2 mg/ml using 1×PBS buffer, and 30 μl of the prepared NP sample was added to a quartz batch cuvette (Malvern, catalog no. ZEN2112). Particle size was measured at 25 °C in back scattering mode. Data processing was performed on the Zetasizer, and the particle size distribution was plotted using Origin 9.0 software.

### Negative stain EM analysis

The initial evaluation of various SApNP samples was performed by the Core Microscopy Facility at The Scripps Research Institute. All SApNPs samples were prepared at a concentration of 0.005-0.02 mg/ml. Carbon-coated copper grids (400 mesh) were glow-discharged and 8 µl of each sample was adsorbed for 2 min. Excess sample was wicked away and grids were negatively stained with 2% uranyl formate for 2 min. Excess stain was wicked away and the grids were allowed to dry. Samples were analyzed at 120 kV with a Talos L120C transmission electron microscope (Thermo Fisher) and images were acquired with a CETA 16M CMOS camera. All SApNP samples purified by mAb148 were validated under 52,000 × magnification before further use.

### Bio-layer interferometry (BLI)

The kinetics profiles of both hM2e and M2ex3 trimers and SApNPs were measured using an Octet RED96 instrument (FortéBio, Pall Life Sciences) against mAb148 and mAb65 antibody. All assays were performed with agitation set to 1000 rpm in FortéBio 1× kinetic buffer. The final volume for all solutions was 200 μl per well. Assays were performed at 30°C in solid black 96-well plates (Geiger Bio-One). For all trimers and SApNPs, 5 μg/ml antibody in 1× kinetic buffer was loaded onto the surface of anti-human Fc Capture Biosensors (AHC) for 300 s. A 60 s biosensor baseline step was applied prior to the analysis of the association of the antibody on the biosensor to the antigen in solution for 200 s. A two-fold concentration gradient of antigen, starting at 25 nM for the hM2e trimer/SApNPs and 22 nM for the M2ex3 trimer/SApNPs, was used in a titration series of six. The dissociation of the interaction was followed for 300 s. The correction of baseline drift was performed by subtracting the mean value of shifts recorded for a sensor loaded with antibody but not incubated with antigen, and for a sensor without antibody but incubated with antigen. Octet data were processed by FortéBio’s data acquisition software v.8.1. Peak signals at the highest antigen concentration were summarized in a matrix to facilitate comparisons between different vaccine platforms.

### Propagation of influenza viruses

For challenge in mice, the following reagents were obtained through BEI Resources, NIAID, NIH: Influenza A Virus, A/Puerto Rico/8/1934 Mouse-Adapted (H1N1), NR-28652 and Influenza A Virus, A/Hong Kong/1/1968-1 Mouse-Adapted 12 (H3N2), NR-28621. In brief, 4.4 × 10^6^ Madin-Darby canine kidney (MDCK) cells (CCL-34™; ATCC^®^) were plated overnight in 100 mm cell culture dishes. The next day, cells were washed with PBS and incubated with a multiplicity of infection (MOI) of 0.001 to 1 for PR8 (H1N1) or HK/68 (H3N2) in serum free media for 1 hour. Next, cells were washed and 10 ml of serum-free DMEM containing 0.2% w/v bovine serum albumin (BSA; VWR international) and 1 μg/ml L-1-tosylamido-2-phenylethyl chloromethyl ketone (TPCK)-treated trypsin (Sigma Aldrich) was added to the dishes. Cells were incubated for 65 h after which the supernatant was collected, centrifuged at 4000 RPM for 10 min, aliquoted, and frozen at -80°C until use.

### Immunoplaque assay to quantify influenza viruses

Virus PFU/ml of grown viruses was quantified using an immunoplaque assay. In brief, MDCK cells were plated in 96-well plates at 25,000 cells/well. The cells were then washed with PBS and infected with 10-fold serially diluted virus stocks. The inoculum was then removed, and cells were overlayed with 0.7% low-melt agarose (Axygen) in serum-free DMEM containing 0.2% w/v BSA and 1 μg/ml TPCK-trypsin. Twenty hours later, cells were fixed with 100 μl of 3.7% w/v formaldehyde (Sigma Aldrich) for 1 h. Cells were then permeabilized with 50 μl ice-cold methanol (Thermo Fisher) for 20 min. Fixed cells were then washed with deionized water and incubated with 50 μl of HA head-targeting IgG antibodies: FluA-20 (non-pandemic H1N1 strains), 2D1 (CA 09 H1N1), or F045-092 (pandemic or seasonal H3N2 strains) at 1 μg/ml for 1 h. Plates were washed and 50 μl of 1:2000 diluted HRP-goat anti-human IgG (Jackson ImmunoResearch Laboratories, Inc) was added to the wells. Plates were then placed on a shaker at 225 rpm for 1 h. Cells were then washed and 50 μl of TrueBlue™ Peroxidase Substrate (SeraCare) was added to wells and incubated for 10-15 min for the development of plaques. Lastly, plates were washed with deionized water and left to dry overnight. Plaques were quantified using a Bioreader^®^ 7000 (BIOSYS Scientific®).

### Mouse immunization and challenge

Six-to-eight-week-old female BALB/c mice were purchased from The Jackson Laboratory and housed in ventilated cages in environmentally controlled rooms at The Scripps Research Institute, in compliance with an approved IACUC protocol and Association for Assessment and Accreditation of Laboratory Animal Care (AAALAC) international guidelines. Institutional Animal Care and Use Committee (IACUC) guidelines were followed for all mice studies. Mice were immunized at weeks 0 and 3, with 80 μl of antigen/adjuvant mix containing 10 μg of hM2e or M2ex3 antigen in 40 μl PBS and 40 μl of adjuvant: aluminum phosphate (alum phosphate, AP) or AddaVax (AV) (both AP and AV from InvivoGen). For the hM2e study, alum phosphate was used with 10 mice/group. For the M2ex3 study, both AP and AV were evaluated with either 8 or 13 mice/group. The following reagent was obtained through BEI Resources, NIAID, NIH: Influenza A Virus, A/Puerto Rico/8/1934 (H1N1), BPL-Inactivated, NR-19325, and used as a positive control for the first strain-matched challenge (3 μg/mouse without adjuvant). Each immunization dose was split amongst four footpad ID injections (20 μl/each). To establish the lethal dose of 50% (LD_50_) in mice for PR8 (H1N1) and HK/68 (H3N2), various dilutions of grown virus stock were administered to mice (n = 7 mice/group that received 25 μl virus/nostril) after light anesthetization with isoflurane. Survival, weight loss, and morbidity were monitored for 14 days. Mice that exhibited > 25% weight loss or showed visible signs of distress were euthanized. Next, Reed-Muench and Spearman-Karber methods were used to determine the 50% endpoint titer for both PR8 (H1N1) and HK/68 (H3N2) in mice (*80, 81*). For first challenge at week 6, mice immunized with hM2e or M2ex3 constructs were lightly anesthetized with isoflurane and LD_50_ × 10 of mouse-adapted PR8 (H1N1) (50 μl) was administered to each mouse (25 μl/nostril). At week 10, surviving mice from the PR8 (H1N1) challenge were lightly anesthetized and challenged with LD_50_ × 10 of mouse-adapted HK/68 (H3N2). For the M2ex3 study, 5 mice from M2ex3 + AV groups were sacrificed on Day 5 post-PR8 (H1N1) challenge to assess the viral loads in lungs. In brief, mice were euthanized, and lungs were isolated and placed in PBS. The lung tissue was then crushed and spun down at 1200 rpm for 10 min. The lung supernatant was then aliquoted and frozen at -80°C for future analysis. Viral loads were evaluated in lung supernatant using the immunoplaque assay mentioned previously. Remaining mice were monitored for survival and weight loss for 14 days post-challenge. Post H1N1 and H3N2 challenges, mice that exhibited > 25% weight loss or showed visible signs of distress were euthanized. Blood of immunized mice was collected 2 weeks after each immunization or challenge (weeks 2, 5, 10, and 14). For the challenge after a single-dose M2ex3 immunization using potent adjuvants, mice were immunized at week 0 with 80 μl of M2ex3 I3-01v9a SApNP antigen/adjuvant mix containing 10 μg or 2 μg of M2ex3 antigen in 40 μl PBS + 40 μl of AP, 2 μg of M2ex3 antigen + 40 μg CpG (oligonucleotide 1826, a TLR9 agonist from InvivoGen), or 2 μg of M2ex3 + 40 μg STING agonist (2’3’-c-di-AM(PS)_2_(Rp,Rp), a cyclic dinucleotide from InvivoGen, identical to MIW815 (*103*)). Blood of the single-dose immunized mice was collected at week 2. To assess the protective efficacy of a single-dose immunization with potent adjuvants, mice were challenged with LD_50_ × 10 of mouse-adapted A/Puerto Rico/8/1934 (PR8) H1N1 at week 3 and monitored for weight loss for 14 days post-challenge. All bleeds were performed through the facial vein. Blood was spun down at 14,000 rpm for 10 min to separate serum from the rest of the whole blood. The serum was then heat-inactivated at 56°C for 30 min and spun down at 8,000 rpm for 10 min to remove precipitates.

### Enzyme-linked immunosorbent assay (ELISA)

For assessing hM2e-specific binding of hM2e-immune sera, 50 μl hM2e-5GS-foldon trimer probe was coated on the surface of half-well 96-well, high-binding polystyrene plates at a concentration of 0.1 μg/well. Plates were kept at 4°C overnight. The next day, plates were washed 5× with PBS containing 0.05% v/v Tween 20® (PBST). Plates were then blocked with 150 μl of 4% w/v nonfat milk (Bio-Rad) for 1 h. Plates were then washed and 50 μl of hM2e-immune sera was added to each well for 1 h. Serum was diluted 20× in 4% nonfat milk followed by seven 10-fold dilutions. M2e antibodies mAb148 and mAb65 were used as positive controls. Next, plates were washed and 50 μl of 1:3000 dilution horseradish peroxidase (HRP)-conjugated goat anti-mouse IgG in PBST was added to the wells and incubated for 1 h. Plates were then washed 6× and 50 µl of 1-Step™ 3,3’,5,5’-tetramethylbenzidine (TMB; Thermo Fisher) substrate was added to each well and incubated for 3 min. Lastly, 50 μl of 2.0 N sulfuric acid (Aqua Solutions, Inc.) was added to each well. Plates were then immediately read on a plate reader (BioTek Synergy) using a wavelength of 450 nm. An identical ELISA method was used for M2ex3-specific binding of M2ex3-immune sera, except for the use of M2ex3-5GS-foldon trimer probe as the coating antigen.

### Cell-based ELISA

For cell-based ELISA, the following 8 reagents were obtained through BEI Resources, NIAID, NIH: 1) Influenza A Virus, A/Puerto Rico/8/1934 (H1N1; NR-348), 2) Influenza A Virus, A/California/04/2009 (H1N1; NR-136583), 3) Influenza A Virus, A/Solomon Islands/3/2006 (H1N1; NR-41798), 4) Influenza A Virus, A/Hong Kong/1/1968 (H3N2) (mother clone),, NR-28620, 5) Influenza A virus, A/Brisbane/10/2007 (H3N2; NR-12283, 6) Influenza A virus, A/Aichi/2/1968 (H3N2; NR-3177), 7) Influenza B Virus, B/Florida/4/2006 (Yamagata Lineage; NR-41795), and 8) Influenza B Virus, B/Brisbane/60/2008 (Victoria Lineage; NR-42005). The viruses were grown in MDCK cells using the same method mentioned previously for propagating and quantifying challenge strains. For cell-based infection ELISA, MDCK cells were plated overnight in 96-well cell culture plates at a density of 18,000 cells/well. The next day the cells were washed and infected with 100 μl of 1 of the 8 viruses at a MOI of 0.1. Twenty hours later, the virus was removed, and cells were washed before being fixed with 100 μl of 3.7% w/v formaldehyde. Cells were then washed, and the previous ELISA protocol was used except with an incubation step with TMB for 5 min.

### Histology, immunostaining, and imaging

To study vaccine distribution, trafficking, retention, cellular interaction, and GC reactions in lymph nodes, M2ex3 trimer and FR and I3-01v9a SApNP immunogens formulated with AV adjuvant were injected intradermally into four mouse footpads using 29-gauge insulin needles. Mice were anesthetized with 3% isoflurane in oxygen during immunization. Similar protocols of mouse injection, lymph node collection and tissue analysis were utilized from our previous study (*56, 74*). The injection dose was 80 μl of antigen/adjuvant mix containing 40 μg of vaccine immunogen per mouse or 10 μg per footpad. Mice were euthanized at 2 h to 8 weeks after a single-dose injection or 2 and 5 weeks after the boost, which occurred at 3 weeks after the first dose. Brachial and popliteal sentinel lymph nodes were collected for immunohistological study. Fresh lymph nodes were isolated and merged into frozen section compound (VWR International, catalog no. 95057-838) in a plastic cryomold (Tissue-Tek at VWR, catalog no. 4565). Tissue samples were frozen in liquid nitrogen and stored at -80°C before shipping to The Centre for Phenogenomics in Canada for tissue processing, immunostaining, and imaging. Lymph node tissue sections were sliced 8 μm thick on a cryostat (Cryostar NX70) and placed on a charge slide. Next, tissue slides were fixed in 10% neutral buffered formalin and permeabilized in PBS buffer that contained 0.5% Triton X-100 before immunostaining. The slides were blocked with Protein Block (Agilent) to prevent nonspecific antibody binding. Primary antibodies were applied on tissue slides and incubated overnight at 4°C. After washing with tris-buffered saline with 0.1% Tween-20 (TBST), biotin or fluorophore-conjugated secondary antibodies were applied and incubated at 25°C for 1 hour. Lymph node tissues were stained with anti-human Ab148 or Ab65 (1:200), and biotinylated goat anti-human secondary antibody (Abcam, catalog no. ab7152, 1:300), followed by streptavidin-horseradish peroxidase (HRP) reagent (Vectastain Elite ABC-HRP Kit, Vector, catalog no. PK-6100) and diaminobenzidine (ImmPACT DAB, Vector, catalog no. SK-4105).

To study cellular interactions between M2ex3 trimer immunogens and cell components in lymph nodes, FDCs were labelled using anti-CD21 primary antibody (Abcam, catalog no. ab75985, 1:1800), followed by anti-rabbit secondary antibody conjugated with Alexa Fluor 555 (Thermo Fisher, catalog no. A21428, 1:200). Subcapsular sinus macrophages were labeled using anti-sialoadhesin (CD169) antibody (Abcam, catalog no. ab53443, 1:600), followed by anti-rat secondary antibody conjugated with Alexa Fluor 488 (Abcam, catalog no. ab150165, 1:200). B cells were labeled using anti-B220 antibody (eBioscience, catalog no. 14-0452-82, 1:100), followed by anti-rat secondary antibody conjugated with Alexa Fluor 647 (Thermo Fisher, catalog no. A21247, 1:200). GC reactions induced by M2ex3 trimers and SApNPs were studied by immunostaining. GC B cells were labeled using rat anti-GL7 antibody (FITC; BioLegend, catalog no. 144604, 1:250). T_fh_ cells were labeled using anti-CD4 antibody (BioLegend, catalog no. 100402, 1:100), followed by anti-rat secondary antibody conjugated with Alexa Fluor 488 (Abcam, catalog no. ab150165, 1:1000). GC forming cells were stained using Bcl6 antibody (Abcam, catalog no. ab220092, 1:300), followed by anti-rabbit secondary antibody conjugated with Alexa Fluor 555 (Thermo Fisher, catalog no. A21428, 1:1000). Nuclei were labeled using 4′,6-diamidino-2-phenylindole (DAPI) (Sigma-Aldrich, catalog no. D9542, 100 ng/ml). The immunostained lymph node tissues were scanned using an Olympus VS-120 slide scanner with a Hamamatsu ORCA-R2 C10600 digital camera. The vaccine transport and induced GC reactions in lymph nodes were quantified through bright-field and fluorescent images using ImageJ software.

### Lymph node disaggregation, cell staining, and flow cytometry

GC reactions in terms of frequency and numbers of GC B cells (GL7^+^B220^+^) and T_fh_ cells (CD3^+^CD4^+^CXCR5^+^PD-1^+^) were characterized using flow cytometry (**Figure S9**). Mice were euthanized at 2, 5, and 8 weeks after a single-dose injection or 2 and 5 weeks after the boost, occurring at 3 weeks after the first dose (4 footpads, 10 μg/footpad). Fresh axillary, brachial, and popliteal sentinel lymph nodes were collected for GC study. Lymph node tissues were disaggregated mechanically and merged in enzyme digestion solution in an Eppendorf tube with 958 μl of Hanks’ balanced salt solution (HBSS) buffer (Thermo Fisher Scientific, catalog no. 14185052), 40 μl of 10 mg/ml collagenase IV (Sigma-Aldrich, catalog no. C5138), and 2 μl of 10 mg/ml DNase (Roche, catalog no. 10104159001) and incubated at 37°C for 30 min. The lymph node tissue was then filtered through a 70 μm cell strainer to obtain a single cell suspension. Cell samples were spun down at 400 × *g* for 10 min and the cell pellets were resuspended in HBSS blocking buffer containing 0.5% (w/v) bovine serum albumin and 2 mM EDTA. Anti-CD16/32 antibody (BioLegend, catalog no. 101302) was added into the Eppendorf tube to block the nonspecific binding on Fc receptors. Cells were kept on ice for 30 min and transferred to 96-well V-shaped-bottom microplates with pre-prepared cocktail antibodies, including Zombie NIR live/dead stain (BioLegend, catalog no. 423106), Brilliant Violet 510 anti-mouse/human CD45R/B220 antibody (BioLegend, catalog no. 103247), FITC anti-mouse CD3 antibody (BioLegend, catalog no. 100204), Alexa Fluor 700 anti-mouse CD4 antibody (BioLegend, catalog no. 100536), PE anti-mouse/human GL7 antibody (BioLegend, catalog no. 144608), Brilliant Violet 605 anti-mouse CD95 (Fas) antibody (BioLegend, catalog no. 152612), Brilliant Violet 421 anti-mouse CD185 (CXCR5) antibody (BioLegend, catalog no. 145511), and PE/Cyanine7 anti-mouse CD279 (PD-1) antibody (BioLegend, catalog no. 135216). Cells were mixed with antibody cocktail and placed on ice for 30 min. Cell samples were spun down at 400 × *g* for 10 min and the cell pellets were resuspended in HBSS blocking solution for washing one more time. Cells were then fixed with 1.6% paraformaldehyde (Thermo Fisher Scientific, catalog no. 28906) in HBSS and placed on ice for 30 min. Cells were spun down at 400 × *g* for 10 min and placed in HBSS blocking buffer at 4°C before test. Sample events were acquired using a 5-laser AZE5 flow cytometer (Yeti, Bio-Rad) with Everest software at the Core Facility of The Scripps Research Institute. The data were analyzed using FlowJo 10 software.

### Antibody-dependent cell cytotoxicity (ADCC) surrogate assay

The potential for M2e-specific antibodies to induce ADCC was evaluated using a mouse FcγRIV ADCC Reporter kit (Promega). In brief, MDCK cells were plated in white 96-well plates overnight at 18,000 cells/well. The next day, cells were washed with PBS and infected with PR8 (H1N1) at a MOI of 0.1. Twenty hours later, the cells were washed and 25 μl of M2ex3-immune sera was added to the wells for 1 h (sera was diluted 20× followed by 10-fold dilutions). Next, as per the kit’s instructions (Promega), 75,000 mouse Fcγ receptor IV (mFcγRIV)-expressing Jurkat effector cells were added to each well to engage with the Fc region of M2e serum antibodies bound to native M2e expressed on the surface of the infected cells, resulting in nuclear factor of activated T cells (NFAT)-mediated luciferase activity. The plates were incubated for 6 h at 37°C. Lastly, 75 μl of Bio-Glo™ Reagent were added to the well. Relative light units were measured using a plate reader after a 5-min incubation of each plate.

### Splenocyte isolation

At week 14, mice were anesthetized using isoflurane and euthanized using cervical dislocation. Spleens were harvested from mice and kept in PBS on ice. Next, spleens were crushed with the back of a syringe and resuspended in 20 ml of PBS. Cells were centrifuged at 1200 rpm for 10 min. Next, supernatant was discarded, and 3 ml of ACK lysis buffer (Lonza) was added to the cells and incubated for 5 min. Next 12 ml of PBS was added to the tubes, which were then centrifuged at 1200 rpm for 5 min. Supernatant was then discarded, and cells were resuspended in 1 ml PBS. Cells were passed through a 70-μm cell strainer. Cells were then centrifuged for 5 min. Lastly, cells were resuspended in 10% DMSO in FBS, transferred into a -80°C freezer overnight, and then stored in the vapor phase of liquid nitrogen until analysis.

### Enzyme-linked immunosorbent spot (ELISpot) assay

For analyzing IFN-γ and IL-4-secreting splenocytes of mice immunized with M2ex3, ELISpot was used. First, Multiscreen® filter plates (Millipore Sigma) were coated with capture IFN-γ or IL-4 (BD Biosciences) at a 1:200 dilution. The plates were incubated at 4 °C overnight. The next day, plates were washed and blocked with 200 μl of complete RPMI 1640 (10% FBS, 1% Penn-Strep, 1% L-glutamine; Gibco) medium. Next, the M2ex3-5GS-foldon trimer probe was prepared at 50 μg/ml in complete RPMI. Concanavalin A (10 μg/ml; BD Biosciences) was prepared as a positive control. Next, spleen samples were thawed, resuspended in RPMI and centrifuged at 1200 RPM for 10 min. Cells were then counted using an automated cell counter (Countess II; Thermo Fisher) and suspended to reach a concentration of 1.6 × 10^7^ cells/ml. Next, RPMI was discarded from the filter plates and 50 μl of antigen was added to the well. Next, 50 μl of cell suspension was added to each well, producing a final cell concentration of 8 × 10^5^ splenocytes/well. After addition of cells, the final concentrations of the M2ex3-5GS-foldon trimer probe and Concanavalin A were 2.5 μg/well and 0.5 μg/well, respectively. Cells without antigen were used as a negative control. Plates were incubated for 48 h at 37°C. Next, cells were washed 2× with deionized water followed by 3 washes with PBST. Next, 50 μl detection IFN-γ or IL-4 antibodies (BD Biosciences) diluted 1:250 in dilution buffer (10% FBS in PBS) were added to corresponding wells. Plates were then incubated for 2 h at room temperature (RT). Next, plates were washed 3× with PBST and 50 μl Streptavidin-HRP (BD Biosciences) diluted 1:100 in dilution buffer was added to the wells and incubated for 1 h. Plates were then washed 4× with PBST and 2× with PBS. AEC Final Substrate (BD Biosciences) was then added to the wells for 15-20 min for the development of spots. Plates were kept in the dark overnight and read using a Bioreader^®^ 7000.

### T cell culture and stimulation

Functional M2e-specific T cell responses were characterized by measuring activation induced marker (AIM) and intracellular cytokine staining (ICS) using flow cytometry (**Figure S10**). Mouse splenocytes were isolated from naïve or vaccinated mice at 5 days after prime-boost immunizations and followed by a H1N1 virus challenge. Cryopreserved splenocytes were thawed by diluting cells in 10 ml pre-warmed complete RPMI media with 10% deactivated fetal bovine serum (FBS) and 1% penicillin/streptomycin (P/S). Cells were spun down at 400 × *g* for 10 min and cell pellets were resuspended in RPMI media. The numbers of splenocytes were counted and adjusted to 10 million cells/ml. One million splenocytes for each mouse were placed into 96-well U-shaped-bottom microplates. Cells were cultured in the presence of the M2ex3-5GS-foldon trimer probe (1 μg per well) at 37°C for a total of 24 h. After 20 h, Brefeldin A Solution (BioLegend, catalog no. 420601) was added in the culture to enhance intracellular cytokine staining signals by inhibiting the protein transport processes in the rough endoplasmic reticulum and Golgi apparatus. After 4 h, cells were then spun down at 400 × *g* for 10 min and cell pellets were resuspended in HBSS blocking buffer. Anti-CD16/32 antibody (BioLegend, catalog no. 101302) was added for 30 min on ice to block nonspecific binding to Fc receptors. Cells were then transferred to 96-well V-shaped-bottom microplates with pre-prepared cocktail antibodies for surface marker staining, including LIVE/DEAD Fixable Blue Dead Cell Stain Kit (Thermo Fisher Scientific, catalog no. L34962), FITC anti-mouse CD3 antibody (BioLegend, catalog no. 100204), Alexa Fluor 700 anti-mouse CD4 antibody (BioLegend, catalog no. 100536), BUV737 Anti-Mouse CD8a antibody (BD Bioscience, catalog no. 612759), APC anti-mouse CD154 antibody (BioLegend, catalog no. 106510), Brilliant Violet 421 anti-mouse CD69 Antibody (BioLegend, catalog no. 104527), APC/Fire 810 anti-mouse CD279 (PD-1) antibody (BioLegend, catalog no. 135251), and Brilliant Violet 605 anti-mouse CD185 (CXCR5) Antibody (BioLegend, catalog no. 145513). Splenocytes were mixed with antibody cocktail and placed on ice for 30 min. Cells were spun down at 400 × *g* for 10 min and the cell pellets were resuspended in HBSS blocking solution, then washed once more. Cells were fixed with 1.6% paraformaldehyde (Thermo Fisher Scientific, catalog no. 28906) in HBSS and placed on ice for 30 min. Cell were then washed two times with intracellular staining permeabilization wash buffer (BioLegend, catalog no. 421002) and stained with previously prepared antibody cocktail for intracellular staining, including PE anti-mouse IFN-γ antibody (BioLegend, catalog no. 505808), Brilliant Violet 785 anti-mouse TNF-α antibody (BioLegend, catalog no. 506341), PE/Cyanine5 anti-mouse IL-2 antibody (BioLegend, catalog no. 503824), PE/Cyanine7 anti-mouse IL-4 antibody (BioLegend, catalog no. 504118), and Brilliant Violet 711 anti-mouse IL-17A Antibody (BioLegend, catalog no. 506941). Cells were mixed with intracellular antibodies and placed on ice for 30 min. Cells were then washed again with intracellular staining permeabilization wash buffer. Cells were stored in HBSS blocking buffer at 4°C prior to analysis. Sample events were acquired using a Cytek Aurora spectral analytical flow cytometer with SpectroFlo software at the Flow Cytometry Core Facility of The Scripps Research Institute. The data were analyzed using FlowJo 10 software.

### Statistical analysis

Data was collected from 7-13 mice per group in the immunization studies, challenge experiments, and serum binding assays. For the vaccine transport and GC study in lymph nodes and T cells in spleens, 5 mice per group with different vaccine constructs were compared using one-way ANOVA, followed by Tukey’s multiple comparison *post hoc* test. Statistical significance is indicated as the following in the figures: ns (not significant), \**p* < 0.05, \*\**p* < 0.01, \*\*\**p* < 0.001, \*\*\*\**p* < 0.0001. The graphs were generated using GraphPad Prism 9.3.1 software.

## SUPPLEMENTARY MATERIALS

Supplementary material for this article is available at XXX. Supplementary Figures 1-11.

## ACKNOWLEDGEMENTS

Y.-N.Z. thanks support from the Natural Sciences and Engineering Research Council of Canada (NSERC) for the postdoctoral fellowship. We acknowledge G. Ossetchkine, K. Duffin, M. Ganguly, and V. Bradaschia at The Centre for Phenogenomics for their expertise and technical support in immunohistology. We acknowledge A. Saluk, B. Seegers, and B. Monteverde of the Flow Cytometry Core Facility at The Scripps Research Institute for their technical support in flow cytometry. We thank V. Tong for proofreading the manuscript.

## Author contributions

Project design by K.B.G., Y.-N.Z., Y.-Z.L., L.H. and J.Z.; immunogen expression and purification by M.E., A.L., G.W., and L.H.; negative stain EM by Y.-Z.L. and L.H.; mouse immunization and sample collection by K.B.G. and Y.-N.Z.; virus challenge and serum binding analysis by K.B.G.; ELISpot and ADCC by K.B.G.; lymph node isolation, immunohistology, and flow cytometry by Y.-N.Z.; manuscript written by K.B.G., Y.-N.Z., Y.-Z.L., S.A., L.H. and J.Z. All authors were asked to comment on the manuscript. The Scripps Research Institute manuscript number is 30235.

## Data and material availability

All data to understand and assess the conclusions of this research are available in the main text and Supplementary Materials.

## SUPPLEMENTAL LEGENDS

**Figure S1.** Schematic representation of computational design for I3-01v9a.

**Figure S2.** Design and in vitro characterization of hM2e-presenting SApNPs.

**Figure S3.** Serum binding of individual mice immunized with hM2e 1TD0 and SApNPs.

**Figure S4.** Design and in vitro characterization of M2ex3-presenting SApNPs.

**Figure S5.** Serum binding of individual mice immunized with M2ex3 1TD0 and SApNPs.

**Figure S6.** M2ex3 serum binding of individual mice to cell-surface homotetrameric M2e.

**Figure S7.** Immunohistological images of M2ex3 1TD0 and SApNPs in lymph nodes.

**Figure S8.** Immunohistological analysis of M2ex3 1TD0 and SApNP vaccine-induced GCs.

**Figure S9.** Flow cytometry analysis of M2ex3 1TD0 and SApNP vaccine-induced GCs.

**Figure S10.** Flow cytometry analysis of M2ex3 1TD0 and SApNP vaccine-induced T cell responses.

**Figure S11.** Serum binding of individual mice immunized with SApNPs and various adjuvants.

