## Supplemental Material for "Single-component multilayered self-assembling protein nanoparticles displaying extracellular domains of matrix protein 2 as a pan-influenza A vaccine"

**Figure S1**

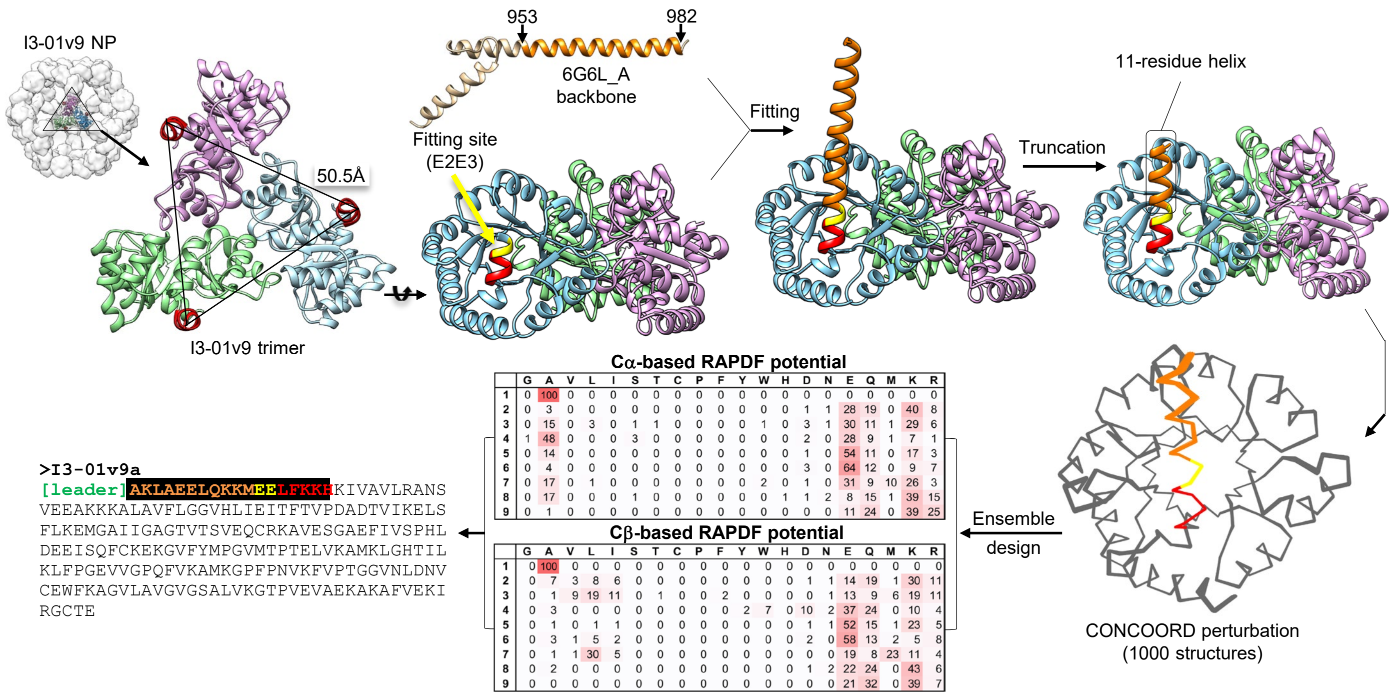


**Cα-based RAPDF potential**

| | G | A | V | L | I | S | T | C | P | F | Y | W | H | D | N | E | Q | M | K | R |
|---|---|---|---|---|---|---|---|---|---|---|---|---|---|---|---|---|---|---|---|---|
| **1** | 0 | 100 | 0 | 0 | 0 | 0 | 0 | 0 | 0 | 0 | 0 | 0 | 0 | 0 | 0 | 0 | 0 | 0 | 0 | 0 |
| **2** | 0 | 3 | 0 | 0 | 0 | 0 | 0 | 0 | 0 | 0 | 0 | 0 | 0 | 1 | 1 | 28 | 19 | 0 | 40 | 8 |
| **3** | 0 | 15 | 0 | 3 | 0 | 1 | 1 | 0 | 0 | 0 | 0 | 1 | 0 | 3 | 1 | 30 | 11 | 1 | 29 | 6 |
| **4** | 1 | 48 | 0 | 0 | 0 | 3 | 0 | 0 | 0 | 0 | 0 | 0 | 0 | 2 | 0 | 28 | 9 | 1 | 7 | 1 |
| **5** | 0 | 14 | 0 | 0 | 0 | 0 | 0 | 0 | 0 | 0 | 0 | 0 | 0 | 1 | 0 | 54 | 11 | 0 | 17 | 3 |
| **6** | 0 | 4 | 0 | 0 | 0 | 0 | 0 | 0 | 0 | 0 | 0 | 0 | 0 | 3 | 0 | 64 | 12 | 0 | 9 | 7 |
| **7** | 0 | 17 | 0 | 1 | 0 | 0 | 0 | 0 | 0 | 0 | 0 | 2 | 0 | 1 | 0 | 31 | 9 | 10 | 26 | 3 |
| **8** | 0 | 17 | 0 | 0 | 0 | 1 | 0 | 0 | 0 | 0 | 0 | 0 | 1 | 1 | 2 | 8 | 15 | 1 | 39 | 15 |
| **9** | 0 | 1 | 0 | 0 | 0 | 0 | 0 | 0 | 0 | 0 | 0 | 0 | 0 | 0 | 0 | 11 | 24 | 0 | 39 | 25 |

**Cβ-based RAPDF potential**

| | G | A | V | L | I | S | T | C | P | F | Y | W | H | D | N | E | Q | M | K | R |
|---|---|---|---|---|---|---|---|---|---|---|---|---|---|---|---|---|---|---|---|---|
| **1** | 0 | 100 | 0 | 0 | 0 | 0 | 0 | 0 | 0 | 0 | 0 | 0 | 0 | 0 | 0 | 0 | 0 | 0 | 0 | 0 |
| **2** | 0 | 7 | 3 | 8 | 6 | 0 | 0 | 0 | 0 | 0 | 0 | 0 | 0 | 1 | 1 | 14 | 19 | 1 | 30 | 11 |
| **3** | 0 | 1 | 9 | 19 | 11 | 0 | 1 | 0 | 0 | 2 | 0 | 0 | 0 | 0 | 1 | 13 | 9 | 6 | 19 | 11 |
| **4** | 0 | 3 | 0 | 0 | 0 | 0 | 0 | 0 | 0 | 0 | 2 | 7 | 0 | 10 | 2 | 37 | 24 | 0 | 10 | 4 |
| **5** | 0 | 1 | 0 | 1 | 1 | 0 | 0 | 0 | 0 | 0 | 0 | 0 | 0 | 1 | 1 | 52 | 15 | 1 | 23 | 5 |
| **6** | 0 | 3 | 1 | 5 | 2 | 0 | 0 | 0 | 0 | 0 | 0 | 0 | 0 | 2 | 0 | 58 | 13 | 2 | 5 | 8 |
| **7** | 0 | 1 | 1 | 30 | 5 | 0 | 0 | 0 | 0 | 0 | 0 | 0 | 0 | 0 | 0 | 19 | 8 | 23 | 11 | 4 |
| **8** | 0 | 2 | 0 | 0 | 0 | 0 | 0 | 0 | 0 | 0 | 0 | 0 | 0 | 1 | 2 | 22 | 24 | 0 | 43 | 6 |
| **9** | 0 | 0 | 0 | 0 | 0 | 0 | 0 | 0 | 0 | 0 | 0 | 0 | 0 | 0 | 0 | 21 | 32 | 0 | 39 | 7 |

**Figure S1. Schematic representation of computational design for I3-01v9a.** Top left: The molecular surface model (gray) of I3-01v9 NP and a zoomed-in view of the ribbon model of an I3-01v9 trimer (chains A, B, and C colored in sky blue, plum, and light green, respectively), with the N-termini forming a triangle of 50.5 Å. Also shown is the ribbon model of a c-MYC transcription factor protein (PDB ID: 6G6L), from which the backbone of a helix (residues 953-982, orange) was grafted onto an I3-01v9 subunit by using residues E2 and E3 (yellow) of I3-01v9 for fitting. Top right: A side view of the ribbon model of an I3-01v9 trimer with the fitted full helix backbone (30 residues) and truncated helix backbone (11 residues). Bottom right: 1,000 slightly perturbed backbone conformations of the modified I3-01v9 subunit with an extended N-terminal helix generated using CONCOORD, a protein structure sampling program. Bottom middle: Predicted amino acid (frequency, %) for each position of the first 9 residues using Cα and Cβ-based RAPDF scoring functions. Bottom left: The final sequence design, termed I3-01v9a, with the predicted residues of the N-terminal helical extension colored in orange. The anchoring residues E2 and E3 are colored in yellow, and the last remaining turn of the N-terminal helix is colored in red.

### Figure S2

A

**>hM2e-5GS-1TD0**

DAMKRGLCCVLLLCGAVFVSPSQEIHARFRRGARSRSLLTEVETPIRNEWGCRCNDSSDSASGGGGSEVRIFAGNDPAHTATGSSGISSPTPALTPLMLDEATGKLVVWDGQKAGSAVGILVLPLEGTETALTYYKS
GTFATEAIHWPESVDEHKKANAFAGSALSHAA

**>hM2e-5GS-FR**

DAMKRGLCCVLLLCGAVFVSPSQEIHARFRRGARSRSLLTEVETPIRNEWGCRCNDSSDASGGGGSASGDIIKLLNEQVNKEMQSSNLYMSMSSWCYTHSLDGAGLFLFDHAAEEYEHAKKLIIFLNENNVPVQLTS
ISAPEHKFEGLTQIFQKAYEHEQHISESINNIVDHAIKSKDHATFNFLQWYVAEQHEEEVLFKDILDKIELIGNENHGLYLAD

**>hM2e-5GS-E2p-LD4-PADRE(hM2e-E2p-L4P)**

DAMKRGLCCVLLLCGAVFVSPSQEIHARFRRGARSRSLLTEVETPIRNEWGCRCNDSSDASGGGGSGAAAKPATTEGEFPETREKMSGIRRAIAKAMVHSKHTAPHVTLMDEADVTKLVAHRKKFKAIAAEKGIKLT
FLPYVVKALVSALREYPVLNTAIDDETEEIIQKHYYNIGIAADTDRGLLVPVIKHADRKPIFALAQEINELAEKARDGKLTPGEMKGASCTITNIGSAGGQWFTPVINHPEVAILGIGRIAEKPIVRDGEIVAAPML
ALSLSFDHRMIDGATAQKALNHIKRLLSDPELLLMGGGGSFSEEQKKALDLAFYFDRRLTPEWRRYLSQRLGLNEEQIERWFRRKEQQIGWSHPQFEKGSAKFVAAWTLKAAA

**>hM2e-5GS-I3-01v9a-LD7-PADRE(hM2e-I3-01v9a-L7P)**

DAMKRGLCCVLLLCGAVFVSPSQEIHARFRRGARSRSLLTEVETPIRNEWGCRCNDSSDASGGGGSGAKLAEELQKKMEELFKKHKIVAVLRANSVEEAKEKALAVFEGGVHLIEITFTVPDADTVIKELSFLKEKG
AIIGAGTVTSVEQCRKAVESGAEFIVSPHLDAEITVFCLEKGVFYMPGVMTPTELVKAMKLGHNILKLFPGEVVGPQFVKAMKGPFPNVKFVPTGGVNLDNVCEWFKAGVLAVGVGSALVKGTPDEVREKAKAFVEK
IRGCTEGGGGSSPAVDIGDRLDELEKALEALSAEDGHDDVGQRLESLLRRWNSRRADGSAKFVAAWTLKAAA

B

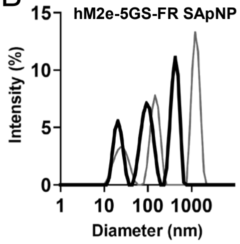


C

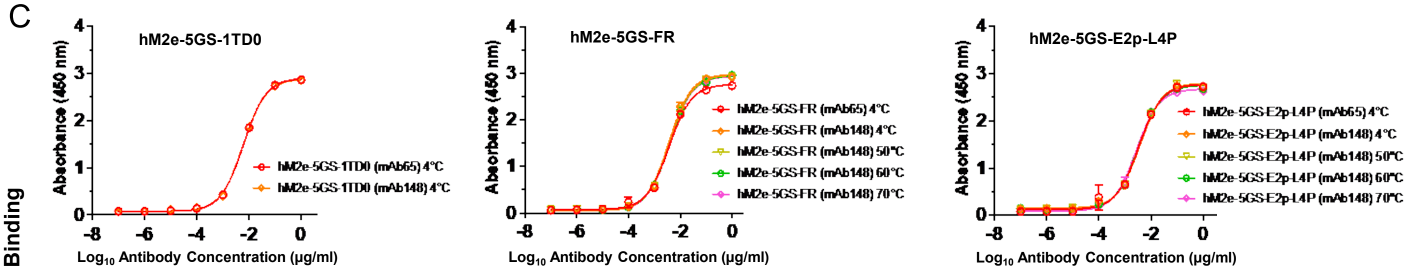


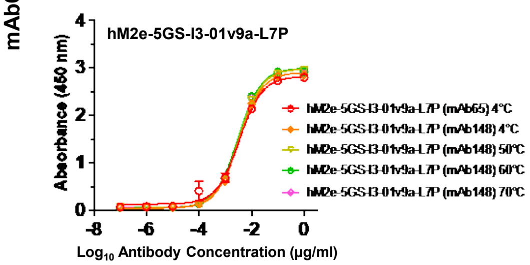


| hM2e | Primary Antibody Binding $EC_{50}$ (μg/ml) | |
|---|---|---|
| Groups | mAb65 | mAb148 |
| hM2e-5GS-1TD0 4°C | 0.006025 | 0.01972 |
| hM2e-5GS-FR 4°C | 0.004023 | 0.01038 |
| hM2e-5GS-FR 50°C | 0.003854 | 0.007114 |
| hM2e-5GS-FR 60°C | 0.003853 | 0.007796 |
| hM2e-5GS-FR 70°C | 0.003909 | 0.007134 |
| hM2e-5GS-E2p-L4P 4°C | 0.003703 | 0.01737 |
| hM2e-5GS-E2p-L4P 50°C | 0.003565 | 0.01721 |
| hM2e-5GS-E2p-L4P 60°C | 0.003391 | 0.01685 |
| hM2e-5GS-E2p-L4P 70°C | 0.002789 | 0.01477 |
| hM2e-5GS-I3-01v9a-L7P 4°C | 0.003615 | 0.01066 |
| hM2e-5GS-I3-01v9a-L7P 50°C | 0.003386 | 0.01315 |
| hM2e-5GS-I3-01v9a-L7P 60°C | 0.003107 | 0.01467 |
| hM2e-5GS-I3-01v9a-L7P 70°C | 0.003153 | 0.0145 |

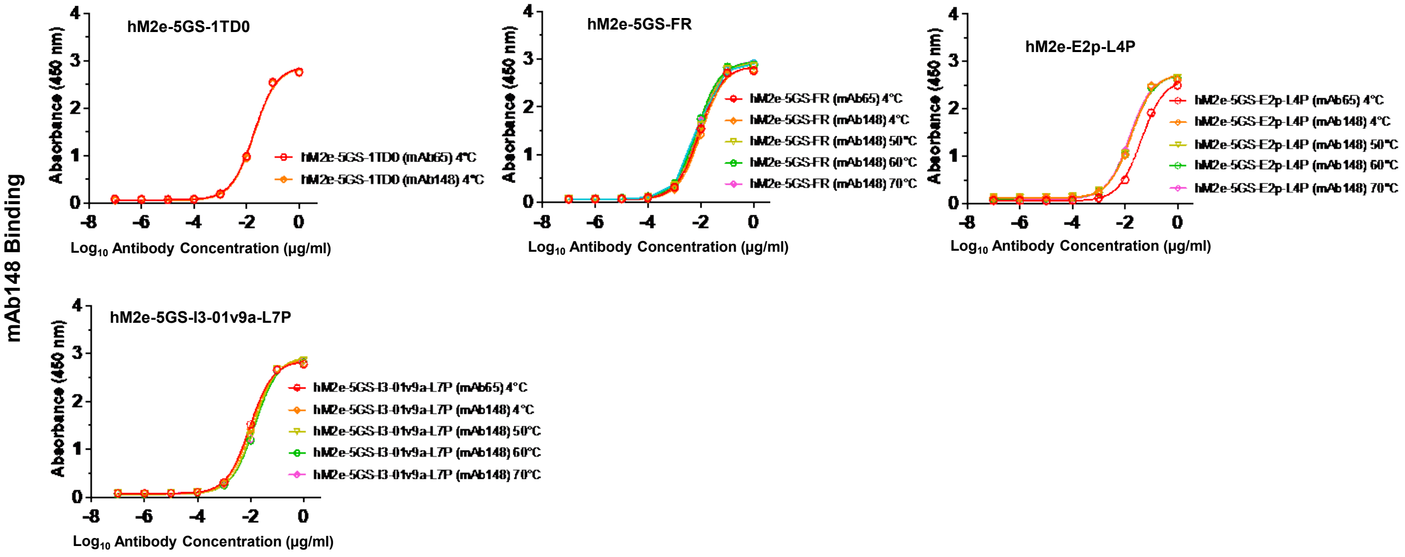

D

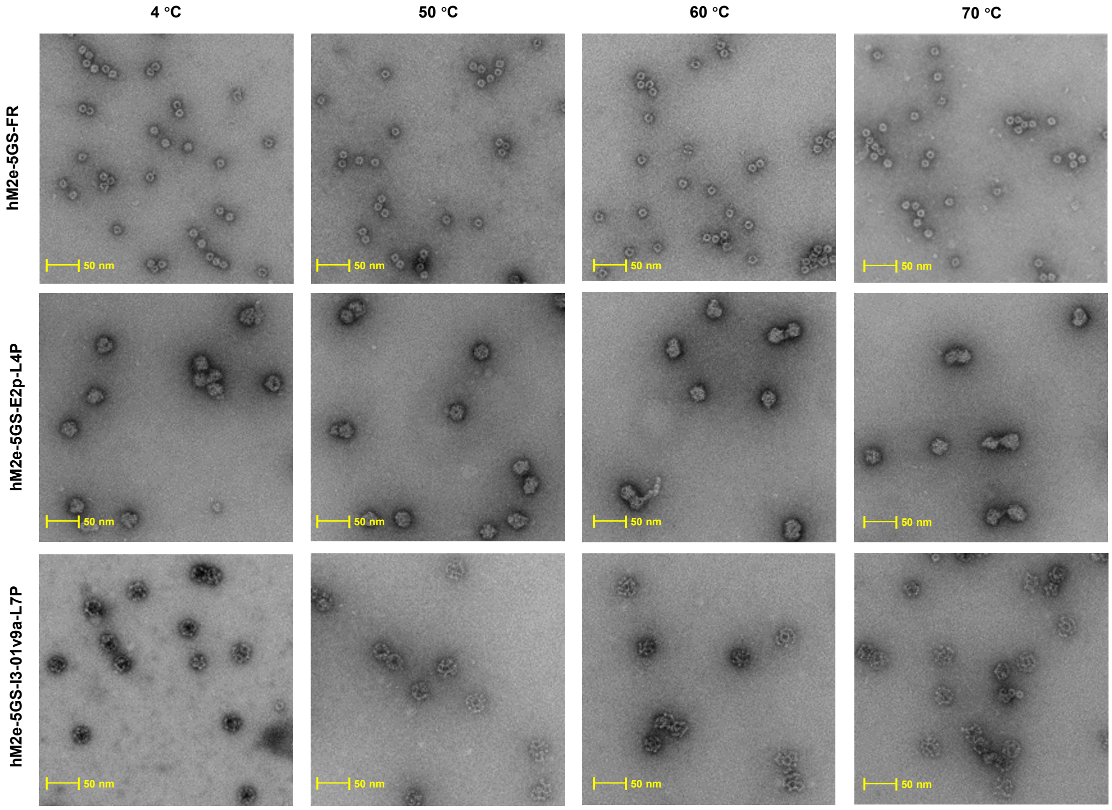


E

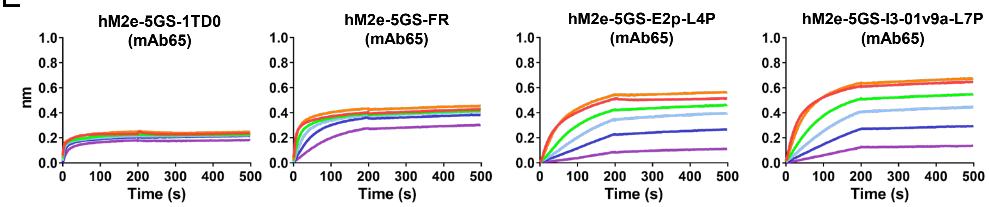


**Figure S2. Design and in vitro characterization of hM2e-presenting SApNPs.** (**A**) Construct sequences of hM2e-presenting FR, E2p-L4P, and I3-01v9-L7P SApNPs, with the gene fragments of leader sequence, restriction site, human Matrix protein 2 extra-virion domain (residue: 2-24), flexible linker, NP-forming subunit, locking domain (LD), and PADRE highlighted in yellow, green, gray, light magenta, cyan, olive green, and red shades, respectively. (**B**) Dynamic Light Scatter results for hM2e-FR after SEC purification. (**C**) ELISA curves of for hM2e-presenting SApNPs binding to mAb148 and mAb65 antibody. (**D**) negative stain EM image for hM2e-presenting SApNPs. (**E**) Antigenic evaluation of hM2e-presenting SApNPs using BLI for mAb65 antibody. A two-fold concentration gradient of antigen, starting at 5.0 μM for hM2e 1TD0 trimer, 80.0 nM for FR-SApNP, and 20.0 nM for E2p and I3-01v9a SApNPs, was used in a titration series of six.

#### Figure S3

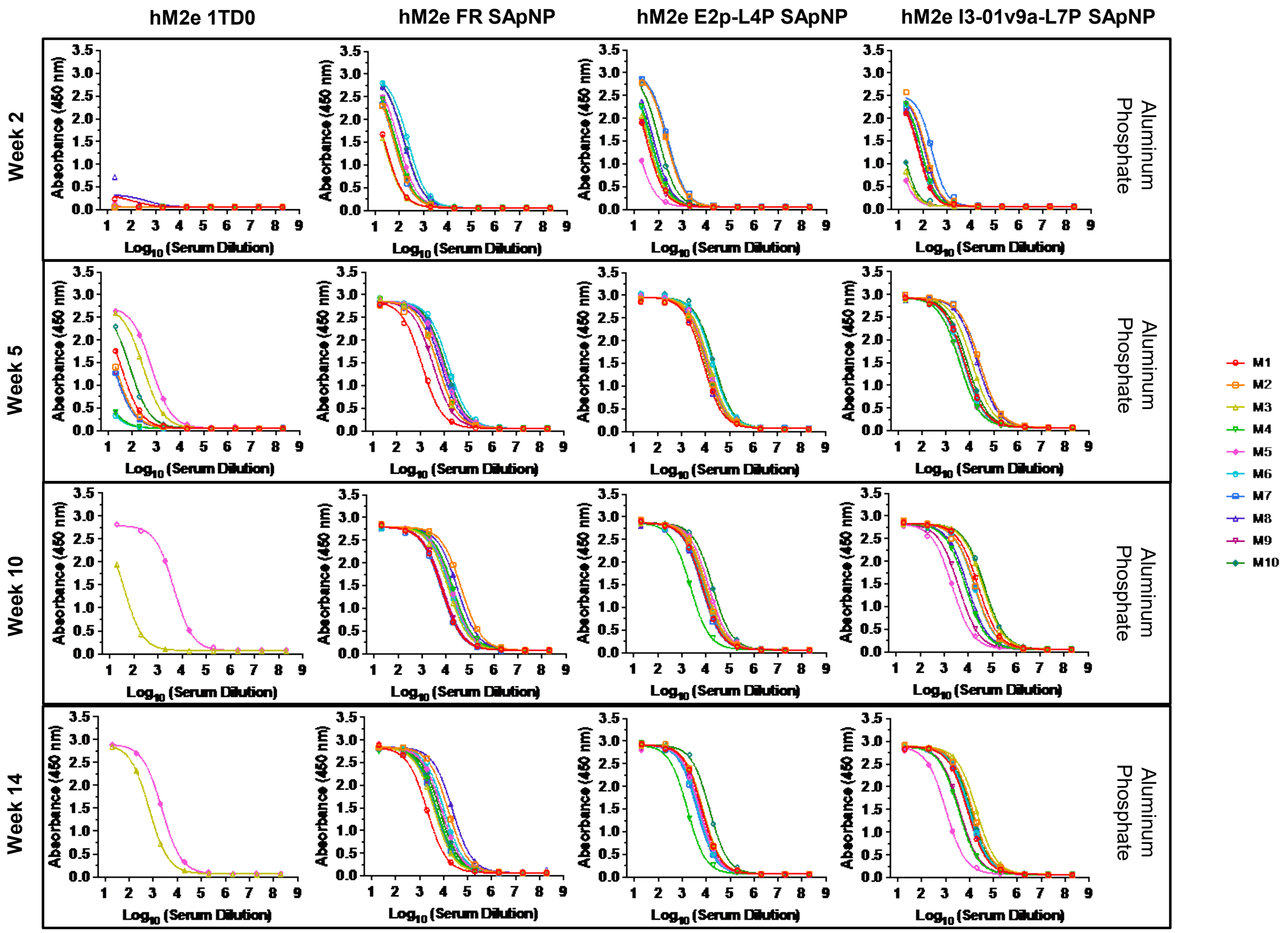


**Figure S3. Serum binding of individual mice immunized with hM2e 1TD0 and SApNPs.** ELISA curves showing hM2e-immune sera (adjuvanted with aluminum phosphate) binding to the hM2e-5GS-foldon trimer probe and calculated 50% effective concentration ($EC_{50}$) values for weeks 2 and 5 (N = 10). N = variable at weeks 10 and 14 based on surviving mice/group post-H1N1 and H3N2 challenges at weeks 6 and 10, respectively. Serum was tested was from mice immunized with hM2e 1TD0 trimer, and FR, E2p, and I3-01v9a SApNP at weeks 0 and 3. The assay was performed in duplicate with a starting serum dilution of 20× followed by seven 10-fold titrations.

#### Figure S4

A

>M2ex3-5GS-1TD0

DAMKRGLCCVLLLCGAVFVSPSQEIHARFRRGARSRSLLTEVETPIRNEWGCRCNDSSDGGGGSLLTEVETPTRNGWESKSSDSSDGGGGSLLTEVETPTRSEWESRSSGSSDASGGGGSEVRIFAGNDPAHTATGS
SGISSPTPALTPLMLDEATGKLVVWDGQKAGSAVGILVLPLEGTETALTYYKSGTFATEAIHWPESVDEHKKANAFAGSALSHAA

>M2ex3-5GS-FR

DAMKRGLCCVLLLCGAVFVSPSQEIHARFRRGARSRSLLTEVETPIRNEWGCRCNDSSDGGGGSLLTEVETPTRNGWESKSSDSSDGGGGSLLTEVETPTRSEWESRSSGSSDASGGGGSGDIIKLLNEQVNKEMQS
SNLYMSMSSWCYTHSLDGAGLFLFDHAAEEYEHAKKLIIFLNENNVPVQLTSISAPEHKFEGLTQIFQKAYEHEQHISESINNIVDHAIKSKDHATFNFLQWYVAEQHEEEVLFKDILDKIELIGNENHGLYLAD

>M2ex3-5GS-E2p-LD4-PADRE (M2ex3-E2p-L4P)

DAMKRGLCCVLLLCGAVFVSPSQEIHARFRRGARSRSLLTEVETPIRNEWGCRCNDSSDGGGGSLLTEVETPTRNGWESKSSDSSDGGGGSLLTEVETPTRSEWESRSSGSSDASGGGGSGAAAKPATTEGEFPETR
EKMSGIRRAIAKAMVHSKHTAPHVTLMDEADVTKLVAHRKKFKAIAAEKGIKLTFLPYVVKALVSALREYPVLNTAIDDETEEIIQKHYYNIGIAADTDRGLLVPVIKHADRKPIFALAQEINELAEKARDGKLTPG
EMKGASCTITNIGSAGGQWFTPVINHPEVAILGIGRIAEKPIVRDGEIVAAPMLALSLSFDHRMIDGATAQKALNHIKRLLSDPELLLMGGGGSFSEEQKKALDLAFYFDRRLTPEWRRYLSQRLGLNEEQIERWFR
RKEQQIGWSHPQFEKGSAKFVAAWTLKAAA

>M2ex3-5GS-I3-01v9a-LD7-PADRE (M2ex3-I3-01v9a-L7P)

DAMKRGLCCVLLLCGAVFVSPSQEIHARFRRGARSRSLLTEVETPIRNEWGCRCNDSSDGGGGSLLTEVETPTRNGWESKSSDSSDGGGGSLLTEVETPTRSEWESRSSGSSDASGGGGSAKLAEELQKKMEELFKK
HKIVAVLRANSVEEAKEKALAVFEGGVHLIEITFTVPDADTVIKELSFLKEKGAIIGAGTVTSVEQCRKAVESGAEFIVSPHLDAEITVFCLEKGVFYMPGVMTPTELVKAMKLGHNILKLFPGEVVGPQFVKAMKG
PFPNVKFVPTGGVNLDNVCEWFKAGVLAVGVGSALVKGTPDEVREKAKAFVEKIRGCTEGGGGSSPAVDIGDRLDELEKALEALSAEDGHDDVGQRLESLLRRWNSRRADGSAKFVAAWTLKAAA

B

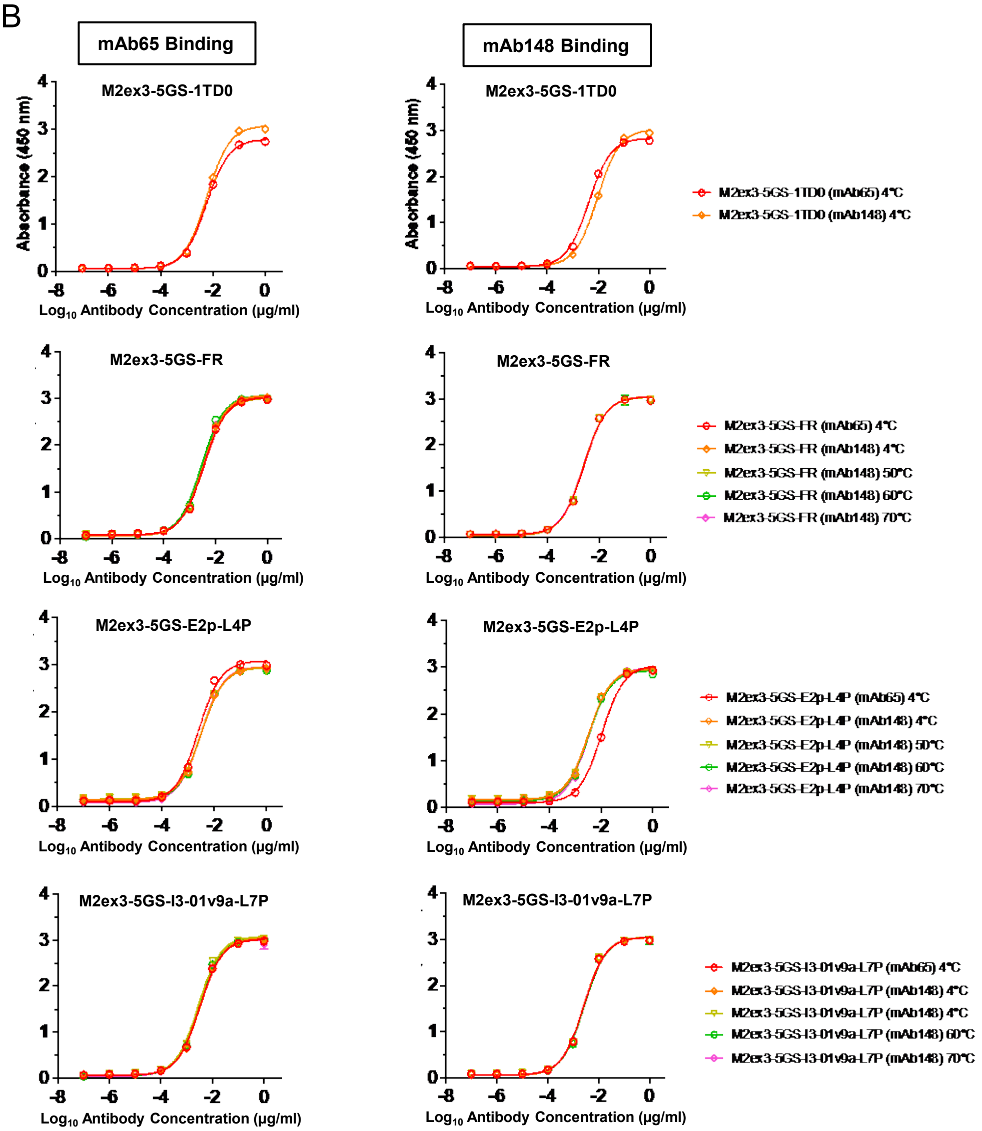


| M2ex3 | Primary Antibody Binding $EC_{50}$ (μg/ml) | |
|---|---|---|
| Groups | mAb65 | mAb148 |
| M2ex3-5GS-1TD0 4°C | 0.005851 | 0.009314 |
| M2ex3-5GS-FR 4°C | 0.003391 | 0.002564 |
| M2ex3-5GS-FR 50°C | 0.003373 | 0.00249 |
| M2ex3-5GS-FR 60°C | 0.002864 | 0.002555 |
| M2ex3-5GS-FR 70°C | 0.003176 | 0.002486 |
| M2ex3-5GS-E2p-L4P 4°C | 0.003277 | 0.003288 |
| M2ex3-5GS-E2p-L4P 50°C | 0.003149 | 0.003302 |
| M2ex3-5GS-E2p-L4P 60°C | 0.003209 | 0.003274 |
| M2ex3-5GS-E2p-L4P 70°C | 0.003013 | 0.00337 |
| M2ex3-5GS-I3-01v9a-L7P 4°C | 0.003325 | 0.002538 |
| M2ex3-5GS-I3-01v9a-L7P 50°C | 0.002818 | 0.0026 |
| M2ex3-5GS-I3-01v9a-L7P 60°C | 0.003039 | 0.002729 |
| M2ex3-5GS-I3-01v9a-L7P 70°C | 0.002895 | 0.002561 |

**Figure S4**

C

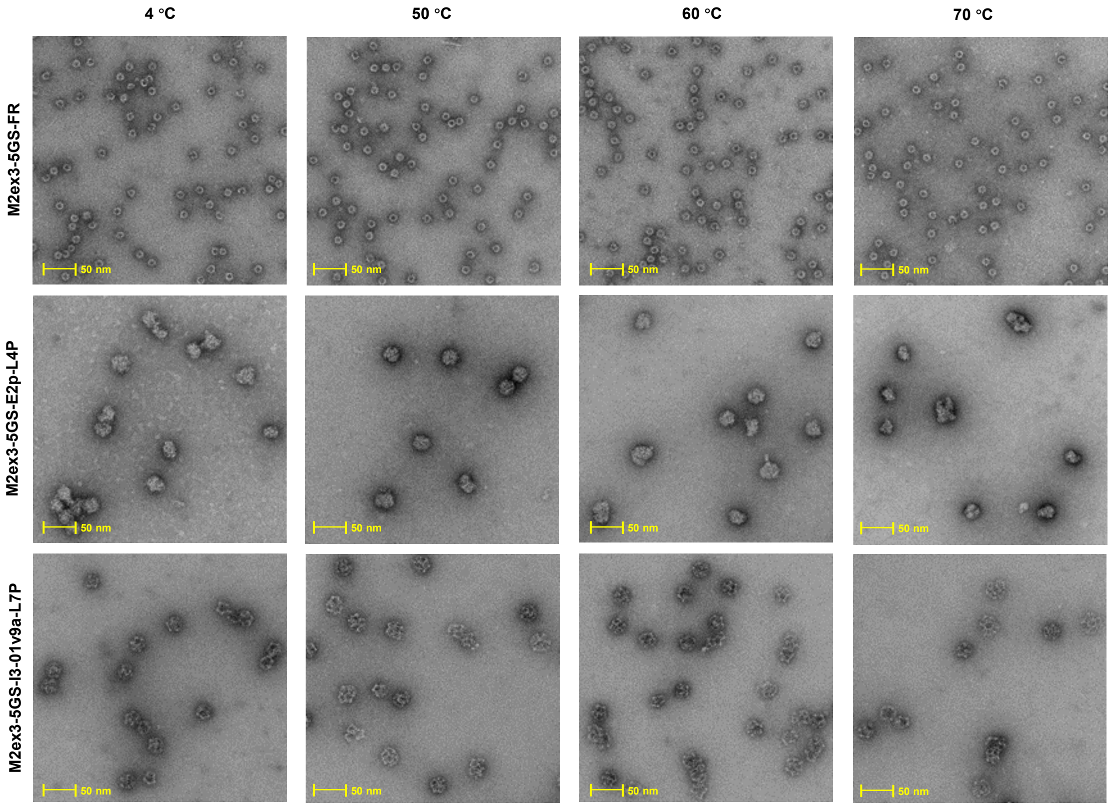


D

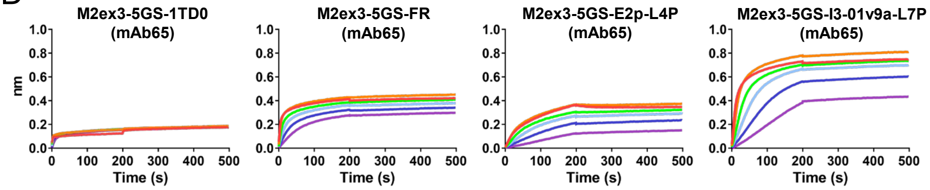


**Figure S4. Design and in vitro characterization of M2ex3-presenting SApNPs.** (**A**) Construct sequences of tandem M2e-presenting FR, E2p-L4P, and I3-01v9-L7P NPs, with the gene fragments of leader sequence, restriction site, tandem Matrix protein 2 extra-virion domain (including human, avian/swine and human/swine), flexible linker, NP-forming subunit, locking domain (LD), and PADRE highlighted in yellow, green, gray, light magenta, cyan, olive green, and red shades, respectively. (**B**) ELISA curves for M2ex3-presenting SApNPs binding to mAb148 and mAb65 antibody. (**C**) negative stain EM image for M2ex3-presenting SApNPs. (**D**) Antigenic evaluation of M2ex3-presenting SApNPs using BLI for mAb65 antibody. A two-fold concentration gradient of antigen, starting at 5 μM for M2ex3 1TD0 trimer, 80 nM for FR SApNP, and 20 nM for E2p and I3-01v9a SApNPs, was used in a titration series of six.

**Figure S5**

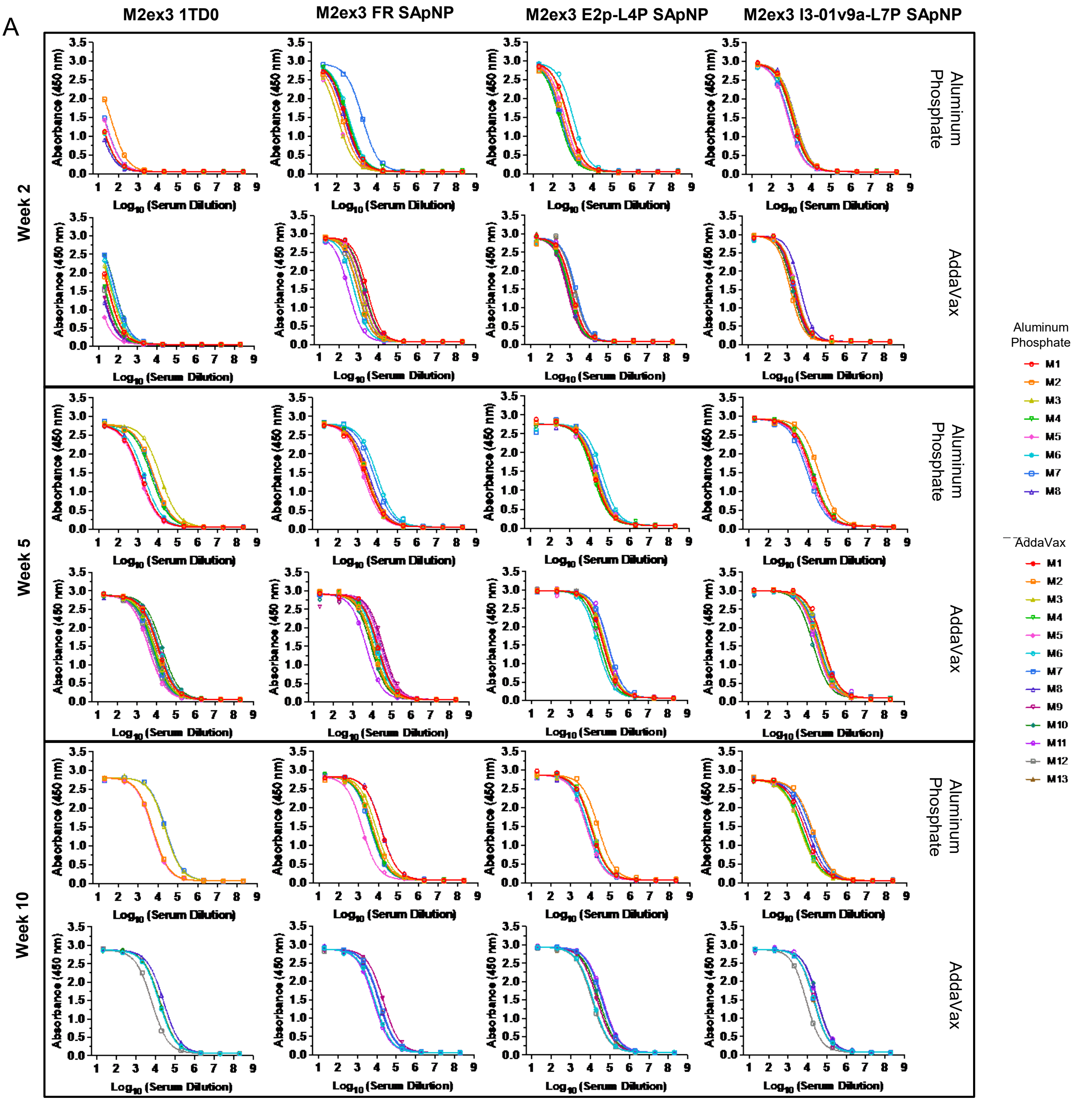
A
M2ex3 1TD0
M2ex3 FR SApNP
M2ex3 E2p-L4P SApNP
M2ex3 I3-01v9a-L7P SApNP
Week 2
Week 5
Week 10
Aluminum Phosphate
AddaVax
Absorbance (450 nm)
Log10 (Serum Dilution)
Aluminum Phosphate
M1
M2
M3
M4
M5
M6
M7
M8
AddaVax
M1
M2
M3
M4
M5
M6
M7
M8
M9
M10
M11
M12
M13

**Figure S5**

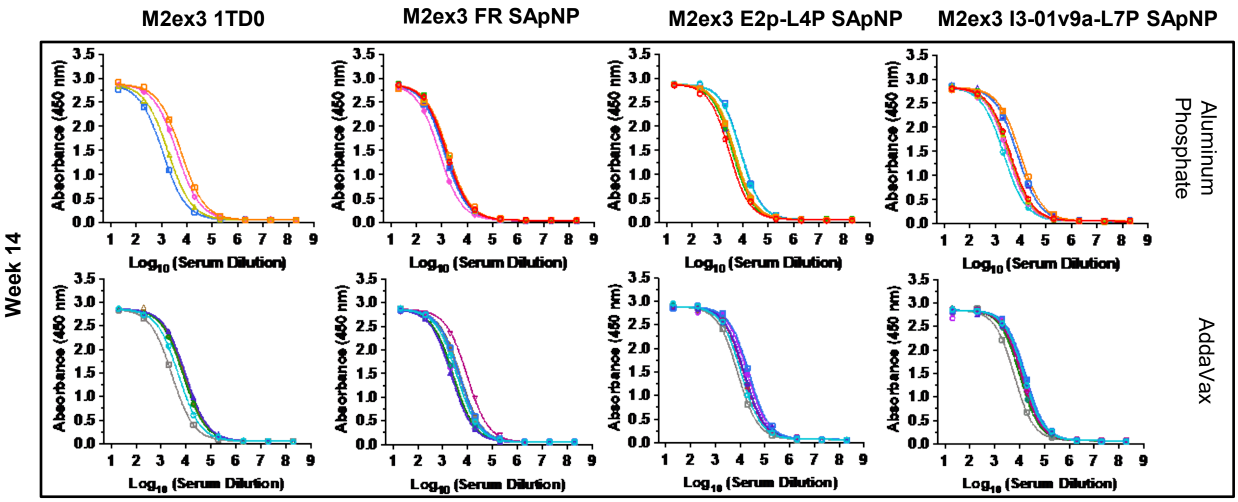


B

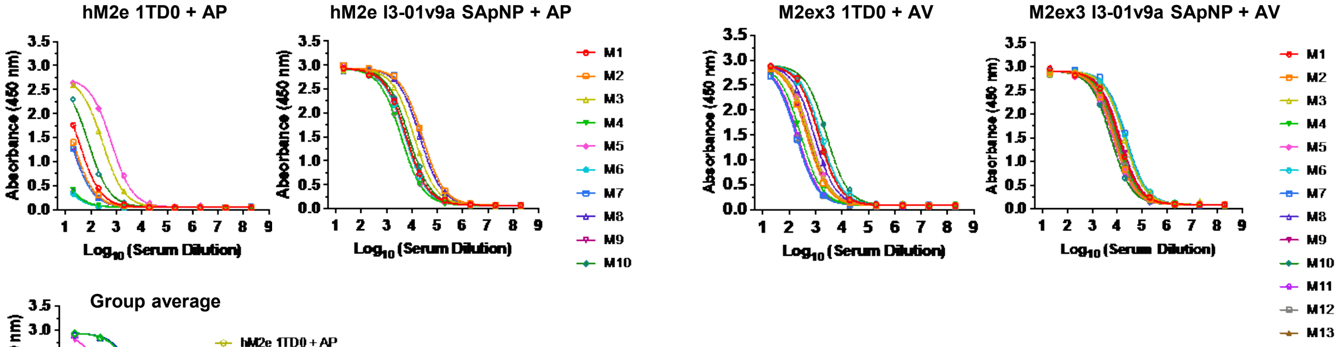


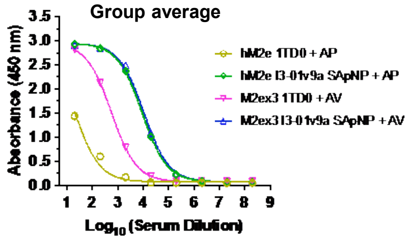


| | hM2e + AP | | M2ex3 + AV | |
|---|---|---|---|---|
| Group | 1TD0 | I3-01v9a SApNP | 1TD0 | I3-01v9a SApNP |
| hM2e coating | 99.01 | 9373 | 596.8 | 10979 |

**Figure S5. Serum binding of individual mice immunized with M2ex3 1TD0 and SApNPs.** (**A**) ELISA curves showing M2ex3-immune sera (adjuvanted with aluminum phosphate or AddaVax) binding to the M2ex3-5GS-foldon trimer probe and calculated 50% effective concentration ($EC_{50}$) values for weeks 2, 5, 10, and 14. N = 8 or 13 at weeks 2 and 5. N = variable at weeks 10 and 14 based on surviving mice/group post-H1N1 and H3N2 challenges at weeks 6 and 10, respectively. Serum was tested from mice immunized with M2ex3 1TD0 trimer, and FR, E2p, and I3-01v9a SApNP at weeks 0 and 3. The assay was performed in duplicate with a starting serum dilution of 20× followed by seven 10-fold titrations. (**B**) ELISA curves showing hM2e (+ AP)- and M2ex3 (+ AV)-immune sera binding to the hM2e-5GS-foldon trimer probe at week 5. N = 10 for hM2e groups. N = 13 for M2ex3 groups. Results indicate that M2ex3-immune sera demonstrates similar or higher binding to hM2e foldon compared to hM2e-immune sera.

#### Figure S6

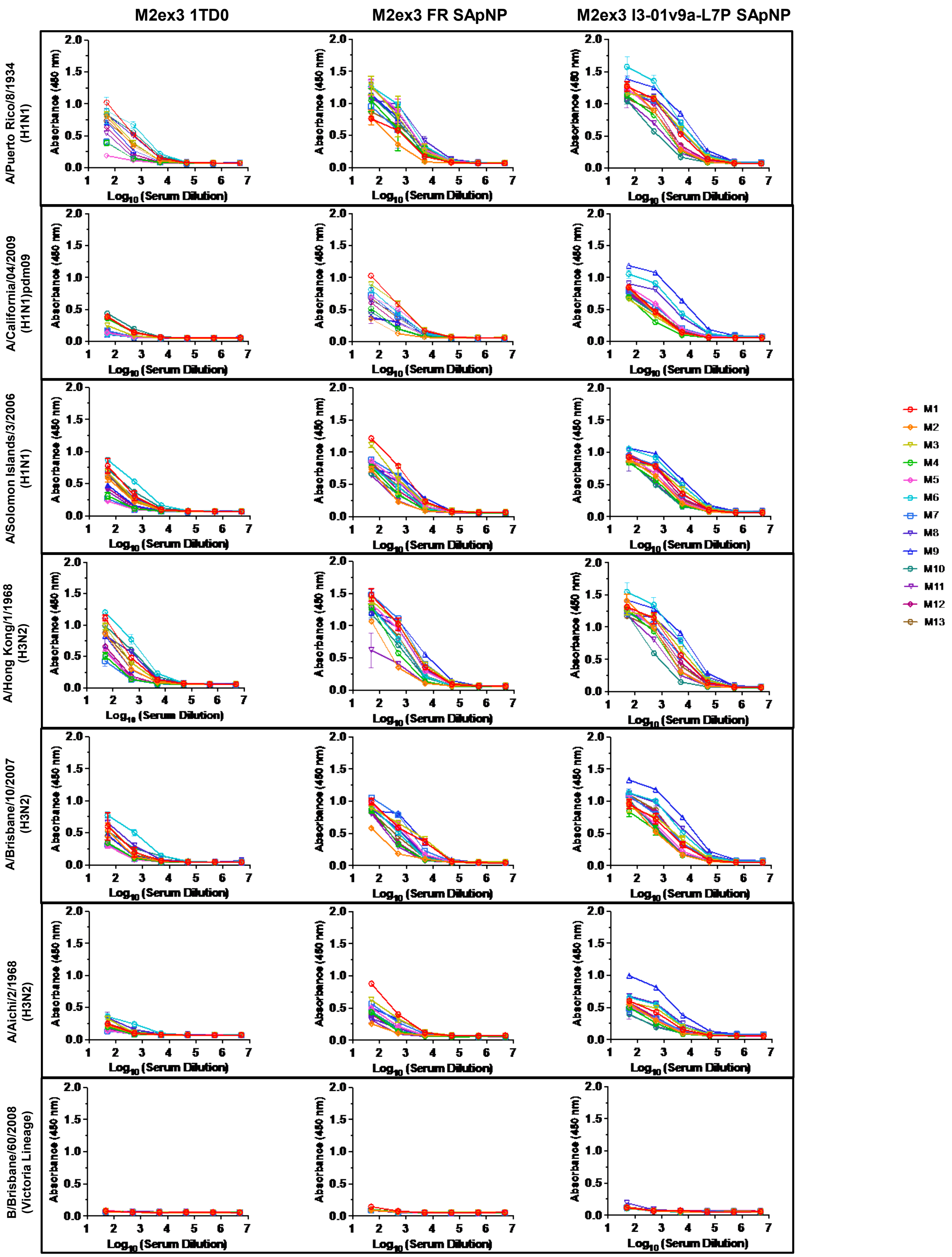

M2ex3 1TD0
M2ex3 FR SApNP
M2ex3 I3-01v9a-L7P SApNP
A/Puerto Rico/8/1934 (H1N1)
A/California/04/2009 (H1N1)pdm09
A/Solomon Islands/3/2006 (H1N1)
A/Hong Kong/1/1968 (H3N2)
A/Brisbane/10/2007 (H3N2)
A/Aichi/2/1968 (H3N2)
B/Brisbane/60/2008 (Victoria Lineage)
Absorbance (450 nm)
Log10 (Serum Dilution)
M1
M2
M3
M4
M5
M6
M7
M8
M9
M10
M11
M12
M13

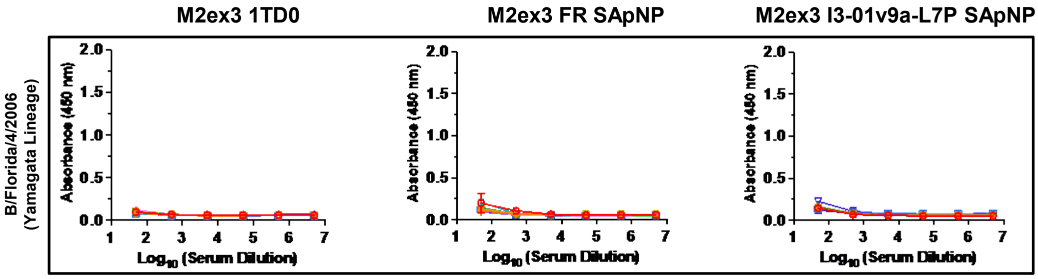


**Figure S6. M2ex3 serum binding of individual mice to cell-surface homotetrameric M2e**. Week 5 serum from mice immunized with M2ex3 1TD0, FR SApNP, and I3-01v9a SApNPs adjuvanted with AddaVax (n = 13) were tested against homotetrameric M2e on various influenza strains: A/Puerto Rico/8/1934 (H1N1), A/California/04/2009 (H1N1)pdm09, A Solomon Islands/2/2006 (H1N1), A/Hong Kong/1/1968 (H3N2), A/Brisbane/10/2007 (H3N2), A/Aichi/2/1968 (H3N2), B/Brisbane/60/2008 (Flu B, Victoria Lineage), and B/Florida/4/2006 (Flu B, Yamagata Lineage). MAb148 (M2e antibody) was used as a IAV positive control. The assay was performed in duplicate with a starting serum dilution of 50× followed by five 10-fold titrations.

**Figure S7**

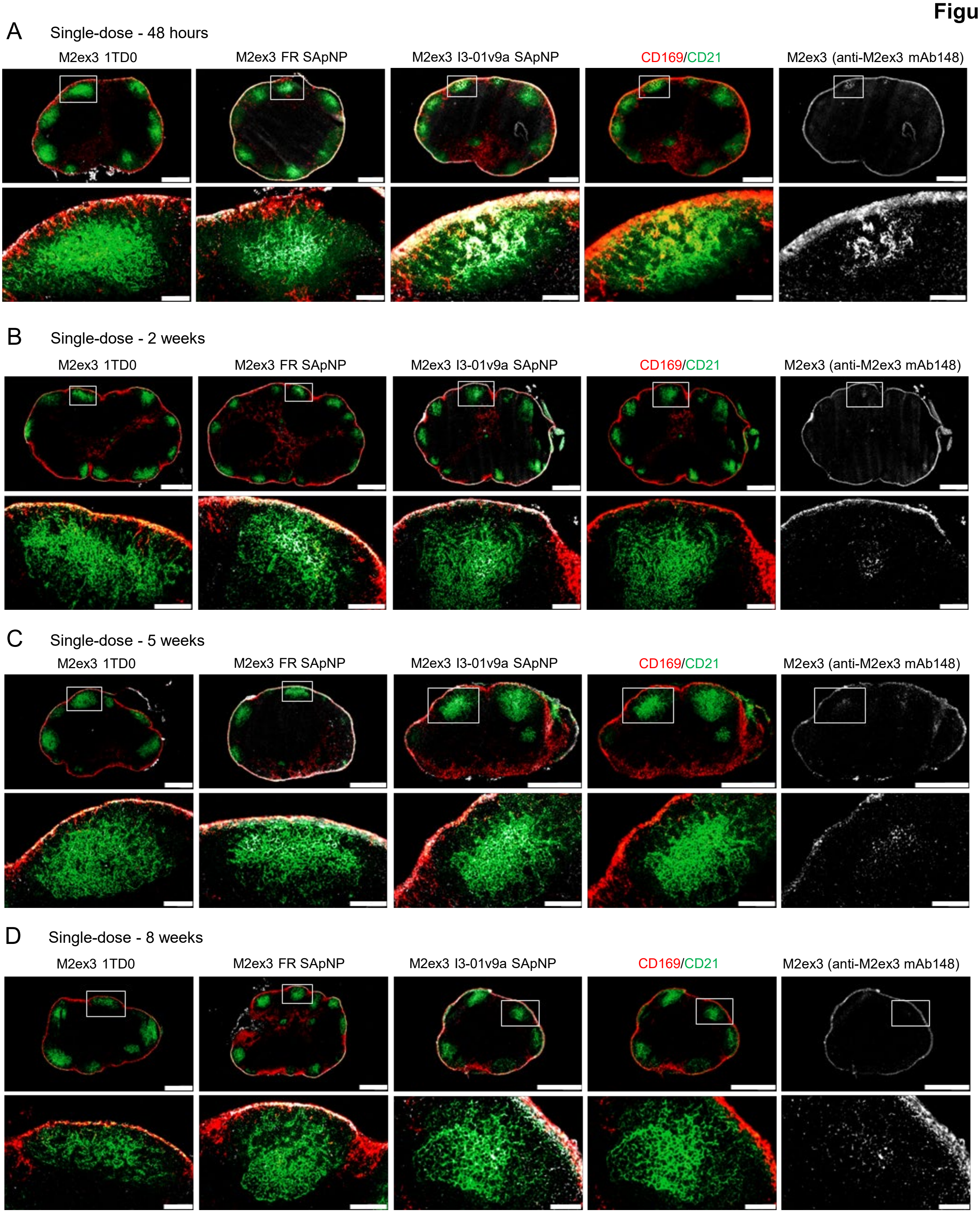


**Figure S7. Immunohistological images of M2ex3 1TD0 and SApNPs in lymph nodes.** Immunostaining images of M2ex3 1TD0 and M2ex3-presenting FR and I3-01v9a SApNP interaction with FDC networks in lymph node follicles at (**A**) 48 hours, (**B**) 2 weeks, (**C**) 5 weeks, and (**D**) 8 weeks after a single-dose injection (10 μg per injection, totaling 40 μg per mouse). Immunofluorescent images are pseudo color coded ($CD21^+$, green; $CD169^+$, red; anti-M2ex3 mAb148, white). Scale bars = 500 and 100 μm for complete lymph node and enlarged image of a follicle, respectively. Data were collected from more than 10 lymph node follicles (n = 3-5 mice/group).

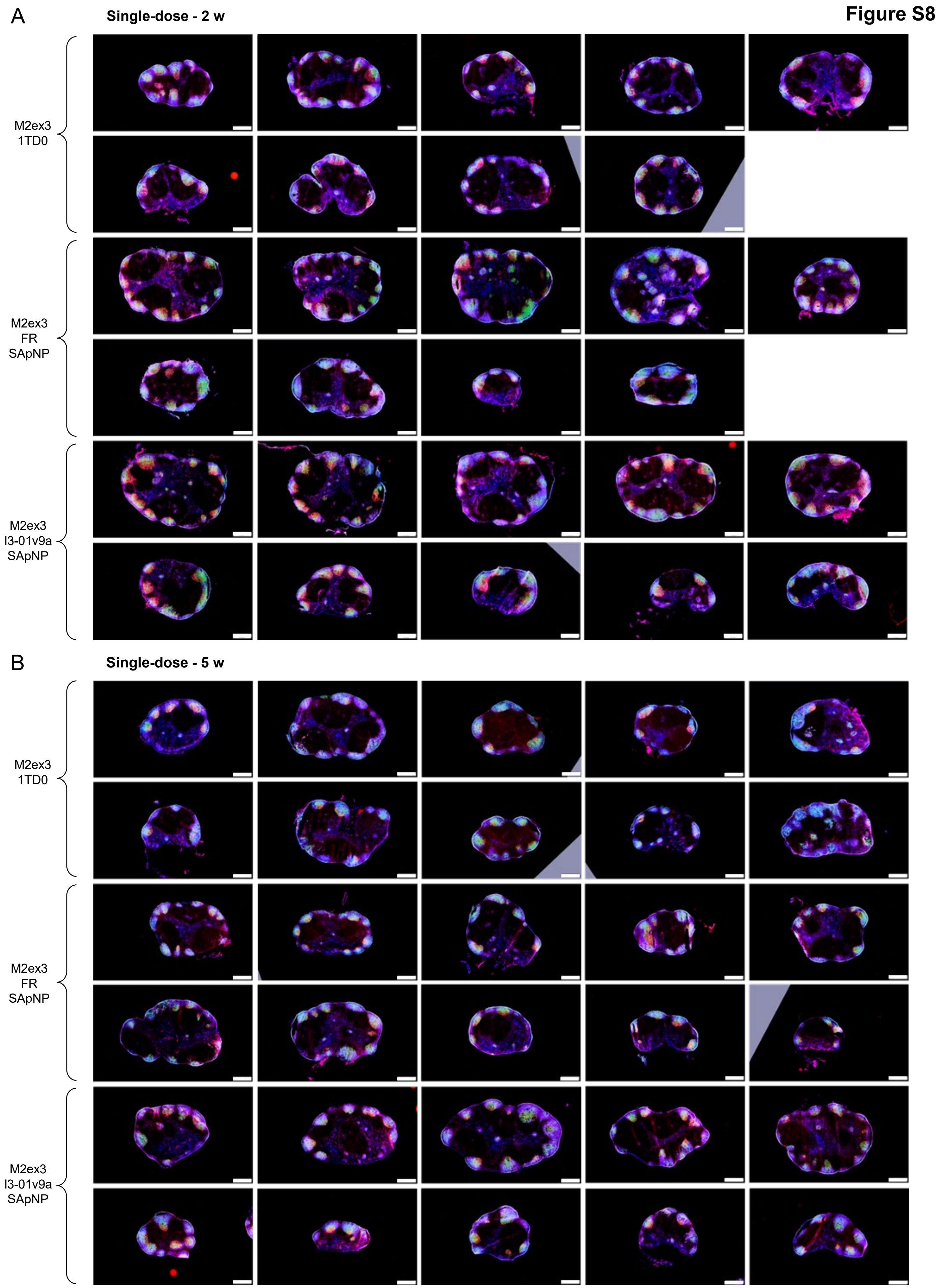
Figure S8
A
Single-dose - 2 w
M2ex3
1TD0
M2ex3
FR
SApNP
M2ex3
I3-01v9a
SApNP
B
Single-dose - 5 w
M2ex3
1TD0
M2ex3
FR
SApNP
M2ex3
I3-01v9a
SApNP

C Single-dose - 8 w

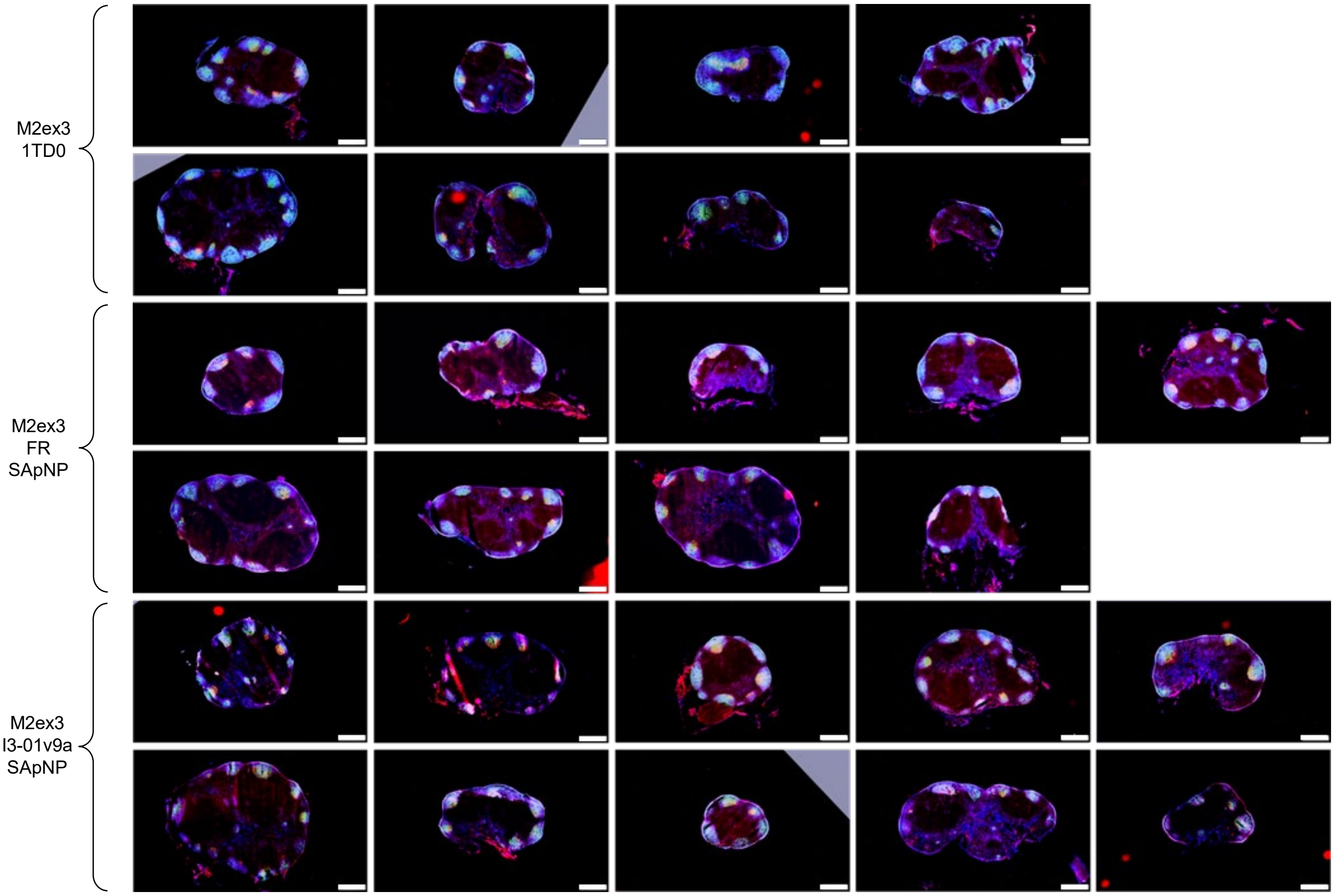


D Prime-boost - 3 w + 2 w

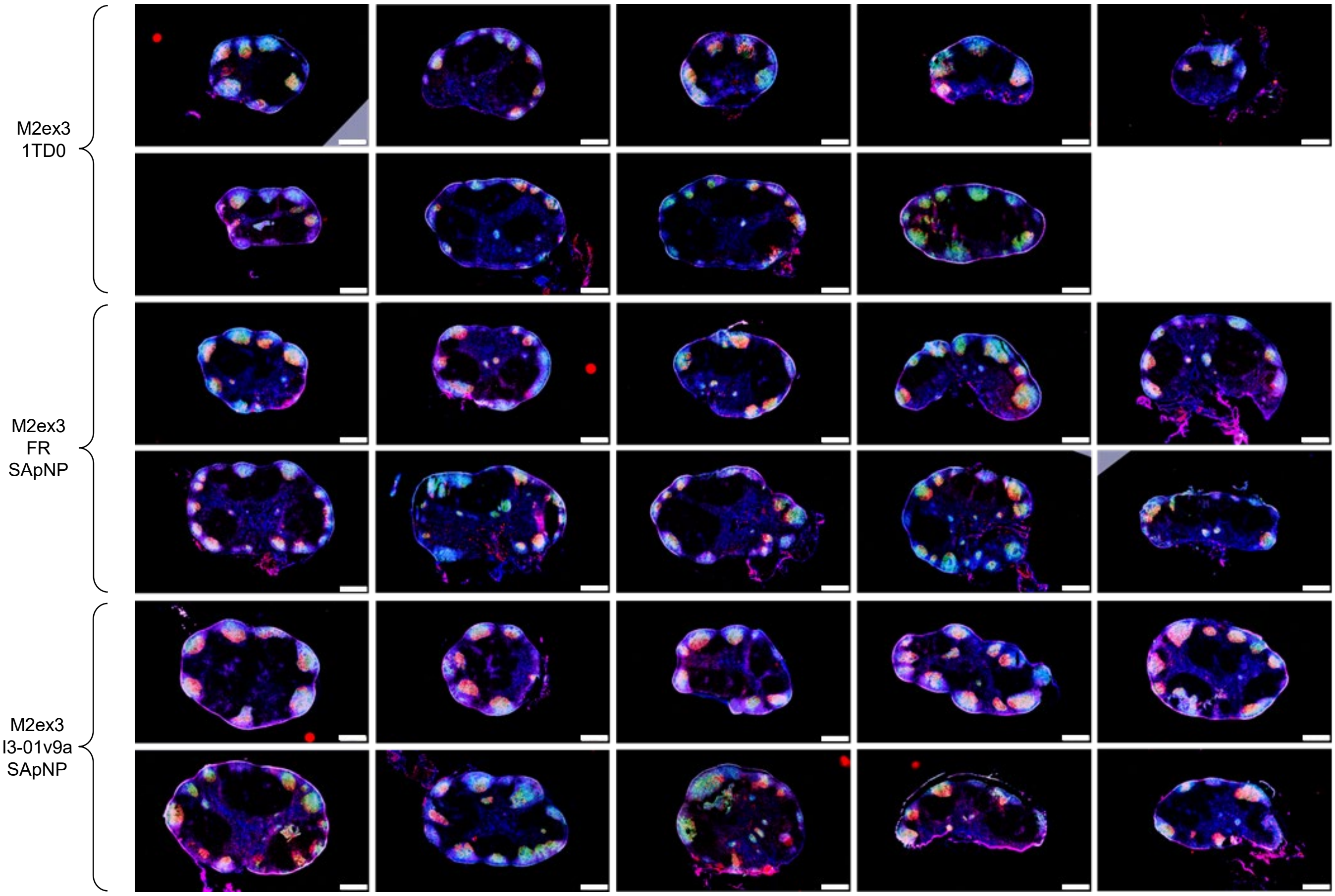

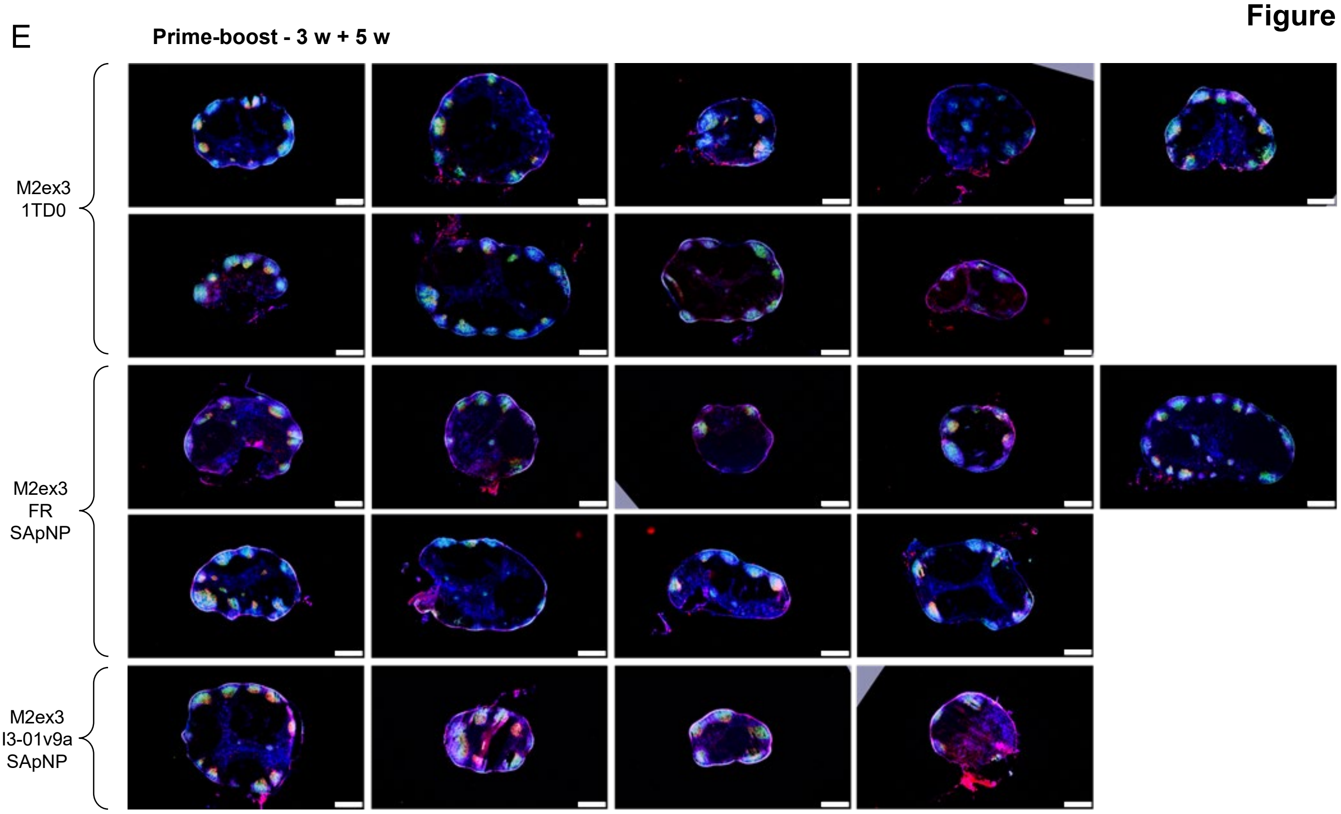


**Figure S8. Immunohistological analysis of M2ex3 1TD0 and SApNP vaccine-induced GCs.** Immunohistological images of GCs at (**A**) 2, (**B**) 5, and (**C**) 8 weeks after a single-dose injection of M2ex3 1TD0 and M2ex3-presenting FR, and I3-01v9a SApNP vaccines (10 μg per injection, totaling 40 μg per mouse), with a scale bar of 500 μm for each image. Images of GCs at (**D**) 2 and (**E**) 5 weeks after the boost, which occurred at 3 weeks after the first dose (n = 5 mice/group).

**Figure S9**

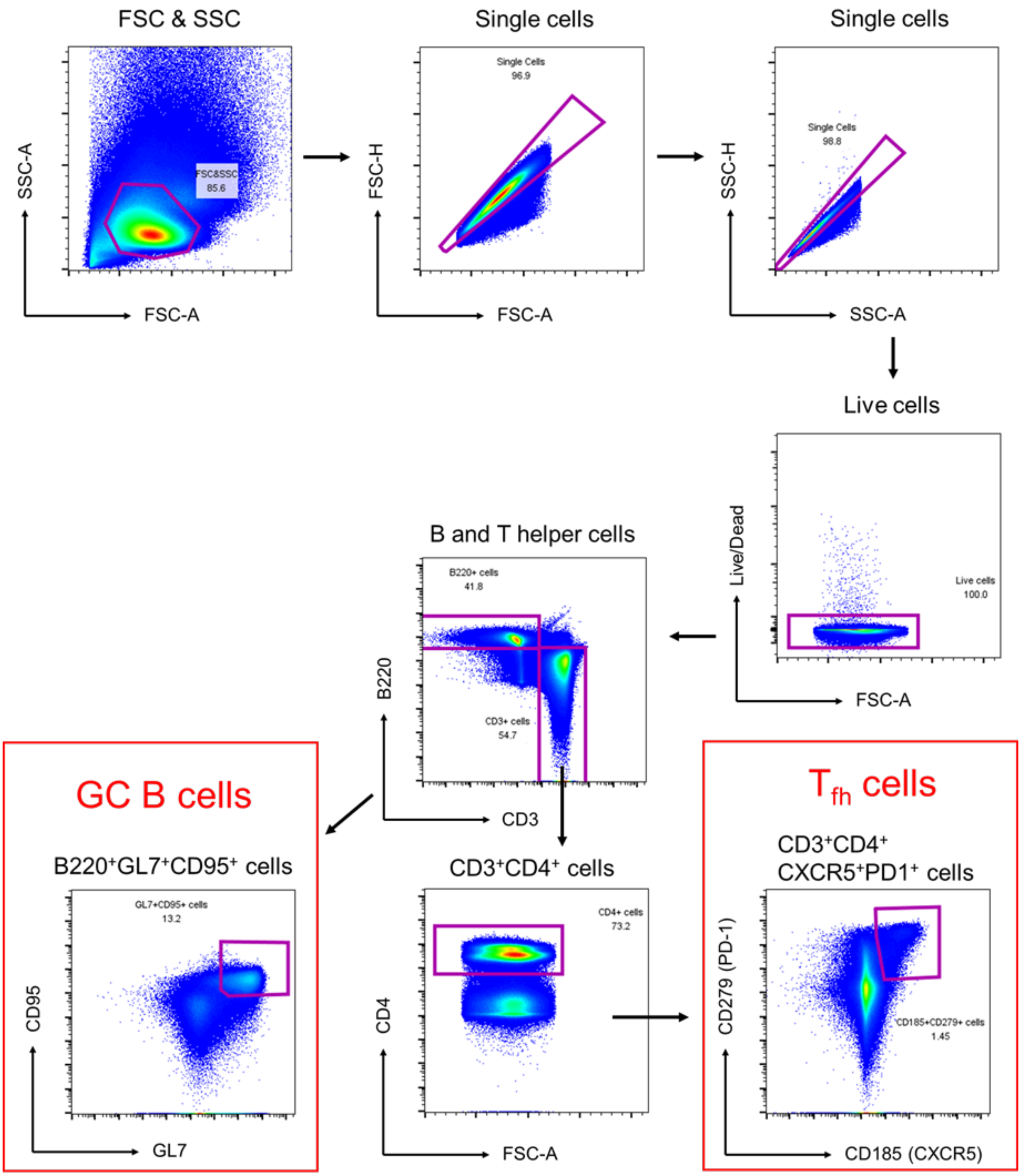


**Figure S9. Flow cytometry analysis of M2ex3 1TD0 and SApNP vaccine-induced GCs.** Gating strategy for analyzing GCs (GC B cells and T follicular helper cells) using flow cytometry (n = 5 mice/group).

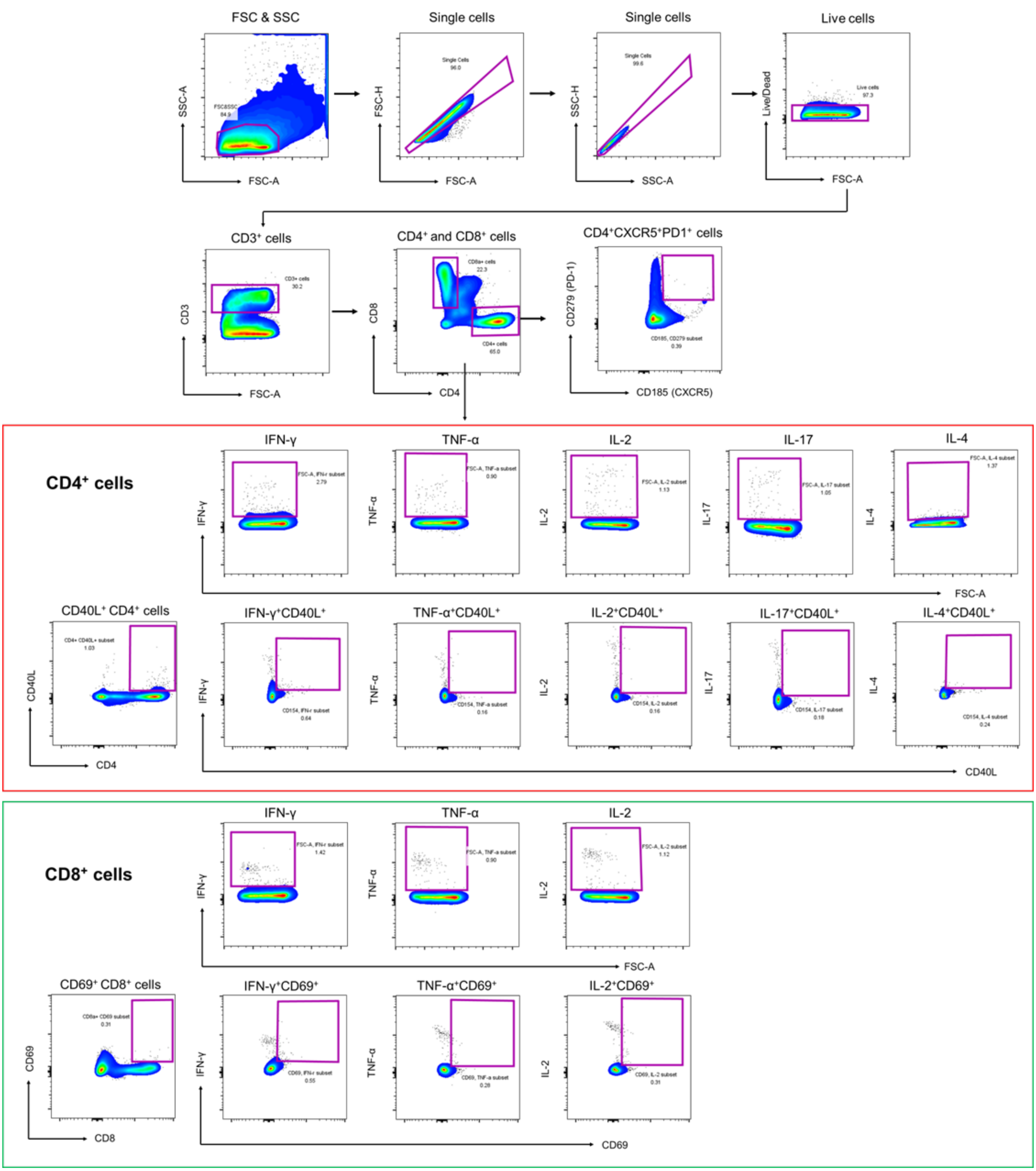


**Figure S10. Flow cytometry analysis of M2ex3 1TD0 and SApNP vaccine-induced T cell responses.** Gating strategy for analyzing CD4⁺ and CD8⁺ T cell responses against M2ex3 1TD0 using flow cytometry at 5 days after prime-boost immunization and virus challenge (n = 5 mice/group).

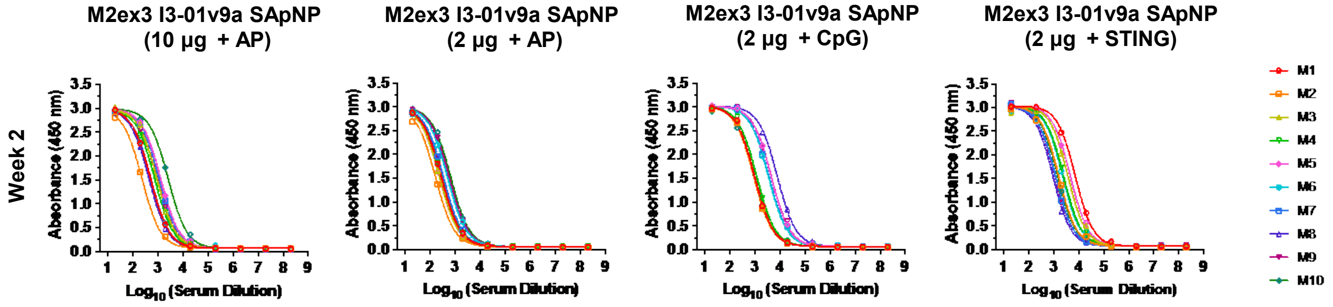


**Figure S11. Serum binding of individual mice immunized with SApNPs and various adjuvants.** ELISA curves showing M2ex-5GS-foldon trimer probe binding of sera from mice immunized with a single-dose of M2ex3 I3-01v9a SApNP + aluminum phosphate, CpG, or STING agonist at week 2 (n = 10). The assay was performed in duplicate starting at a serum dilution of 20× followed by seven 10-fold titrations.
